# Neuromodulation of Behavioral Specialization: Tachykinin Signaling Inhibits Task-specific Behavioral Responsiveness in Honeybee Workers

**DOI:** 10.1101/2021.01.12.426394

**Authors:** Bin Han, Qiaohong Wei, Fan Wu, Han Hu, Chuan Ma, Lifeng Meng, Xufeng Zhang, Mao Feng, Yu Fang, Olav Rueppell, Jianke Li

## Abstract

Behavioral specialization is key to the success of social insects and often compartmentalized among colony members leading to division of labor. Response thresholds to task-specific stimuli proximally regulate behavioral specialization but their neurobiological regulation is not understood. Here, we show that response thresholds to task-relevant stimuli correspond to the specialization of three behavioral phenotypes of honeybee workers. Quantitative neuropeptidome comparisons suggest two tachykinin-related peptides (TRP2 and TRP3) as candidates for the modification of these response thresholds. Based on our characterization of their receptor binding and downstream signaling, we then confirm the functional role of tachykinins: TRP2 injection and RNAi cause consistent, opposite effects on responsiveness to task-specific stimuli of each behaviorally specialized phenotype but not to stimuli that are unrelated to their tasks. Thus, our study demonstrates that TRP-signaling regulates the degree of task-specific responsiveness of specialized honeybee workers and may control the context-specificity of behavior in animals more generally.

## 1. Introduction

Behavioral responses of animals to external and internal stimuli have evolved to optimize survival and reproduction under average circumstances [1]. However, environmental and inter-individual variability commonly cause deviations from the average, resulting in selection for context-specific and condition-dependent behavior [2–4]. Evolutionary constraint [5] of behavior occurs in form of behavioral syndromes, differences among individuals that manifest across different contexts [6]. Advantages of behavioral plasticity and specificity have been documented in many systems and some neuroendocrine mechanisms have been identified [7, 8]. However, general neural mechanisms that allow the sophistication of behavioral repertoires by increasing context-specificity of behavioral responses remain insufficiently understood.

Behavioral modulation is particularly important in social species in which social interactions provide a high diversity of behavioral context [9, 10]. However, social evolution also allows individuals to restrict their behavioral repertoires through temporal or permanent behavioral specialization [11]. This specialization and the resulting division of labor are believed to be major contributors to the successful colony life of many social insects despite its potential costs [12]. Advanced social evolution thus allows inter-individual plasticity to replace individual behavioral plasticity and decoupling of behavioral responses may be more efficient across different individuals than within solitary individuals. Nevertheless, the principal problem of behavioral plasticity across different contexts remains the same, and social insects can be constrained in their behavioral evolution by correlated selection responses across different behaviors or castes [13, 14].

Behavior often occurs in response to a specific stimulus exceeding an individual’s specific response threshold [15, 16]. Response thresholds depend on internal physiological states [17], particularly the concentration of neurotransmitters and neuromodulators in the central nervous system [18, 19]. Response thresholds translate the value of perceived stimuli into probabilities of behavioral responses and vary among individuals [20]. In social insects, individual variation in response thresholds is linked to division of labor [21–23] and numerous studies have characterized this link across multiple levels of biological organization [20, 24, 25]. Many aspects of the division of labor in the social model *Apis mellifera* are driven by a life-long behavioral ontogeny, leading to age-polyethism [26]. Young bees perform numerous inside tasks, most prominently brood care in form of alloparental nursing behavior, before transitioning to a mix of other in-hive tasks [27]. Similar to the highly-specialized nursing stage, the final behavioral state of older bees as outside foragers is almost exclusive of other tasks [26]. Moreover, foragers often specialize on collecting only one of the principal food sources, pollen or nectar [28]. These behavioral specialists (nurses, nectar foragers, and pollen foragers) exhibit pronounced differences in their responsiveness to task-related stimuli. Responsiveness to brood pheromones peaks at typical nursing age [29]. In contrast, foragers have a lower response threshold to sugars and light than nurses [30, 31]. Among foragers, pollen specialists exhibit higher responsiveness to sucrose and pollen stimuli than nectar foragers [32, 33]. Response thresholds can be quantified based on the honeybees’ reflexive extension of their proboscis in response to stimuli, such as sucrose [20]. The spontaneous proboscis extension reflex (PER) to sucrose has been expanded to other stimuli that bees spontaneously respond to [34, 35] and conditioned stimuli to which no spontaneous responses occur [36].

Response thresholds can be modified by biogenic amines, and dopamine, 5-hydroxy-tryptamine, octopamine, and tyramine have been implicated in the regulation of different behaviors of worker bees [37]. However, neuropeptides have not been studied although they are a diverse group of neurotransmitters that can also act as neurohormones on distal targets to coordinate a wide range of internal states and behavioral processes [38]. Neuropeptides are intimately involved in food perception and social interaction of insects [39], two processes that are central to division of labor in social insects [40]. Neuropeptides mediate pheromonal effects on physiology [41, 42] and usually exhibit a high degree of specificity [43, 44]. Therefore, neuropeptides are prime candidates for mediating the independent adjustment of socially relevant response thresholds that mediate honeybee workers specialization and division of labor.

More than 100 mature neuropeptides derived from 22 protein precursors have been identified in the Western honeybee, *Apis mellifera* [45, 46]. Several neuropeptides, including allatostatin and tachykinin-related peptides (TRPs), may be involved in the control of social behavior of honeybees, such as aggression [47], foraging [48], brood care [45], and possibly a wide array of other behaviors [49]. However, these results are based on correlations between behavior and neuropeptide expression and more detailed studies are needed to understand the causal roles of neuropeptides in the behavioral specialization among honeybee workers. Here, we report the results of a comprehensive study to test the hypothesis that neuropeptides regulate the division of labor in honeybees. We initially compared response thresholds to task-relevant stimuli among behaviorally-defined worker groups of two honeybee species.

These response thresholds were correlated with neuropeptide expression levels, especially TRPs, suggesting a role of TRPs in worker specialization. Based on these results, we characterized the TRP signaling pathway molecularly. Finally, we demonstrated in a series of TRP injections and RNAi-mediated knockdown of the *TRP* and its receptor *TRPR* a causal role of this pathway in modulating different response thresholds in a task-specific manner.

## 2. Results

### 2.1 The task-specific responsiveness of worker bees shows significant variations between behavioral phenotypes and the two honeybee species

In our comparisons of the PER of worker bees to task-specific stimuli, including sucrose solution, pollen, and larva, significant differences were found between behavioral phenotypes and the two honeybee species (Fig. 1A, Table S1 and S2).

**Fig. 1:**
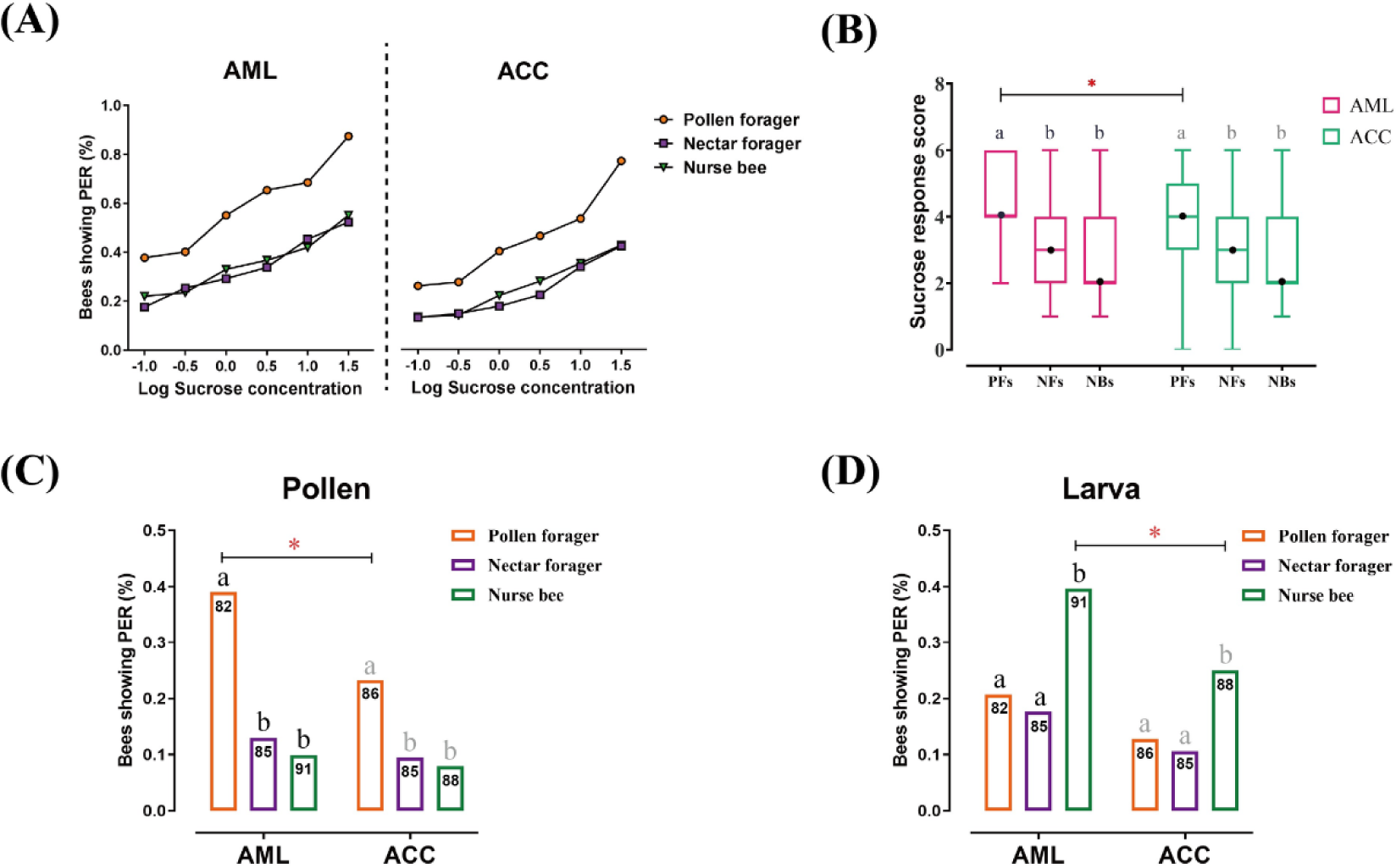
Responses to sucrose solution, pollen, and larva stimulations are significant different among behavioral phenotypes and between honeybee species. (A) The proportion of pollen foragers (PFs), nectar foragers (NFs), and nurse bees (NBs) showing a proboscis extension reflex (PER) increased with increasing concentrations of sucrose solutions. Left: *Apis mellifera ligustica* (AML), right: *Apis cerana cerana* (ACC). Details of the statistical results of our comparisons of sucrose responsiveness between behavioral phenotypes and bee species are listed in Table S2. (B) Median sucrose response scores (SRS; intermediate lines) and quartiles (upper and lower lines) of PFs, NFs, and NBs. Kruskal-Wallis tests with Bonferroni correction were used to compare the SRSs of the three behavioral phenotypes in the same species and significant differences are denoted by letters at p < 0.05. Pairwise Mann-Whitney U tests were used for comparing the same phenotype between two honeybee species (* denotes p < 0.05). (C) Proportion of PFs, NFs, and NBs showing PER to pollen stimulation of their antennae. (D) Proportion of PFs, NFs, and NBs showing PER to antennal stimulation with larvae. Numbers in bars represent the number of individuals sampled in each group. Independent Chi-square tests were used to compare the responsiveness to pollen or larvae between species (* denotes p < 0.05) and among behavioral phenotypes within species (letters indicate significant difference at p < 0.05).

The percentage of bees showing a PER increased with sucrose concentration across all experimental groups (Fig. 1A). In both, AML and ACC, the sucrose response scores (SRSs) of PFs were higher than the SRSs of NFs (AML: Z = 7.0, *p =* <0.001; ACC: Z = 6.1, *p* < 0.001) and NBs (AML: Z = 5.9, *p* < 0.001; ACC: Z = 5.2, *p* < 0.001), while no significant difference between NFs and NBs was observed in either species. PFs were more responsive than NFs and NBs to all sucrose concentrations. The species comparison between AML and ACC showed significant higher sucrose responsiveness in PFs of AML than in PFs of ACC (Z = 2.361, *p* = 0.018), specifically at sucrose concentrations of 0.3% (*χ*^2^ = 4.1, *p* = 0.042), 1.0% (*χ*^2^ = 5.2, *p* = 0.001), 3.0% (*χ*^2^ = 8.4, *p* = 0.023), and 10.0% (*χ*^2^ = 5.3, *p* = 0.021). Nectar foragers of AML and ACC showed no significant difference in overall SRS, but NFs of AML were more responsive than NFs of ACC at sucrose concentrations of 0.3% (*χ*^2^ = 4.5, *p* = 0.035), 1.0% (*χ*^2^ = 4.5, *p* = 0.033), and 3.0% (*χ*^2^ = 4.0, *p* = 0.046). There was no significant difference between NBs of AML and ACC in sucrose responsiveness.

In AML, PFs were more responsive to pollen stimulation than NFs (*χ*^2^ = 14.9, *p* = 0.002) and NBs (*χ*^2^ = 20.2, *p* < 0.001), while there were no significantly statistical differences between NFs and NBs. Likewise, PFs of ACC were more sensitive than NFs (*χ*^2^ = 6.0, *p* = 0.015) and NBs (*χ*^2^ = 7.8, *p* = 0.001) without a statistically significant difference between NFs and NBs. Pollen foragers of AML showed a significant higher pollen responsiveness than of ACC (*χ*^2^ = 4.9, *p* = 0.031), with no significant species differences in NFs and NBs (Fig. 1B).

In larva responsiveness assay, NBs of AML showed increased responsiveness to larva stimulation compared to PFs (*χ*^2^ = 7.2, *p* = 0.006) and NFs (*χ*^2^ = 10.3, *p* = 0.001). Likewise, NBs of ACC were more sensitive than PFs (*χ*^2^ = 4.2, *p* = 0.013) and NFs (*χ*^2^ = 6.1, *p* = 0.002). Nurse bees of AML were significantly more sensitive to larvae (*χ*^2^ = 4.3, *p* = 0.027) than NBs of ACC, with no significant species differences in PFs and NFs. (Fig. 1C).

### 2.2 Quantitative peptidomics reveal brain neuropeptide signatures of behavior

Our LC-MS/MS-based comparisons of the brain neuropeptidomes of NBs, PFs, and NFs of AML and ACC revealed numerous differences among experimental groups but only two tachykinins showed consistent patterns relating to the task-specific responsiveness of the experimental groups. Overall, 132 unique neuropeptides derived from 23 neuropeptide families were identified in the brain of AML worker bees (Table S3). In the brain of ACC worker bees, for the first time, 116 unique neuropeptides derived from 22 neuropeptide families were identified (Table S4).

Quantitative comparison among the three behavioral phenotypes of AML showed that 40 neuropeptides derived from 16 neuropeptide families were differentially expressed the brain (Fig. 2, Table S5). Among 19 differential expressed neuropeptides between PFs and NFs, 9 neuropeptides were upregulated in PFs and 10 were upregulated in NFs. Among 24 differential expressed neuropeptides between PFs and NBs, 18 were upregulated in PFs and 6 were upregulated in NBs. Moreover, 21 differential expressed neuropeptides were found between NFs and NBs, with 14 upregulated in PFs and 7 upregulated in NBs.

**Fig. 2:**
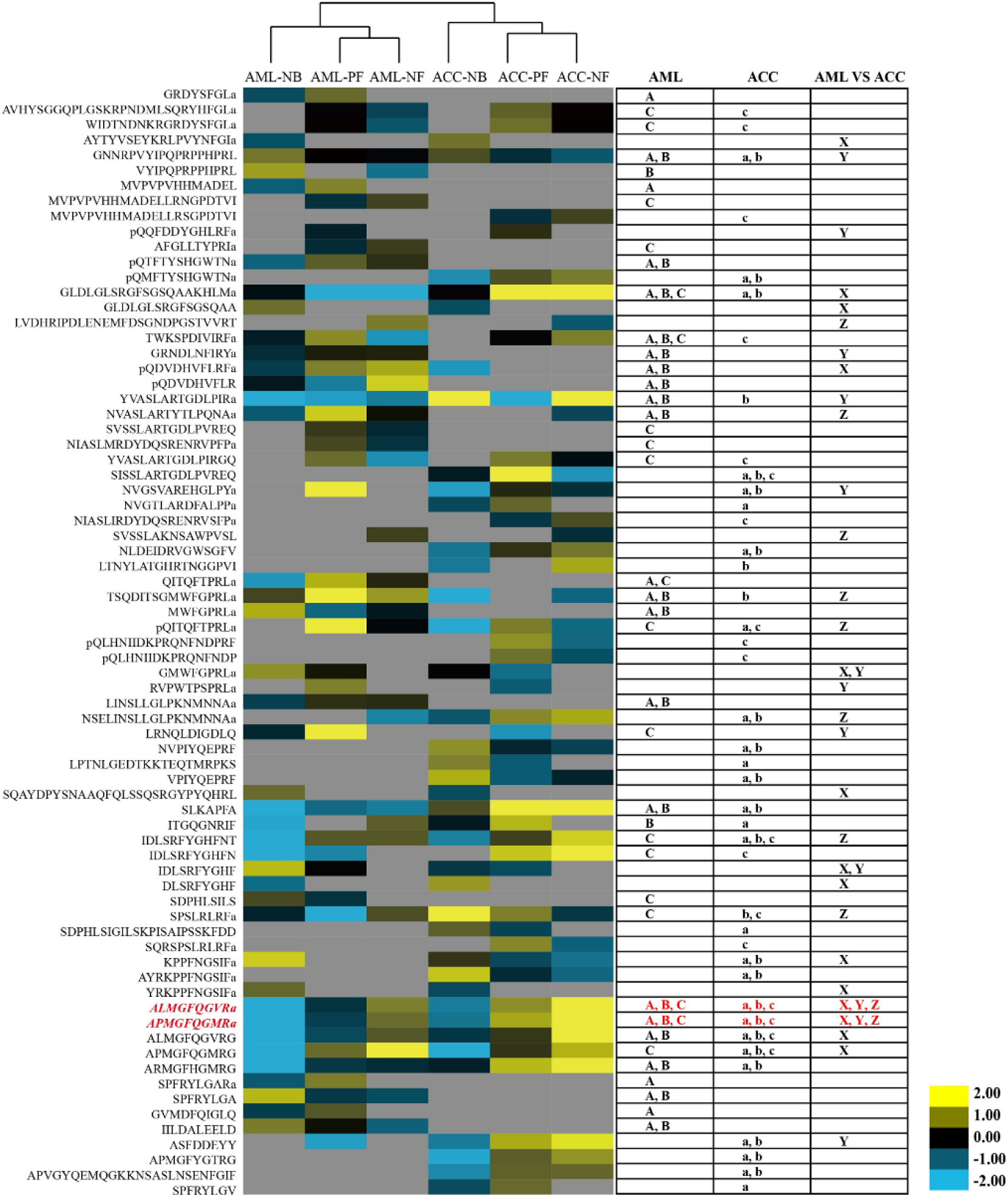
Quantitative comparison of the brain neuropeptides. The brain neuropeptides were quantitatively compared between nurse bees (NBs), pollen foragers (PFs), and nectar foragers (NFs) of *Apis* mellifera *ligustica* (AML) and *Apis cerana cerana* (ACC). The up- and down-regulated peptides are indicated by yellow and blue colors, respectively. Color intensity indicates the relative expressional level, as noted in the key. Letters A, B, and C on the right represent significant differences between NBs and PFs, NBs and NFs, and PFs and NFs in AML, respectively; a, b, and c represent significant differences between NBs and PFs, NBs and NFs, and PFs and NFs in ACC, respectively; X, Y, and Z represent significant differences of NBs, PFs, and NFs between AML and ACC, respectively. For detailed quantification data, see Table S5 S6, and S7.

In ACC 18 neuropeptides were differentially expressed between PFs and NFs, with 9 upregulated in each group. Between PFs and NBs, 27 neuropeptides showed different expression levels: 20 were upregulated in PFs and 7 were upregulated in NBs (Table S6). Twenty-five neuropeptides were differentially expressed between NFs and NBs, with 19 upregulated in NFs and 6 in NBs. The species comparison between AML and ACC, the number of differentially expressed neuropeptides in NBs, PFs and NFs was 13, 10, and 11, of which 7, 6, and 6 were upregulated in AML respectively (Table S7).

### 2.3 TRP/TRPR signaling couples to G_αq_ and G_αs_ pathways and triggers the ERK cascade

A series of cellular and molecular experiments confirmed that honeybee TRPR was expressed in the cell membrane and specifically activated by TRP, triggering intracellular cAMP accumulation, Ca^2+^ mobilization, and ERK phosphorylation by dually coupling G_αs_ and G_αq_ signaling pathways.

The honeybee *TRPR* gene was successfully cloned and expressed in the human embryonic kidney cells (HEK293) and the insect *Spodoptera frugiperda* pupal ovary cells (Sf21). Significant cell surface expression was observed by fluorescence microscopy (Fig. 3A and 3B), revealing that the honeybee TRPR was exclusively localized in the cell membrane in HEK293 and Sf21 cells.

**Fig. 3:**
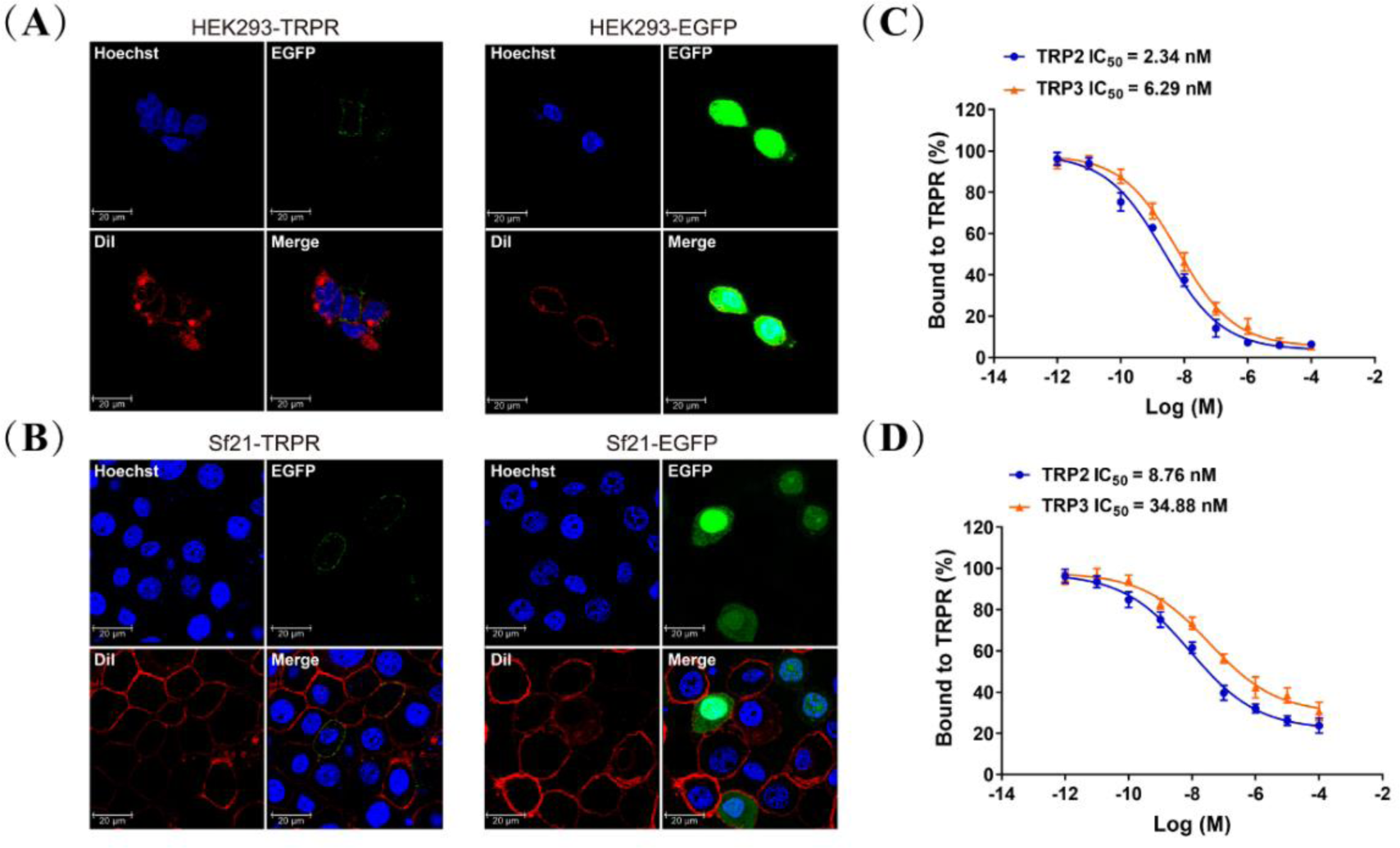
Expression of TRPR and direct interaction of TRPs with TRPR in cell culture. (A) and (B) HEK293 and Sf21 cells expressing TRPR-EGFP and EGFP (green) were stained with a membrane plasma probe DiI (red) and a nuclei probe Hoechst (blue), and assessed by confocal microscopy. (C) and (D) Competitive inhibition of TAMRA-TRP2 and TAMRA-TRP3 binding to TRPR in HEK293 and Sf21 cells, and all data are presented as mean ± s.e.m. from three independent experiments.

Competitive binding assays with labeled TRP2 and TRP3 confirmed high affinity of the TRPR for both. The observed IC_50_ values for TRP2 and TRP3 were 2.34 nM and 6.29 nM in HEK293 cells and 8.76 nM and 34.88 nM in Sf21 cells, respectively (Fig. 3C and 3D). These competition binding analyses strongly suggested a direct binding of TRP to TRPR, and also indicated that TRP2 displayed a higher affinity than TRP3 to TRPR.

The detected accumulation of intracellular cAMP concentration only in HEK293 cells transformed with TRPR (Fig. 4A) confirmed that TRP2 and TRP3 can activate TRPR and trigger cAMP signaling. This effect was confirmed in a second experiment and compared to other neuropeptides, including short neuropeptide F (NPF), pigment spreading hormone (PSH), and corazonin (CRZ), which did not induce any detectable responses in both HEK293 cells (Fig. 4B) and Sf21 cells (Fig. 4C).

**Fig. 4:**
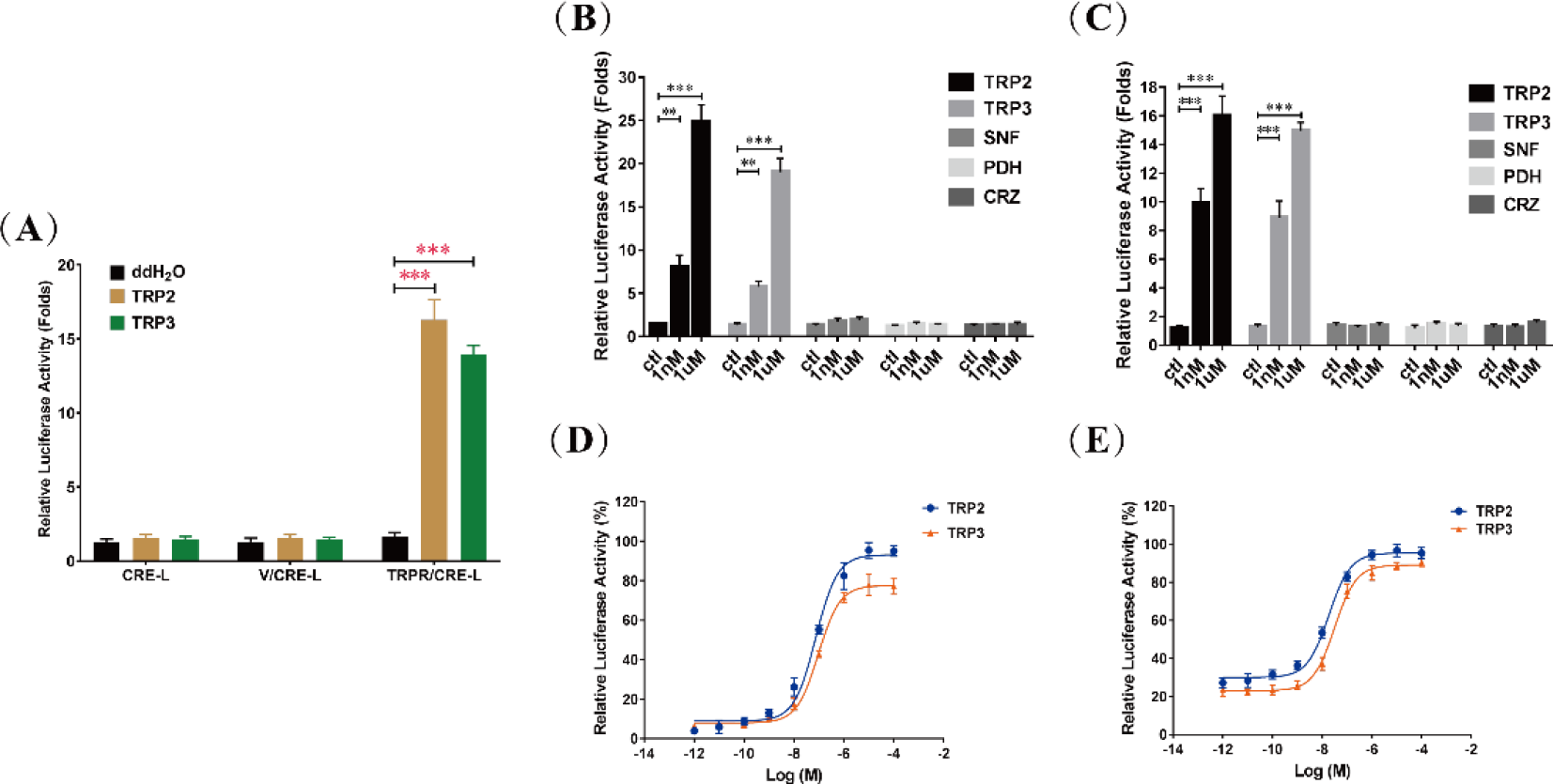
TRP/TRPR-mediated cAMP accumulation in cells. (A), Luciferase activity of HEK293 cells transfected with the reporter gene pCRE-Luc (CRE-L), and co-transfected with pFLAG-TRPR (TRPR) or vehicle vector (V) were determined in response to ddH_2_O and TRP (TRP2 or TRP3, 1 μM) treatment. TRP-dependent TRPR activation increases cAMP levels more than 10-fold. Luciferase activity of HEK293 cells (B) and Sf21 cells (C) co-transfected with TRPR and CRE-L were determined in response to different neuropeptides (TRP2, TRP3, short neuropeptide F (SNF), pigment-dispersing hormone (PDH), and corazonin (CRZ)) at different concentrations (1 nM or 1 μM). Increase of cAMP was specific to TRP2 and TRP3. Dose-dependent changes of luciferase activities, indicating cAMP increases, in HEK293 cells (D) and Sf21 cells (E) co-transfected with TRPR and CRE-L revealed typical kinetics in response to TRP2 and TRP3. All data are presented as mean ± s.e.m. from three independent experiments. Student’s t-tests were used for pairwise comparisons (**p<0.01, ***p<0.001).

Additional dose-depended assays of TPR2 and TPR3 on cAMP accumulation in both HEK293 cells (Fig. 4D) and Sf21 cells (Fig. 4E) confirmed the direct correlation between TRP stimulation and cAMP signaling, and indicated that TRPR was more sensitive to TRP2 than to TRP3. Further analysis showed that pretreatment with PTX (an inhibitor of G_αi_ subunit) had no effect on cAMP accumulation, whereas stimulation with CTX (an activator of G_αs_ subunit) elicited a dramatically increase in abundance of cAMP (Fig. 5A), suggesting that G_αs_ was involved in TRPR-mediated cAMP signaling. In addition, TRP-induced cAMP generation was significantly inhibited by G_αq_ inhibitor YM-254890, and PKA inhibitor H89 (Fig. 5B). Collectively, these results established that both G_αs_ and G_αq_ are involved in TRP/TRPR-mediated cAMP signaling.

**Fig. 5:**
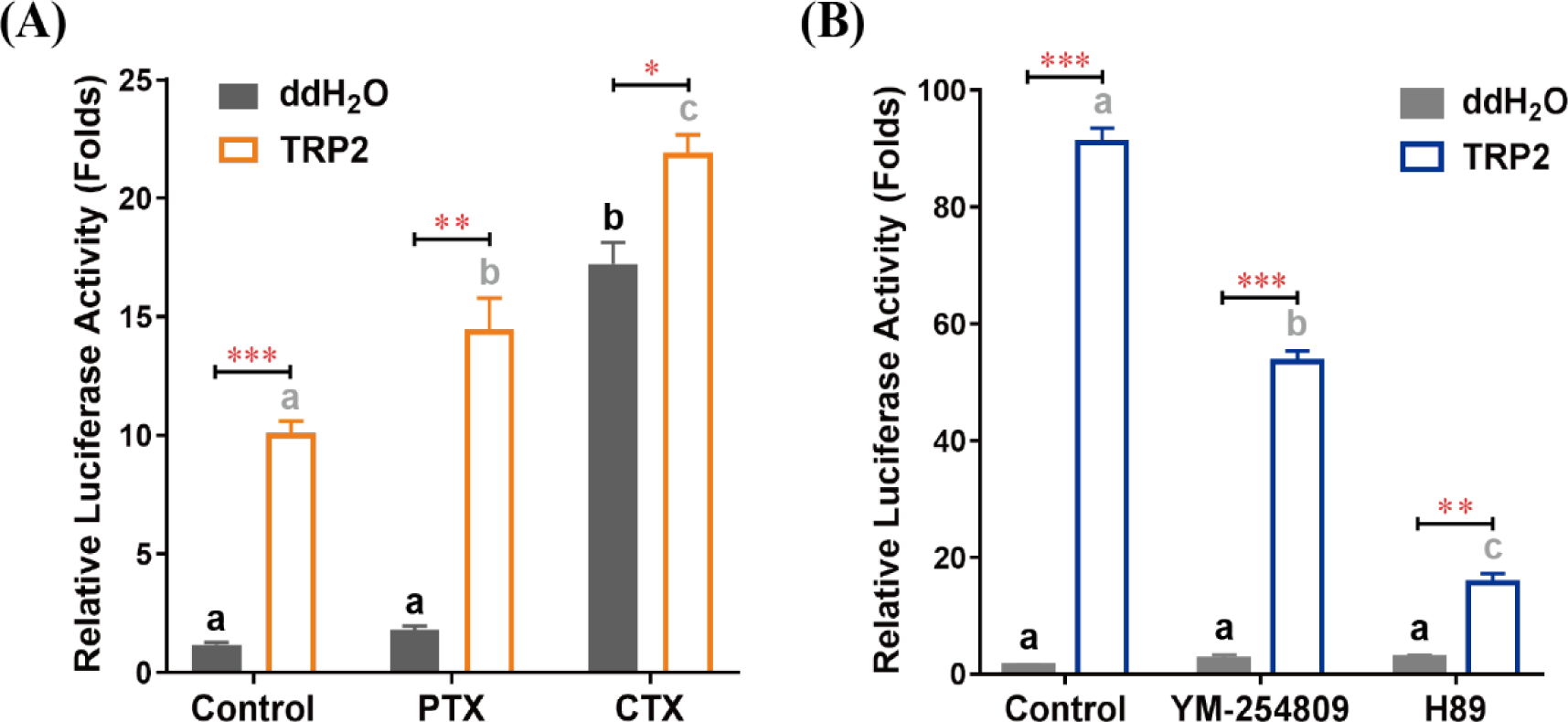
TRP/TRPR signaling induces cAMP accumulation via G_αq_ and G_αs_ pathways. (A), Effects of G_αi_ inhibitor pertussis toxin (PTX) and G_αs_ activator cholera toxin (CTX) on TRP2-mediated stimulation of cAMP accumulation. HEK293 cells expressing TRPR were pretreated with PTX (100 ng/ml) or CTX (300 ng/ml) overnight prior to treatment with TRP2 (1 μM). (B), Effects of G_αq_ inhibitor YM-254890 and PKA inhibitor H89 on TRP2-mediated stimulation of cAMP accumulation. HEK293 cells expressing TRPR were pretreated with YM-254890 (1 μM) or H89 (10 μM) for 2 hours prior to treatment with TRP2 (1 μM). All data are presented as mean ± s.e.m. from three independent experiments. Student’s t-tests were used for pairwise comparisons between water and TRP2 treatments (*: p<0.05, **: p<0.01, ***: p<0.001). One-way ANOVAs followed by Tukey’s post-hoc tests were used for comparisons among control, PTX, and CTX groups within water or TRP2 treatments, and significant differences (p < 0.05) are denoted by letters.

Measurements of a Ca^2+^-sensitive fluorescent indicator suggested that intracellular Ca^2+^ signaling was also elicited by TRP/TRPR signaling. Both, TRP2 and TRP3, could induce a rapid intracellular Ca^2+^ accumulation in HEK293 cells (Fig. 6A) and Sf21 cells (Fig. 6B). The TRP/TRPR-mediated intracellular Ca^2+^ mobilization was decreased by G_αq_ inhibitor YM-254890 and phospholipase C (PLC) inhibitor U73122 (Fig. 6C), suggesting the G_αq_/PLC pathway was involved in TRP/TRPR-mediated Ca^2+^ signaling.

**Fig. 6:**
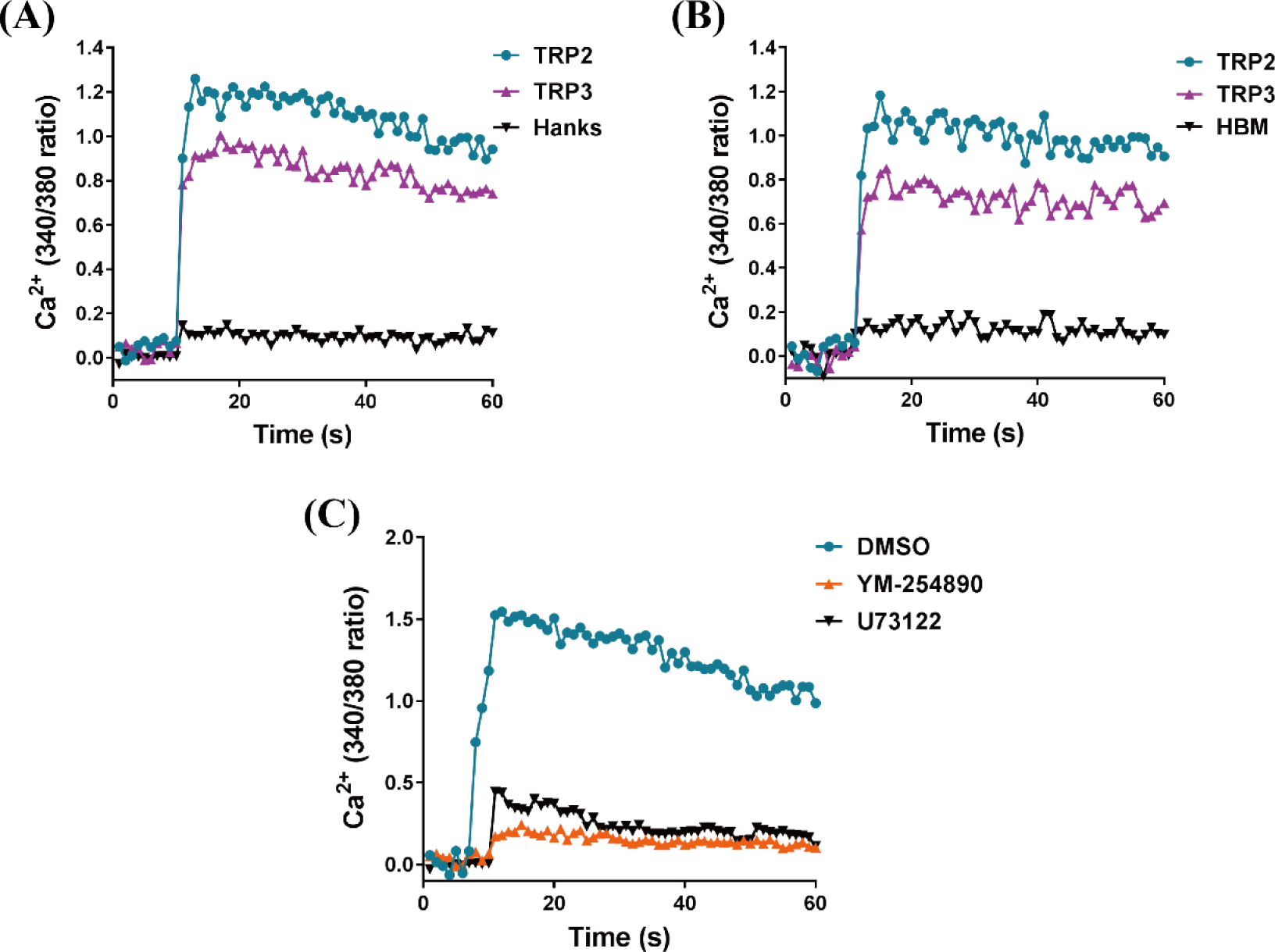
TRP/TRPR-mediated intracellular Ca^2+^ influx via G_αq_/PLC pathways. HEK293 cells (A) and Sf21 cells (B) expressing TRPR were measured in response to TRP2 and TRP3 using the fluorescent Ca^2+^ indicator Fura-2 AM. Hanks solution (Hanks) and Hepes-buffered medium (HBM) were used as a control, respectively. (C), Effects of G_αq_ inhibitor YM-254890 and PLC inhibitor U73122 compared to vehicle control DMSO on TRP2-mediated intracellular Ca^2+^ influx. HEK293 cells expressing TRPR were pretreated with YM-254890 (1 μM) or U73122 (10 μM) for 2 hours then stimulated with TRP2 (1 μM). Each figure is representative of three independent repliates of each experiment.

Western blot analyses proved that phosphorylation of ERK was induced by TRP/TRPR signaling. Treatment with different concentrations of TRP2 induced a dose-dependent phosphorylation of ERK in both HEK293 (EC_50_=68.04 nM) and Sf21 (EC_50_=1.68 nM) cells (Fig. 7A and 7B). Further time-dependent analysis indicated that TRP2 elicited transient phosphorylation of ERK with maximal phosphorylation at 2 min and near basal levels by 90 min (Fig. 7C). Moreover, specific inhibitors were used to elucidate TRP/TRPR signaling-mediated ERK activation in both HEK293 and Sf21 cells. Treatment with MEK inhibitor U0126, PKA inhibitor H89, and PKC inhibitor Go6983, respectively, led to a significant inhibition of TRP/TRPR-mediated ERK activation, whereas G_αi_ inhibitor PTX had no effect, demonstrating that honeybee TRP/TRPR signaling dually coupled to G_αs_ and G_αq_ proteins to activate the ERK signaling pathway (Fig. 7D).

**Fig. 7:**
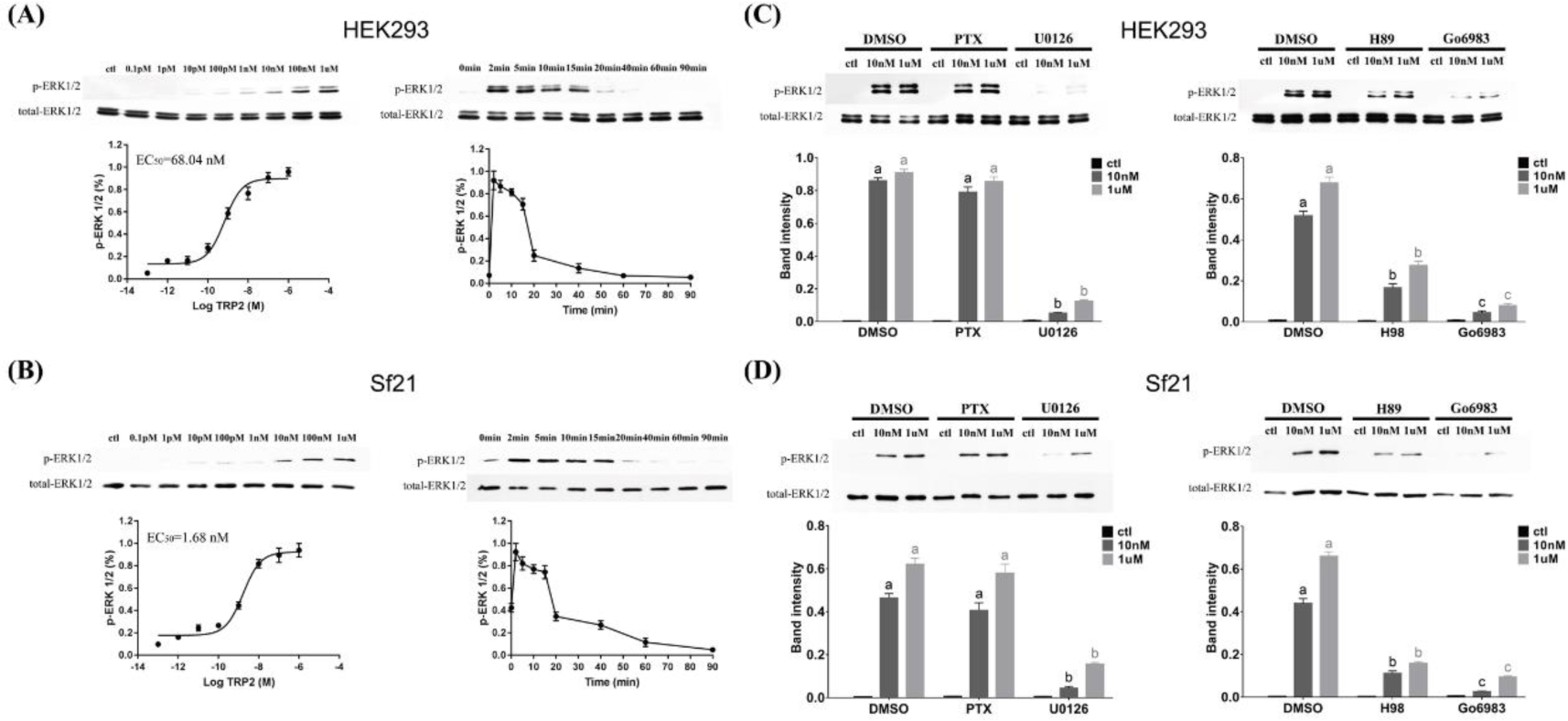
G_αq_/PKC and G_αs_/PKA pathways involved in TRP/TRPR-induced ERK1/2 phosphorylation. Dose- and time-response analyses of TRP/TRPR-induced ERK1/2 phosphorylation in HEK293 cells (A) and Sf21 cells (B). Cells expressing TRPR were serum-starved then incubated either with an increasing dose of TRP2, (from 0.1 pM to 1 μM) for 10 min or with 100 nM TRP2 for different times (from 0 to 90 min), then harvested to quantify ERK1/2 phosphorylation. Effects of G_αi_ inhibitor pertussis toxin (PTX), MEK inhibitor U0126, PKA inhibitor H89, and PKC inhibitor Go6983 on TRP2-induced ERK1/2 phosphorylation in HEK293 cells (C) and Sf21 cells (D). The cells were pretreated with or without inhibitors for 2 hours then stimulated with ddH_2_O (control) or TRP2 (10 nM or 1 μM) for 10 min. The phosphorylated ERK was normalized to a loading control (total ERK). All data are presented as mean ± s.e.m. from three independent replicates, and blots shown are representative of these experiments. One-way ANOVAs followed by Tukey’s post-hoc tests were used for multi-group comparisons, and significant differences (p < 0.05) are denoted by letters.

### 2.4 TRP/TRPR signaling acts as negative regulator of task-specific responsiveness

#### 2.4.1 TRP2 injection decreases task-specific responsiveness

Task-specific responsiveness of the different behavioral phenotypes (PFs, NFs, and NBs) was decreased by injection of TPR2 in a task-specific manner (Fig. 8, Table S8).

**Fig. 8:**
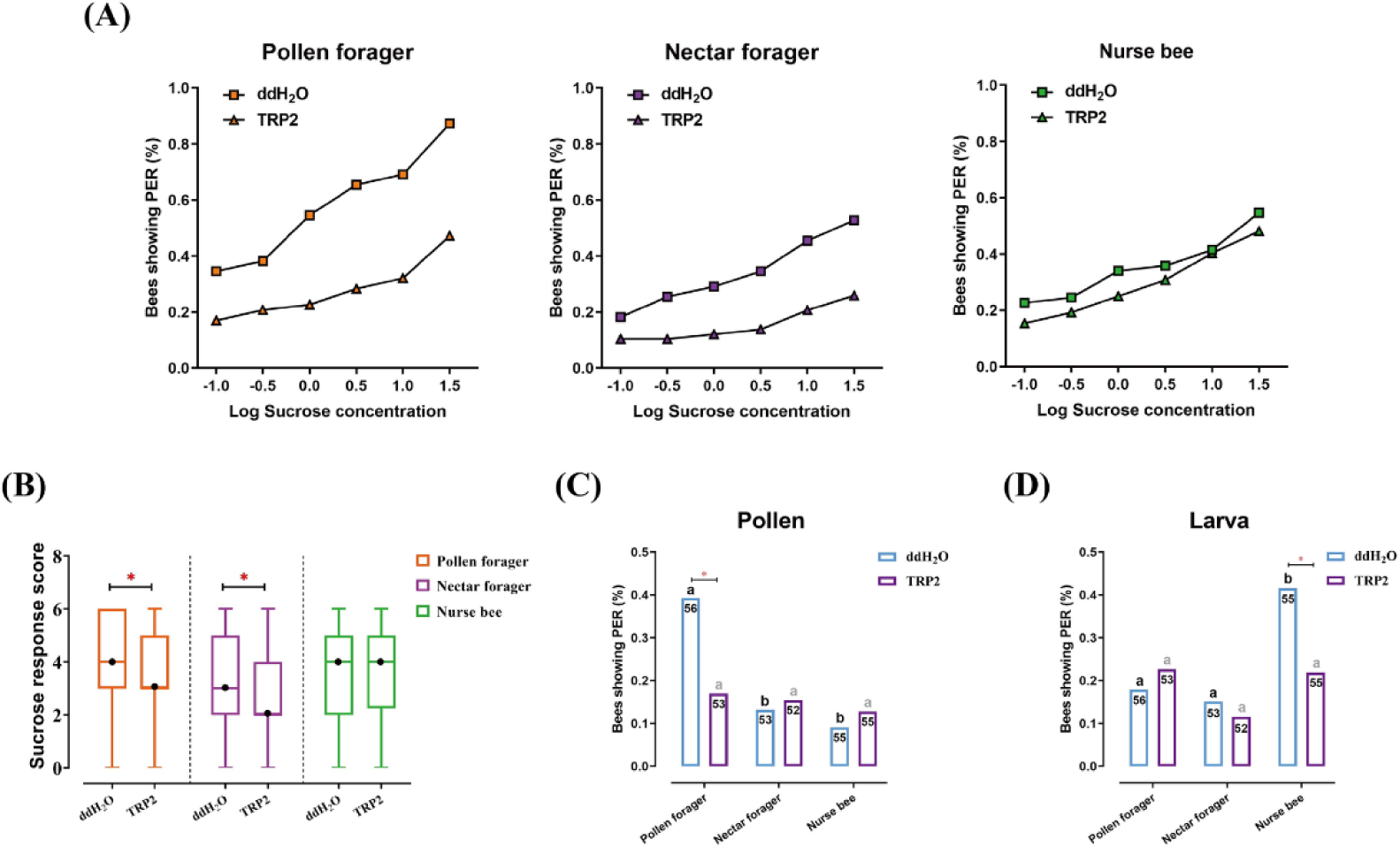
Injection of TRP2 decreases task-specific responsiveness of worker bees. (A) The proportion of pollen foragers (PFs), nectar foragers (NFs), and nurse bees (NBs) exhibiting a positive proboscis extension reflex (PER) increases with increasing concentrations of sucrose solutions but is overall decreased in PFs and NFs after injection of TRP2 compared to ddH_2_O injection. (B) Median sucrose response scores (SRS; intermediate lines) and quartiles (upper and lower lines) of ddH_2_O injected and TRP2 injected groups of PFs, NFs, and NBs. Mann-Whitney U tests were used to compare the SRS (*: p < 0.05). The proportion of PFs, NFs, and NBs showing PER to pollen stimulation (C) and larva stimulation (D) after injection of TRP2 or ddH_2_O. Numbers in bars are the number of individuals sampled in each group. Independent Chi-square tests were used to compare the responsiveness between different treatments (*: p < 0.05) and between different behavioral phenotypes within treatments (significant differences are denoted by letters, p < 0.05).

Injection of the TRP2 peptide significantly reduced the SRS of PFs (Z = 2.2, *p* = 0.031), significantly reducing PER responses to all sucrose concentrations used. Similarly, NFs injected with TRP2 displayed significantly lower SRS than control-injected NFs (Z = 2.3, *p* = 0.019), significantly reducing PER responses to all sucrose concentrations except 0.1% (Fig. 8A and 8B). In contrast, TRP2-injected NBs did not show significant responsiveness changes to sucrose relative to controls. For pollen stimulation, PFs showed significantly decreased responsiveness to pollen loads after TRP2 injection (*χ*^2^ = 6.7, *p* = 0.017), while no significant effects were observed in PFs and NFs (Fig. 8C). In the larval responsiveness assay, injection of TRP2 only significantly affected the responsiveness of NBs (*χ*^2^ = 6.1, *p* = 0.001) but not NFs or PFs (Fig. 8D).

#### 2.4.2 Downregulation of *TRP* or *TRPR* increased task-specific responsiveness

The function of TPR/TRPR signaling on task-specific responsiveness was further confirmed by RNAi-mediated downregulation of *TRP* or *TRPR* that complemented the results of the TRP2 injection.

Knockdown efficiencies were close to 60% for *TRP* and *TRPR* mRNA levels at 24 hours post-injection of the corresponding dsRNA (Fig. S1). Therefore, subsequent PER assays were performed 24 hours after dsRNA injection. Relative to control injections, knockdown of either *TRP* or *TRPR* significantly increased the SRS of NFs (ds*TRP*: Z = 2.4, *p* = 0.049; ds*TRPR*: Z = 2.6, *p* = 0.025), specifically increasing the responses of NFs to sucrose at concentrations of 0.1% (ds*TRP*: *χ*^2^ = 3.9 *p* = 0.039; ds*TRPR*: *χ*^2^ = 4.9, *p* = 0.023), 0.3% (ds*TRP*: *χ*^2^ = 5.3, *p* = 0.018; ds*TRPR*: *χ*^2^ = 4.3, *p* = 0.030), 1.0% (ds*TRP*: *χ*^2^ = 7.0, *p* = 0.007; ds*TRPR*: *χ*^2^ = 6.6, *p* = 0.009), and 3.0% (ds*TRP*: *χ*^2^ = 6.0, *p* = 0.012; ds*TRPR*: *χ*^2^ = 7.4, *p* = 0.006) (Fig. 9A and 9B, Table S9 and S10). Knockdown of *TRP* and *TRPR* didn’t significantly change the overall SRS of PFs and NBs, although it significantly increased the responses of PFs to sucrose at concentrations of 0.1% (ds*TRP*: *χ*^2^ = 4.4, *p* = 0.029; ds*TRPR*: *χ*^2^= 6.1, *p* = 0.011), 0.3% (ds*TRP*: *χ*^2^ = 5.2, *p* = 0.018; ds*TRPR*: *χ*^2^ = 6.0, *p* = 0.011), and 1.0% (ds*TRP*: *χ*^2^ = 5.0, *p* = 0.020; ds*TRPR*: *χ*^2^ = 4.7, *p* = 0.025). Responses to pollen stimulation after dsRNA injection indicated that knockdown of either *TRP* or *TRPR* specifically increased the pollen responsiveness of PFs (ds*TRP*: *χ*^2^ = 6.5, *p* = 0.018; ds*TRPR*: *χ*^2^ = 6.4, *p* = 0.010), whereas the effects on NFs and NBs were not significant (Fig. 9C). The responsiveness of NBs to larvae was significantly increased after gene knockdown of either *TRP* (*χ*^2^ = 4.4, *p* = 0.029) or *TRPR* (*χ*^2^ = 4.8, *p* = 0.023) but NFs and PFs were not affected (Fig. 9D).

**Fig. 9:**
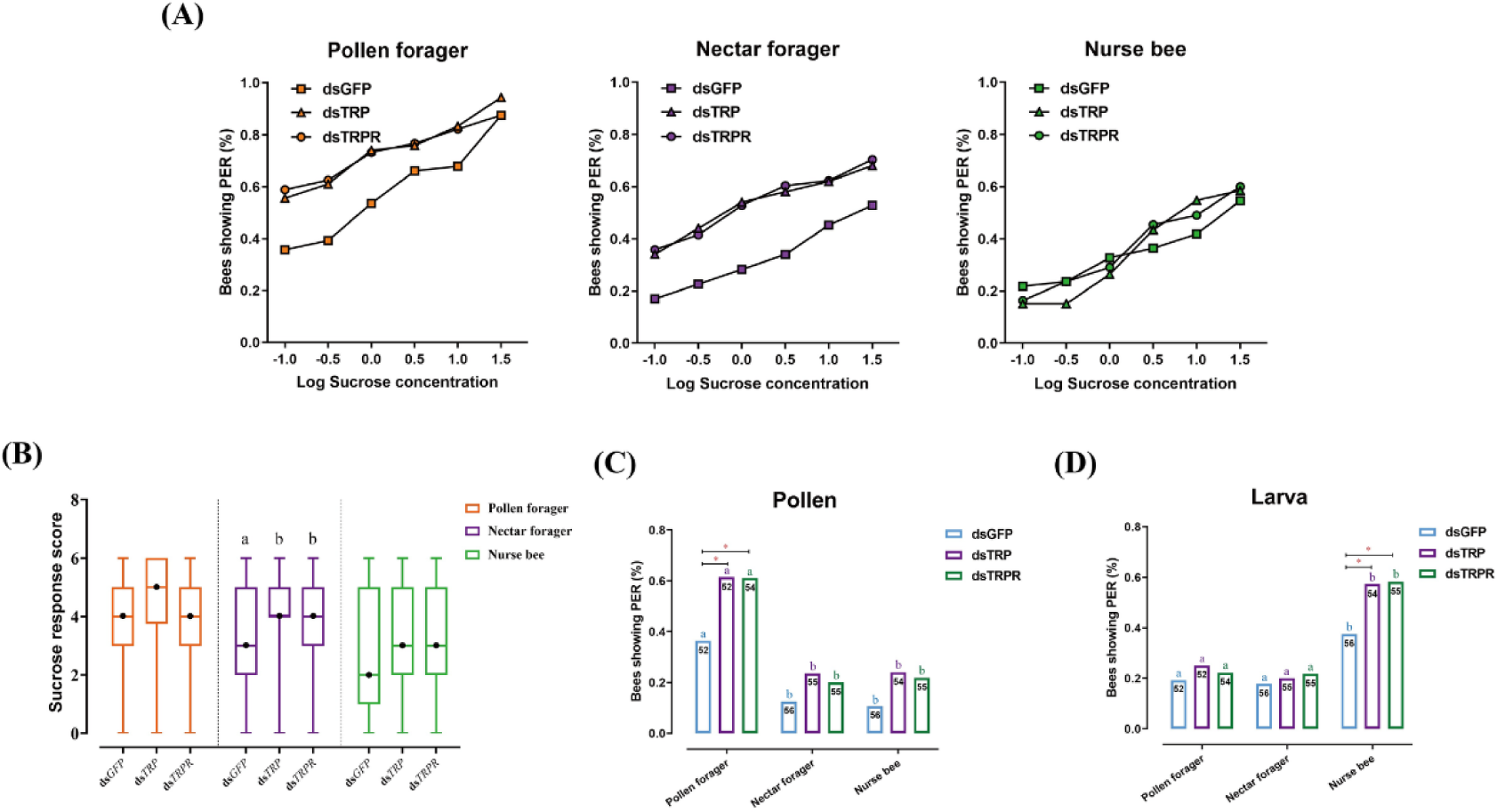
RNAi-mediated knockdown of *TPR* and *TRPR* expression alter task-specific responses of worker bees. (A) Proportion of positive proboscis extension reflex (PER) responses of pollen foragers (PFs), nectar foragers (NFs), and nurse bees (NBs) increases with increasing concentrations of sucrose solutions but overall increases occur only in PFs and NFs after knockdown of *TPR* or *TRPR* transcripts compared to GFP control. Statistical details of these sucrose responsiveness comparisons are shown in Table S10. (B) Median sucrose response scores (SRS; intermediate lines) and quartiles (upper and lower lines) of ddH_2_O injected and TRP2 injected PFs, NFs, and NBs. Kruskal-Wallis tests with Bonferroni correction were used to compare the SRSs of the three treatment groups of each behavioral phenotype and significant differences are denoted by letters (p < 0.05). The proportion of PFs, NFs, and NBs showing PER to pollen stimulation (C) and larvae stimulation (D) after *GFP*, *TPR*, or *TRPR* knockdown. Numbers in bars are the number of individuals sampled in each group. Independent Chi-square tests were used to compare the task-specific responsiveness between different treatments (*: p < 0.05, **: p < 0.01) within behavioral phenotypes and between different behavioral phenotypes within each treatment (significant differences are denoted by letters, p < 0.05).

### 2.5 TRP/TRPR signaling regulates ERK signaling *in-vivo*

To complement our finding that TRP/TRPR signaling activates ERK phosphorylation in cell culture, we used our *in-vivo* manipulations of TRP-signaling to confirm the link between TRP- and ERK signaling in living honeybee workers. Western blot results confirmed that TRP/TRPR signaling triggers ERK signaling *in vivo*. The level of phosphorylated ERK significantly increased after injection of TRP2 peptide into NBs, PFs, and NFs (Fig. 10A) and decreased after knockdown of the *TRP* or *TRPR* transcripts (Fig. 10B).

**Fig. 10:**
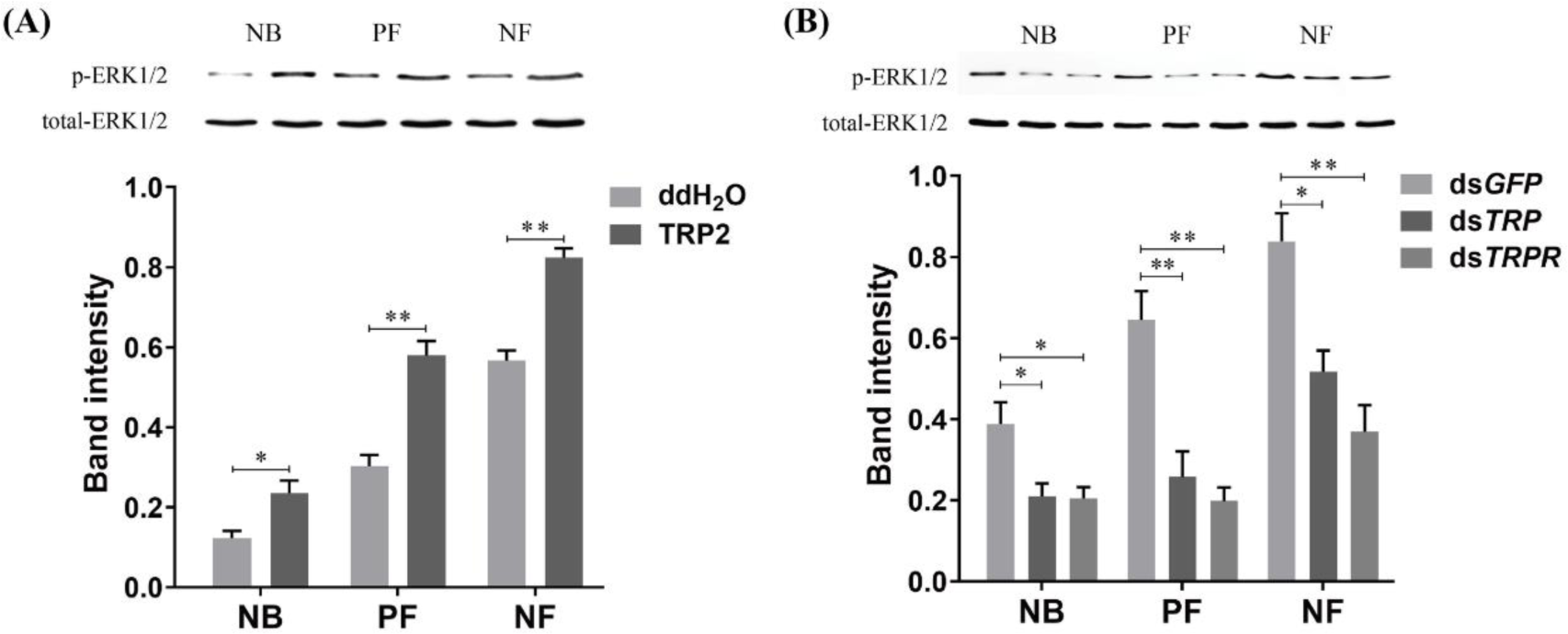
Manipulations of TRP and TRPR levels change ERK phosphorylation states in the worker bee brains. (A) The ERK phosphorylation (p-ERK) levels after injection of TRP2 or ddH_2_O into pollen foragers (PFs), nectar foragers (NFs), and nurse bees (NBs) of *Apis mellifera ligustica*. (B) The p-ERK levels after transcript knockdown of *GFP*, *TPR*, or *TRPR* in PFs, NFs, and NBs. The p-ERK was normalized to a loading control (total-ERK). The data shown are representative of three independent experiments, and blots shown are representative of these experiments. Student’s t-tests were used for pairwise comparisons between control and treatment groups within each behavioral phenotype (*: p < 0.05, **: p < 0.01, ***: p < 0.001).

## 3. Discussion

Behavioral plasticity plays a central role in animal adaptation and modulating behavioral responsiveness to different stimuli and contexts is key to individual fitness. The success of social insects is partly due to their efficient division of labor, a form of behavioral plasticity among instead of within individuals. In this study, we demonstrated that the responsiveness to task-relevant stimuli correlates with behavioral specialization in two different honeybee species. Through parallel characterization of the neuropeptidome, we identified two tachykinin-related peptides (TRP2 and TRP3) as putative mechanism to adjust task-specific response thresholds and thus proximally guide division of labor. Subsequently, we characterized the molecular action of TRP2 and TRP3 in cell culture by verifying their binding to their membrane-bound receptor and demonstrating activation of multiple down-stream signaling mechanisms. Finally, we verified causal involvement of TRP signaling in modulating task-specific behavioral response thresholds through complementary outcomes of TRP2 injection and RNAi-mediated knockdown of *TRP* and its receptor *TRPR*: while injection decreased task-specific responses, down-regulation of TRP or TRPR increased the same specific responses. Thus, we present the first process that tunes the behavioral responsiveness of animals to specific stimuli compared to others. We use behaviorally specialized honeybee workers as models but hypothesize that this function of TRP signaling could be more widely conserved to adjust the context-specificity of behavioral responses in animals.

Among all the signaling molecules in the nervous system, neuropeptides represent the largest and most diverse category and are crucial in orchestrating various biological processes and behavioral actions [50, 51]. Thus, we quantitatively compared the entire neuropeptidome among three behavioral worker phenotypes of *Apis mellifera ligustica* (AML) and *Apis cerana cerana* (ACC) without an *a-priori* assumption. In addition to characterizing the ACC neuropeptidome for the first time and discovering several new neuropeptides from the AML brain, we identified TRP2 and TRP3 as candidates. TRPs have been associated with the modulation of appetitive olfactory sensation [52–54], sex pheromone perception [41], and aggression [55]. Particularly in honeybees, *TRP* is preferentially expressed in the mushroom body and some neurons scattered in the antennal and optic lobes [56]. This expression is consistent with our hypothesis that TRP-signaling may be a general modulator of behavioral responsiveness. TRPs expression in the honeybee worker brain increases during the transition from nursing to foraging, further implicating it in the regulation honeybee social behavior [49, 57].

In our study, only expression of TPR2 and TRP3 varied consistently among behavioral phenotypes of AML and ACC. In both species, TRP2 and TRP3 were most abundant in the brain of NFs, followed by PFs, and finally NBs. This is consistent with the very specific responsiveness of NBs to brood stimuli observed in our PER experiments, while the responsiveness of PFs and NFs was successively less specific: PFs responded specifically to two stimuli, while NFs did not show specifically strong responses to any stimuli. Moreover, the comparison between AML and ACC indicated higher TRP2 and TRP3 abundance in ACC in each behavioral phenotype, commensurate with the less specific PER responsiveness in ACC compared to AML. A few other neuropeptides, such as apidaecins, diuretic hormone, and prohormone-3, showed somewhat similar expression patterns in both species, but none of these was as tightly correlated to behavioral responsiveness and none has previously been connected with behavioral regulation in insects or other animals. Therefore, the TRPs were chosen as candidates of the control of honeybee division of labor for subsequent functional tests and molecular characterization.

The action of most insect neuropeptides is mediated by binding to G-protein–coupled receptors (GPCRs) and often involves cAMP and Ca^2+^ as second messengers [58]. The TRPR is activated by TRPs triggering intracellular cAMP accumulation and Ca^2+^ mobilization in fruit flies and silkworms (*Bombyx mori*) [59, 60], while no cAMP-responses were discovered in stable flies (*Stomoxys calcitrans*) [61]. The results of our peptide-based binding assays functionally confirmed that the honeybee TRPR is indeed the receptor for TRP2 and TRP3. The subsequent functional assays revealed that TRP signaling results in a dose-dependent increase in both intracellular cAMP and Ca^2+^. Together, these results indicate that TRPs can activate TRPR and trigger second messengers to regulate downstream functions. TRP2 displayed a higher affinity to TRPR and induced higher cAMP and Ca^2+^ signaling than TRP3, leading us to focus on TRP2 in the later *in vivo* experiments. Moreover, TRP signaling is sensitive to G_αs_ activation and is significantly blocked by G_αq_ and PKA inhibitors, suggesting both G_αs_ and G_αq_ are involved in TRP signaling in honeybees. Many GPCRs are able to induce mitogen-activated protein kinase (MAPK) cascades via cooperation of G_αs_, G_αq,_ and G_αi_ signals, leading to the phosphorylation of ERK1/2, which plays critical roles in diverse biological processes [62]. Our results indicate that honeybee TRP signaling mediates phosphorylation of ERK1/2 in a dose- and time-dependent manner in both HEK293 and Sf21 cells. In addition, ERK1/2 activation was significantly inhibited by the PKA, PKC, and MEK inhibitors, which is in line with the observation of intracellular cAMP accumulation and Ca^2+^ mobilization. Thus, honeybees seem to be very similar to silkworms with regards to the involvement of the G_αs_/cAMP/PKA and G_αq_/Ca^2+^/PKC signaling pathways in the regulation of TRP-induced ERK1/2 activation [60]. Taken together, our results demonstrate that the honeybee TRPR is specifically activated by TRPs, eliciting intracellular cAMP accumulation, Ca^2+^ mobilization, and ERK phosphorylation by dually coupling G_αs_ and G_αq_ signaling pathways.

Our *in vitro* and *in vivo* demonstrations that TRP signaling activates the ERK putatively link TRP signaling also to the insulin/insulin-like signaling (IIS) pathway. IIS is controlled by neuropeptides through ERK in *Drosophila* [63], and this connection in honeybees ties TRP back to the age-based division of labor among workers: IIS signaling influences the timing of the behavioral maturation of honeybee workers and brain *AmIlp1* is significantly higher expressed in foragers than nurses [64], consistent with our finding that TRPs are higher in foragers than nurses. Numerous other physiological changes accompany the transition from in-hive nurse bees to foragers [37, 65–67] and our results integrate TRPs as the most important neuropeptides into the regulation of the behavioral ontogeny of honeybee workers and potential feedback loops to the modulation of behavioral response thresholds. The specialization of nectar and pollen foragers has also been linked to IIS signaling [68, 69] and explained by differences in sucrose response thresholds [70]. Our findings here may connect the differences in response thresholds and IIS mechanistically through the TRP and ERK signaling pathways.

The PER paradigm is well-suited to test behavioral response thresholds and has been used for over 50 years in honeybees [36]. Consistent with previous studies, we found pollen foragers to be more responsive to sucrose than nectar foragers and nurses in *Apis mellifera* [32]. Moreover, we found corresponding differences between these behavioral groups in the closely related *Apis cerana*. The pollen forager’s responsiveness to low sucrose concentrations might also make them more responsive to pollen, but the causation of the PER to pollen is unclear [71] and other components of pollen may functionally distinguish pollen from sucrose responsiveness [34]. Our results support the view that pollen and sucrose are distinct stimuli: While our experimental manipulations of TRP signaling altered the responsiveness of pollen foragers to pollen and sucrose, only responsiveness to sucrose was affected in nectar foragers and only responsiveness to larvae was affected in nurses. The functional significance of the PER in response to larvae is currently unclear, but we show that it is specific to nurses and it has previously been linked to brood provisioning [35]. Thus, our diverse PER results in two species comprehensively support the hypothesis that task-specific response thresholds guide behavioral specialization, leading to division of labor among honeybee workers [21–23].

TRPs may adjust specific sensory neural circuits, potentially acting in concert with other neuromodulators [72, 73]. However, we have currently no evidence to support the hypothesis of different molecular TRP actions in different stimulus-response pathways and our consistent results from two very different cell cultures indicate that TRP signaling may be relatively robust to the cellular environment. Thus, we favor the more parsimonious explanation is that TRP signaling acts generally through the identified mechanisms to decreases task-specific response thresholds of behavioral specialists: It decreases pollen and sucrose responsiveness in pollen foragers, sucrose responsiveness in nectar foragers, and responsiveness to larvae in nurses. TRP signaling may thus be a general regulator of how task-specific stimuli are weighted relative to others and consequently how specialized behavioral specialists are. This effect translates to different degrees of division of labor in social insect colonies and may control the context-specificity of behavioral responses in animals more generally [74].

Although AML and ACC are close relatives with similar basic biology, some behavioral differences have evolved since their speciation [75]. AML and ACC share the age-based division of labor, with younger bees specializing on nursing before maturing to foraging activities [76] and ACC foragers also specialize in nectar or pollen collection [77] similar to AML [28]. Accordingly, we found the main differences of stimulus responsiveness and TRPs expression among worker phenotypes conserved. However, ACC exhibited less responses to the task-specific stimuli than AML. Consistent PER differences in AML and ACC between nectar and pollen foragers and a generally lower responsiveness of ACC have been identified before [78], but the biological interpretation has remained unclear. It is possible that the species differences arise due to methodological bias, favoring AML performance in PER assays. However, our study offers the alternative explanation that ACC workers are less specialized than AML workers due to higher TRP signaling. Lower innate specialization may accompany better learning of ACC [79], facilitating its more opportunistic worker task allocation and resource exploitation than AML [80]. These alternative life history strategies are plausible, given the typical differences in colony size and habitat [73, 74, 81]. All three worker phenotypes of ACC exhibited higher levels of TRPs than their AML counterparts but functional verification at the level of colony phenotypes will be required to unambiguously link TRP signaling to such interspecific differences in life history.

## 4. Materials and Methods

### 4.1. Honeybee sources and sampling

Two honeybee species, *Apis mellifera ligustica* (AML) and *Apis cerana cerana* (ACC), were maintained in the apiary of the Institute of Apicultural Research at the Chinese Academy of Agricultural Sciences in Beijing. Three colonies of each species with mated queens of identical age were selected as experimental colonies, and before experiments the colonies were equalized in terms of adult bee population, brood combs, and food storage. Frames containing old pupae (1-2 days before emergence) were put into an incubator (34°C and 80% relative humidity) for eclosion. Newly emerged worker bees were paint marked on their thoraxes and placed back into their parent colonies. Ten days later, marked bees that had their head and thorax in open brood cells while contracting their abdomen for more than 10 seconds were collected as nurse bees (NBs). Twenty day-old, marked bees were collected during early morning (between 8:00 am and 10:00 am) in good weather conditions during the blooming period of black locusts (*Robinia pseudoacacia* L.) as forager bees. The entrance to the hives were blocked to facilitate collecting. Bees flying into the hive with pollen loads were collected as pollen foragers (PFs), returning foragers without pollen loads were collected as nectar foragers (NFs). The experimental design of six groups (three behavioral phenotypes in two species) was used to compare responsiveness to task-specific stimuli (section 4.2) and to relate these phenotypes to differences in the brain neuropeptidome (section 4.3).

### 4.2. Comparative Proboscis Extension Reflex (PER) experiments

To investigate the responsiveness of different worker bee behavioral phenotypes (NBs, PFs, and NFs of AML and ACC) to different stimulus modalities (sucrose solution, pollen, and larva), series of PER experiments were performed. One hundred bees of each behavioral phenotype were collected from each experimental colony in the morning, transferred to the laboratory and narcotized on ice, then harnessed using a previously described protocol [82]. All harnessed bees were fed to satiation with 50% sucrose solution and placed in a dark incubator (20°C and 65% relative humidity) overnight. After 24 hours, all surviving bees were assayed for their PER following the methodology of Page et al. [32]. Each stimulus was assessed independently with a new set of bees.

To investigate the sucrose responsiveness, bees were assayed using an ascending order of sucrose concentrations: 0.1, 0.3, 1, 3, 10, and 30% (weight/weight). A small droplet of each solution was touched to the bees’ antennae for 3 seconds and the extension of the proboscis was monitored during this time. The interval between each sucrose solution trial was 5 min to exclude sensitization or habituation effects. The total number of PER responses after stimulation with the six different sucrose concentrations were combined into a sucrose response score (SRS) of a bee [83–85]. The SRSs of the three behavioral phenotypes in the same species were compared using Kruskal-Wallis tests with Bonferroni correction. Pairwise Mann-Whitney U tests were used for comparing the same phenotype from two honeybee species. The sucrose responsiveness for specific sucrose concentrations was further compared between different groups with independent Chi-square tests.

To test pollen stimulation, fresh pollen loads that had been removed from the leg of randomly selected pollen foragers of the test group were used: AML were tested with pollen collected by AML foragers and ACC with pollen collected by ACC foragers. These loads contained a mixture of different pollen, predominated by black locust (*Robinia pseudoacacia*). As a control for mechanical stimulation, each bee had both antennae first touched with a piece of filter paper, and spontaneous responders were excluded. Subsequently both antennae of each bee were gently touched with a pollen load and PER responses were recorded. The pollen responsiveness was compared with independent Chi-square tests between different groups.

To test responsiveness to larva, one-day-old larvae from each honeybee species were collected, briefly rinsed in distilled water to remove royal jelly residue and dried on a filter paper. As before, both antennae of bees were touched with a piece of filter paper first, and spontaneous responders were excluded, then PERs in response to a larva touching the antennae were recorded. The responsiveness to larvae was compared with independent Chi-square tests between different groups. Statistical analyses were conducted using SPSS 20.0 (IBM, USA).

### 4.3. Quantitative comparisons of brain neuropeptidomes

To explore brain neuropeptide functions in behavioral regulation, a label-free quantitative strategy was employed to compare neuropeptidomic variations between behavioral phenotypes and the two honeybee species. Three independent biological replicate samples (120 bees/sample) of NBs, PFs, and NFs of both AML and ACC (18 samples total) were collected and immediately frozen in liquid nitrogen. Individual brains were carefully dissected from the head capsule while remaining chilled on ice, and the dissected brains were frozen at -80°C until neuropeptide extraction.

The brains were homogenized at 4°C by using a 90:9:1 solution of methanol, H_2_O, and acetic acid. The homogenates were centrifuged at 12000g for 10 min at 4°C. The resulting supernatant containing the neuropeptides was collected and dried. The extracted neuropeptide samples were dissolved in 0.1% formic acid in distilled water, and the peptide concentration was quantified using a Nanodrop 2000 spectrophotometer (Thermo Fisher Scientific, USA). LC-MS/MS analysis was performed on an Easy-nLC 1200 (Thermo Fisher Scientific) coupled Q-Exactive HF mass spectrometer (Thermo Fisher Scientific). Buffer A (0.1% formic acid in water) and buffer B (0.1% formic acid in acetonitrile) were used as mobile phase buffers. Neuropeptides were separated using the following gradients: from 3 to 8% buffer B in 5 min, from 8 to 20 % buffer B in 80 min, from 20 to 30% buffer B in 20 min, from 30 to 90% buffer B in 5 min, and remaining at 90% buffer B for 10 min. The eluted neuropeptides were injected into the mass spectrometer via a nano-ESI source (Thermo Fisher Scientific). Ion signals were collected in a data-dependent mode and run with the following settings: full scan resolution at 70,000, automatic gain control (AGC) target: 3 × 10^6^, maximum inject time (MIT): 20 ms, scan range: m/z 300-1,800; MS/MS scans resolution at 17,500, AGC target: 1 × 10^5^, MIT: 60 ms, isolation window: 2 m/z, normalized collision energy: 27, loop count 10, and dynamic exclusion: charge exclusion: unassigned, 1, 8, >8; peptide match: preferred; exclude isotopes: on; dynamic exclusion: 30 s. Raw data were retrieved using Xcalibur 3.0 software (Thermo Fisher Scientific).

The extracted MS/MS spectra were searched against a composite database of *Apis mellifera* (23,491 protein sequences, downloaded from NCBI on July, 2018) or *Apis cerana* (20,934 protein sequences, downloaded from NCBI on July, 2018) using in-house PEAKS 8.5 software (Bioinformatics Solutions, Canada). Amidation (A, -0.98) and pyro-glu from Q (P, -17.03) were selected as variable modifications. The other parameters used were: parent ion mass tolerance, 20.0 ppm; fragment ion mass tolerance, 0.05 Da; enzyme, trypsin; allowing a nonspecific cleavage at both ends of the peptide; maximum missed cleavages per peptide, 2; maximum allowed variable PTM per peptide, 2. A fusion target-decoy approach was used for the estimation of the false discovery rate (FDR) and controlled at ≤ 1.0% (−10 log P ≥ 20.0) both at protein and peptide levels. Neuropeptide identifications were only used if ≥ 2 spectra were identified in at least two of the three replicates of each sample type.

Quantitative comparison of brain neuropeptidomes was performed by the label-free approach in PEAKS Q module. Feature detection was performed separately on each sample by using the expectation-maximization algorithm. The features of the same peptide from different samples were reliably aligned together using a high-performance retention time alignment algorithm [86]. Peptide features were considered significantly different between experimental groups if pairwise p < 0.01 and fold change ≥ 1.5. A heat map of differentially expressed proteins was created by Gene cluster 3.0 using the unsupervised hierarchical clustering, and the result was visualized using Java Tree view software. The LC−MS/MS data and search results are deposited in ProteomeXchange Consortium (http://proteomecentral.proteomexchange.org) via the PRIDE partner repository with the dataset identifier PXD018713.

### 4.4. Characterization of honeybee tachykinin related peptide (TRP) signaling pathway

To characterize honeybee TRP signaling pathway, the TRP receptor (TRPR) gene was first cloned and expressed in human and insect cell lines to identify its cellular location and verify its binding to TRPs. Additionally, these cells were used to test whether TRP/TRPR signaling triggers intracellular cAMP accumulation, Ca^2+^ mobilization, and ERK phosphorylation.

#### 2.4.1. TRPR gene clone and expression

To amplify the full-length sequence encoding *TRPR* of *Apis mellifera*, primers were designed using Primer Premier 5.0 software (PREMIER Biosoft, USA) based on the sequence from GenBank^TM^ KT232312. The coding sequence of TRPR was amplified and cloned into FLAG-tag expression vectors (pCMV-FLAG and pBmIE1-FLAG) and EGFP-tag expression vectors (pEGFP-N1 and pBmIE1-EGFP). The primers used are documented in Table S11. All constructs were sequenced to verify the correct sequence, orientation, and reading frame of the inserts.

The human embryonic kidney cell line HEK293 and the insect *Spodoptera frugiperda* pupal ovary cell line Sf21 were used for honeybee TRPR expression. HEK293 cells were cultured in DMEM medium (Gibco, USA) supplemented with 10% fetal bovine serum (FBS). Sf21 cells were cultured in TC100 medium (Gibco) supplemented with heat-inactivated 10% FBS. Transfection of HEK293 cells was performed using Lipo6000™ transfection reagent (Beyotime, China), while transfection of Sf21 cells was performed using LipoInsect™ transfection reagent (Beyotime), according to the manufacturer’s instructions.

#### 2.4.2. Cellular location of TRPR

To confirm the location of the honeybee TRPR, receptor surface expression assays were performed. HEK293 or Sf21 cells expressing TRPR-EGFP were seeded onto poly-L-lysine coated glass coverslips and allowed to attach overnight under normal growth conditions. After 24 hours, cells were incubated with the membrane probe DiI (Beyotime) and the nucleic acid probe Hoechst 33342 (Beyotime) at 37°C for 10 min, then fixed with 4% paraformaldehyde for 15 min. Cells transfected with empty EGFP-tag expression vectors were used as a control. The cells were imaged using a Leica SP8 (Leica Microsystems, Germany) confocal microscope equipped with an HC PL APO CS2 63×/1.40 oil objective. Images were acquired with the sequence program in the Leica LAS X software.

#### 2.4.3. Binding of TRPs to TRPR

To confirm the direct binding of the honeybee TRPs to TRPR, competitive binding experiments were performed using synthesized TAMRA-TRP2 (TAMRA-ALMGFQGVRa) and TAMRA-TRP3 (TAMRA-APMGFQGMRa), with TAMRA labeled at the N-terminus. The neuropeptides used as ligands in here and in later sections were commercially synthesized by SynPeptide Co, Ltd (China). All peptides were purified by reverse-phase high performance liquid chromatography with a purity > 98%, lyophilized, and diluted to the desired concentrations for subsequent experiments. The peptide sequences were verified by us using a Q-Exactive HF mass spectrometer (Thermo Fisher Scientific). HEK293 and Sf21 cells expressing FLAG-TRPR were first seeded onto poly-L-lysine-coated 96-well plates and cultured overnight. On the next day, cells were washed once with phosphate-buffered saline (PBS), then incubated with 25 mL TAMRA-TRP2 or TAMRA-TRP3 (10 nM) in the presence of increasing concentrations of unlabeled TRP2 and TRP3 in a final volume of 100 mL of binding buffer (PBS containing 0.2% bovine serum albumin). Cells were incubated at room temperature for 2 hours. Fluorescence intensity was measured with a fluorescence spectrometer microplate reader (Tecan Infinite 200 PRO, Tecan, Switzerland) after washing twice with binding buffer. The cells transfected with empty FLAG-tag expression vectors were used as a control. The binding displacement curves were analyzed by GraphPad Prism 8.0 (GraphPad Software, USA) using the non-linear logistic regression method.

#### 2.4.4. TRP/TRPR signaling targets: cAMP, Ca^2+^, and ERK

To test whether TRP/TRPR signaling affects cAMP accumulation, intracellular cAMP was measured after incubation of HEK293 and Sf21 cells expressing FLAG-TRPR and pCRE-Luc with TRP2 and TRP3. After seeding in a 96-well plate overnight, HEK293 or Sf21 cells co-transfected with pFLAG-TRPR and pCRE-Luc were grown to about 90% confluence.

After washing once with PBS, cells were incubated with the neuropeptides TRP2, TRP3, short neuropeptide F (SNF), pigment-dispersing hormone (PDH), and corazonin (CRZ) in serum-free medium for 4 hours at 37°C for HEK293 cells, and at 28°C for Sf21 cells. Cells transfected with empty EGFP-tag expression vectors were used as a control. Luciferase activity was detected by a luciferase assay system (Promega, USA). Fluorescence intensity was measured with a Tecan fluorescence spectrometer. When characterizing the TRP-mediated cAMP accumulation, cells were pretreated with G_αi_ inhibitor pertussis toxin (PTX), G_αs_ activator cholera toxin (CTX), G_αq_ inhibitor YM-254890, and PKA inhibitor H89before stimulation with TRP2.

To test whether TRP signaling also affects intracellular Ca^2+^ concentrations, intracellular Ca^2+^ was measured after incubation of HEK293 and Sf21 cells expressing FLAG-TRPR with TRP2 or TRP3. Cells were detached by a non-enzymatic cell dissociation solution (Sigma-Aldrich, USA), washed twice with PBS, and resuspended at a density of 5 × 10^6^ cells/ml in HEPES buffered saline (Macklin, China). Cells were then incubated with 3 μM Fura-2 AM (MedChemExpress, USA) for 30 min at 37°C for HEK293 cells, and at 28°C for Sf21 cells. Intracellular Ca^2+^ flux was measured using excitation wavelengths alternating at 340 and 380 nm with emission measured at 510 nm in a Tecan fluorescence spectrometer. When characterizing the detailed TRP-mediated intracellular Ca^2+^ mobilization, cells were pretreated with G_αq_ inhibitor YM-254890 and PLC inhibitor U73122 before stimulation with TRP2.

To assess whether TRP signaling mediates ERK1/2 signaling, ERK1/2 phosphorylation was measured by Western blot analysis after incubation of HEK293 and Sf21 cells expressing FLAG-TRPR with TRP2. Cells were seeded in 24-well plates and starved for 4 hours in serum-free medium to reduce background ERK1/2 activation and eliminate the effects of the change of medium. After incubation with TRP2, cells were lysed by RIPA buffer (Beyotime) at 4°C for 30 min. Protein concentration was determined according to the Bradford method using BSA as the standard and the absorption was measured at 595 nm (spectrophotometer DU800, Beckman Coulter, Los Angeles, CA), then all the samples were kept in -80°C for further use. For Western blot, equal amounts of total cell lysate (20 μg/lane) were fractionated by SDS-PAGE (10%) and transferred to a PVDF membrane (Millipore, USA) using an iBlot dry blotting system (Invitrogen, USA). The membranes were blocked for 2 hours at room temperature and then incubated with rabbit monoclonal anti-pERK1/2 antibody (Cell Signaling Technology, USA) and anti-rabbit horseradish peroxidase-conjugated secondary antibody (Cell Signaling Technology) according to the manufacturers’ protocols. Antibody reactive bands were visualized using Pierce^TM^ ECL western blotting substrate (Thermo Fisher Scientific, USA) followed by photographic film exposure. Total ERK1/2 was assessed as a loading control after p-ERK1/2 chemiluminescence detection. Quantification analyses were performed using Gel-Pro Analyzer 4.0 software (Media Cybernetics, USA).

To explore the detailed TRP-mediated ERK1/2 signaling, cells were pretreated with G_αi_ inhibitor pertussis toxin (PTX), MEK inhibitor U0126, PKA inhibitor H89, and PKC inhibitor Go6983 before stimulation with TRP2.

### 4.5. Effects of TRP2 injection on task-specific responsiveness

To confirm the function of TPR on task-specific responsiveness, NBs, PFs, and NFs of AML were injected with TRP2 and tested for their PER response to sucrose solution, pollen, and larva. About 150 bees of each behavioral phenotype were collected in the morning, then harnessed, fed and placed in a dark incubator as described in section 4.2. After 24 hours, all surviving bees were evenly divided into two groups and injected with 1 μl TRP2 solution (1 μg/μl, synthesized TRP2 dissolved in ddH_2_O) or 1 μl of ddH_2_O into the head of honeybees via the central ocellus using a glass capillary needle coupled to a microinjector. Bees injected with ddH_2_O were used as control. All injected bees were put back to the dark incubator and 1 hour after injection all surviving bees were assayed for their PER to stimulations of sucrose solution, pollen, and larva as described in section 4.2. Each experiment was performed with a new set of bees containing about 55 individuals per experimental and control group.

The average sucrose response scores of the TRP2 injection group and the ddH_2_O injection group were compared separately for each of the three behavioral phenotypes (NBs, PFs, and NFs) using pairwise Mann-Whitney U tests. The sucrose responsiveness was further compared between different groups at each specific sucrose concentration with independent Chi-square tests. The responsiveness to pollen and larvae was compared between TRP2 injection group and ddH_2_O injection group with independent Chi-square tests for each behavioral phenotype separately. All statistical analyses were performed with SPSS Statistics 20.0 (IBM).

### 4.6. Effects of RNAi-mediated downregulation of *TRP* or *TRPR* on responsiveness

To further confirm the hypothesized effects of TPR/TRPR signaling on task-specific responsiveness, RNAi-mediated downregulation of *TRP* and *TRPR* were performed on NBs, PFs, and NFs of AML and then their PER to sucrose solution, pollen, and larva were compared to controls.

Before evaluating the behavioral effects of transcript knockdown of *TRP* or *TRPR*, preliminary experiments were performed to test the dsRNA-mediated knockdown efficiencies of *TRP* and *TRPR*. The dsRNAs of the *TRP* and *TRPR* genes were prepared using the T7 RiboMAX Express RNAi system (Promega). The primers used are listed in Table S11. Sixty bees were randomly collected from each of the three AML colonies. Bees were harnessed, fed with sucrose and put into the dark incubator (20°C and 65% relative humidity) to acclimatize to the experimental conditions. After 30 min, dsRNA (200 ng/bee for *TRP*, 2 μg/bee for *TRPR*) was microinjected into the head of honeybees via the central ocellus using a glass capillary needle coupled microinjector. dsRNA of green fluorescent protein gene (ds*GFP*, 2 μg/bee) was used as control in all RNAi experiments. All harnessed bees were fed with 50% sucrose solution every 12 hours. At 0, 12, 24, and 48 hours after injection, a group of 6 individual bees were collected from each injection group (ds*TRP*, ds*TRPR*, and ds*GFP*) for comparing *TRP* and *TRPR* expression. Individual brains were carefully dissected and frozen at -80°C until RNA extraction. Three independent replicate groups per condition were collected and qRT-PCR was performed to calculate the RNAi efficiency. Total RNA was isolated using TRIzol reagent (Takara, Japan). Total RNA quantification was performed by NanoDrop 2000 spectrophotometer (Thermo Fisher Scientific), and the quality of RNA was evaluated by 1.0% denaturing agarose gel electrophoresis. Reverse transcription was performed using a PrimeScript^TM^ RT reagent kit (Takara), according to the manufacturer’s instructions. Gene-specific mRNA levels were assessed by qPCR using TB Green Fast qPCR Mix (Takara) on a LightCycler 480II instrument (Roche, Switzerland). The *β-actin* gene was used as a reference gene. After verifying amplification efficiency of the selected genes and β-actin (from 96.8% to 100.5%), the differences in gene expression levels were calculated using the 2^−ΔΔCt^ method. Pairwise differences in gene expression were considered significant at p < 0.05, using one-way ANOVA (SPSS Statistics 20.0). The primers used for qPCR are shown in Table S11.

After determination of knockdown efficiencies (see results), 24 hours post-injection was chosen as the timepoint to study the PER effects of dsRNA-mediated knockdown of *TRP* and *TRPR*. About 200 bees of each behavioral phenotype (NBs, PFs, and NFs of AML) were collected in the morning, harnessed, and remained in a dark incubator to acclimatize. After 30 min, all surviving bees of each behavioral phenotype were evenly divided into three groups, injected with ds*TRP*, ds*TRPR*, and ds*GFP* and kept as described above. After 24 hours, all surviving bees were assayed for their PER to stimulations of sucrose solution, pollen, or larvae as described in section 4.2. Each stimulus was assessed with a new set of bees containing about 55 individuals for each treatment group (ds*TRP*, ds*TRPR*, and ds*GFP*). The SRSs of the *TPR*-knockdown*, TRPR*-knockdown, and control groups were compared using Kruskal-Wallis tests with Bonferroni correction for each behavioral phenotype separately. The sucrose responsiveness was further compared between the different groups at the same sucrose concentration with independent Chi-square tests. The responsiveness to pollen and larvae was compared between the *TPR*-knockdown*, TRPR*-knockdown, and control groups using independent Chi-square tests for each behavioral phenotype separately. All statistical analyses were performed with SPSS Statistics 20.0 (IBM).

### 4.7. Effects of TRP2 injection and RNAi-mediated downregulation of *TRP* and *TRPR* on ERK signaling in honeybee workers

To test whether manipulating TRP/TRPR signaling has effect on honeybee ERK signaling a group of 10 individual worker bees were collected from each injection group (ddH_2_O, TRP2, ds*TRP*, ds*TRPR*, and ds*GFP*) to compare ERK phosphorylation levels. Three independent replicate groups per condition were collected and Western blot analyses were performed: Honeybeebrains were carefully dissected and frozen at -80°C until protein extraction. Brain protein extractions were carried out according to our previously described method with some modifications. Briefly, the larvae were homogenized with lysis buffer (LB, 8 M urea, 2 M thiourea, 4% CHAPS, 20 mM Tris-base, 30 mM dithiothreitol). The mixture was homogenized for 30 min on ice and sonicated 20 s per 5 min during this time, then centrifuged at 12 000g and 4 °C for 10 min. Ice-cold acetone were added to the collected supernatants, and then the mixture was kept on ice for 30 min for protein precipitation. Subsequently, the mixture was centrifuged at 12 000g and 4 °C for 10 min. The supernatant was discarded and the pellets were resolved in LB and kept at -20°C for further use. Western blot analyses were performed as described in section 4.4.4.

## Data Availability

Original data have been deposited to ProteomeXchange Consortium with the dataset identifier PXD018713 under http://proteomecentral.proteomexchange.org. Other data not provided in the supplementary materials and materials are available from the first author upon request.

## Acknowledgments

We appreciate Dr. Huipeng Yang at Institute of Apicultural Research for kind gifts of vectors.

## Competing Interests

None of the authors have any competing interests.

## Supplementary Materials

**Fig S1.**
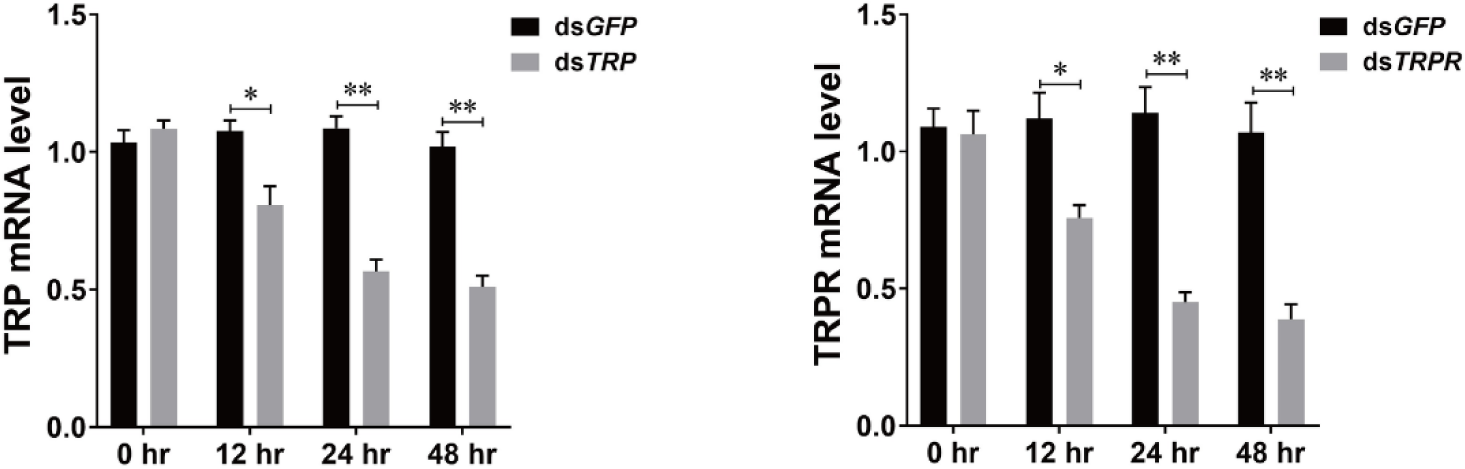
Efficiencies of dsRNA-mediated knockdown of *TRP* and *TRPR*. dsRNA (200 ng/bee for *TRP*, 2 μg/bee for *TRPR*) was microinjected into the head of honeybees via the central ocellus using a microinjector. dsRNA of green fluorescent protein gene (ds*GFP*, 2 μg/bee) was used as control. At 0, 12, 24, and 48 hours after injection, a group of 6 individual bees were collected from each injection group. Three independent replicate groups per condition were collected and qRT-PCR was performed to calculate the RNAi efficiency. Student’s t-tests were used for pairwise comparisons (*p<0.05, **p<0.01, ***p<0.001).

**Table S1.** The proboscis extension response of different behavioral phenotypes to different sucrose solutions. The proboscis extension response of *Apis mellifera ligustica* (AML) and *Apis cerana cerana* (ACC) worker bees to different sucrose solutions.

**AML pollen foragers**
| Concentration | Show PER | No PER | PER ratio |
| --- | --- | --- | --- |
| 0.1% | 48 | 79 | 37.80% |
| 0.3% | 51 | 76 | 40.16% |
| 1.0% | 70 | 57 | 55.12% |
| 3.0% | 83 | 44 | 65.35% |
| 10.0% | 87 | 40 | 68.50% |
| 30.0% | 111 | 16 | 87.40% |
| Pollen | 32 | 50 | 39.02% |
| Larva | 17 | 65 | 20.73% |

**ACC pollen foragers**
| Concentration | Show PER | No PER | PER ratio |
| --- | --- | --- | --- |
| 0.1% | 33 | 92 | 26.40% |
| 0.3% | 35 | 90 | 28.00% |
| 1.0% | 51 | 74 | 40.80% |
| 3.0% | 59 | 66 | 47.20% |
| 10.0% | 68 | 57 | 54.40% |
| 30.0% | 98 | 27 | 78.40% |
| Pollen | 20 | 66 | 23.26% |
| Larva | 11 | 75 | 12.79% |

**AML nectar foragers**
| Concentration | Show PER | No PER | PER ratio |
| --- | --- | --- | --- |
| 0.1% | 23 | 107 | 17.69% |
| 0.3% | 33 | 97 | 25.38% |
| 1.0% | 38 | 92 | 29.23% |
| 3.0% | 44 | 86 | 33.85% |
| 10.0% | 59 | 71 | 45.38% |
| 30.0% | 68 | 62 | 52.31% |
| Pollen | 11 | 74 | 12.94% |
| Larva | 15 | 70 | 17.65% |

**ACC nectar foragers**
| Concentration | Show PER | No PER | PER ratio |
| --- | --- | --- | --- |
| 0.1% | 17 | 111 | 13.28% |
| 0.3% | 19 | 109 | 14.84% |
| 1.0% | 23 | 105 | 17.97% |
| 3.0% | 29 | 99 | 22.66% |
| 10.0% | 44 | 84 | 34.38% |
| 30.0% | 55 | 73 | 42.97% |
| Pollen | 8 | 77 | 9.41% |
| Larva | 9 | 76 | 10.59% |

**AML nurse bees**
| **Concentration** | **Show PER** | **No PER** | **PER ratio** |
| --- | --- | --- | --- |
| **0.1%** | **30** | **106** | **22.06%** |
| **0.3%** | **32** | **104** | **23.53%** |
| **1.0%** | **45** | **91** | **33.09%** |
| **3.0%** | **50** | **86** | **36.76%** |
| **10.0%** | **57** | **79** | **41.91%** |
| **30.0%** | **75** | **61** | **55.15%** |
| **Pollen** | **9** | **82** | **9.89%** |
| **Larva** | **36** | **55** | **39.56%** |

**ACC nurse bees**
| **Concentration** | **Show PER** | **No PER** | **PER ratio** |
| --- | --- | --- | --- |
| **0.1%** | **18** | **113** | **13.74%** |
| **0.3%** | **19** | **112** | **14.50%** |
| **1.0%** | **30** | **101** | **22.90%** |
| **3.0%** | **38** | **93** | **29.01%** |
| **10.0%** | **48** | **83** | **36.64%** |
| **30.0%** | **58** | **73** | **44.27%** |
| **Pollen** | **7** | **81** | **7.95%** |
| **Larva** | **22** | **66** | **25.00%** |

**Table S2.** Statistical differences in sucrose responsiveness of different behavioral phenotypes.

| Concentration | 0.10% | 0.30% | 1.00% | 3.00% | 10.00% | 30.00% |
| --- | --- | --- | --- | --- | --- | --- |
| **AML** |  |  |  |  |  |  |
| PF vs NF | *** | * | *** | *** | *** | *** |
| PF vs NB | ** | ** | *** | *** | *** | *** |
| NF vs NB | ns | ns | ns | ns | ns | ns |
| **ACC** |  |  |  |  |  |  |
| PF vs NF | ** | ** | *** | *** | ** | *** |
| PF vs NB | * | ** | ** | ** | ** | *** |
| NF vs NB | ns | ns | ns | ns | ns | ns |
| **AML vs ACC** |  |  |  |  |  |  |
| PF | ns | * | * | ** | * | ns |
| NF | ns | * | * | * | ns | ns |
| NB | ns | ns | ns | ns | ns | ns |

AML: *Apis mellifera ligustica*, ACC: *Apis cerana cerana*, PF: pollen forager, NF: nectar forager, NB: nurse bee. ns = $P > 0.05$ , \* $P < 0.05$ , \*\* $P < 0.01$ , \*\*\* $P < 0.001$

**Table S3.** Neuropeptides identified in the brain of *Apis mellifera ligustica* workers. “NB” is nurse bee. “PF” is pollen forager. “NF” is nectar forager. “Protein Accession” is the unique number given to mark the entry of a protein in the database NCBInr. “Peptide” is the amino acid sequence of the peptide as determined in PEAKS Search. “-10lgP” is the score indicates the scoring significance of a peptide-spectrum match. “Mass” is monoisotopic mass of the peptide. “ppm” is precursor mass error, calculated as 10^6^ × (precursor mass - peptide mass) / peptide mass. “m/z” is precursor mass-to-charge ratio. “z” is peptide charge. “RT” is retention time (elution time) of the spectrum as recorded in the data. “#Spec” is the number of scanned spectrums of the peptide. “PTM” is post translational modification types present in the peptide.

| Sample | Protein Accession | Peptide | 10lgP | Mass | ppm | m/z | z | RT | #Spec | PTM |
| --- | --- | --- | --- | --- | --- | --- | --- | --- | --- | --- |
| NB | Q868G6.1 | NSIINDVKNELFPEDIN | 67.29 | 1972.974 | -0.3 | 987.494 | 2 | 98.51 | 10 |  |
| NB | Q868G6.1 | VLSMDGYQNILDKKDELLGEWE | 61.58 | 2594.257 | -7 | 1298.127 | 2 | 96.42 | 10 |  |
| NB | A8CL69.1 | pQLHNIVDKPRQN | 51.73 | 1443.758 | 0.7 | 482.2603 | 3 | 13.7 | 6 | Pyro-glu from Q |
| NB | A8CL69.1 | pQLHNIVDKPRQNFNDPRF | 51.12 | 2220.119 | 0.3 | 556.0372 | 4 | 41.34 | 6 | Pyro-glu from Q |
| NB | A8CL69.1 | TSQDITSGMWFGPRLa | 47.39 | 1693.825 | 0.1 | 847.9196 | 2 | 80.85 | 11 | Amidation |
| NB | A8CL69.1 | pQLHNIVDKP | 45.99 | 1045.556 | 0.8 | 523.7855 | 2 | 23.83 | 4 | Pyro-glu from Q |
| NB | A8CL69.1 | GMWFGPRLa | 33.41 | 961.4956 | 0 | 481.7551 | 2 | 68.13 | 9 | Amidation |
| NB | A8CL69.1 | RVPWTPSPRLa | 30.85 | 1206.699 | 0.3 | 604.3567 | 2 | 25.21 | 6 | Amidation |
| NB | A8CL69.1 | pQITQFTPRL | 27.13 | 1085.587 | -0.1 | 543.8007 | 2 | 78.09 | 3 | Pyro-glu from Q |
| NB | A8CL69.1 | MWFGPRLa | 26.77 | 904.4741 | -0.5 | 453.2441 | 2 | 71.2 | 5 | Amidation |
| NB | A8CL69.1 | QITQFTPRLa | 25.1 | 1101.63 | 0.5 | 551.8223 | 2 | 34.24 | 12 | Amidation |
| NB | A8CL69.1 | pQITQFTPRLa | 37.99 | 1084.603 | 0 | 543.3087 | 2 | 69.95 | 21 | Pyro-glu from Q;<br>Amidation |
| NB | ACI90290.1 | TWKSPDIVIRFa | 50.93 | 1359.766 | -0.3 | 454.2625 | 3 | 59.62 | 13 | Amidation |
| NB | ACI90290.1 | GRNDLNFIRYa | 48.35 | 1265.663 | -0.1 | 633.8386 | 2 | 33.11 | 11 | Amidation |
| NB | NP_001161192.1 | PEIFTSPEELRRYIDHVS DY YLLSGKARYa | 43.49 | 3515.784 | 0.4 | 586.9714 | 6 | 95.9 | 5 | Amidation |
| NB | P85527.1 | QDVDHVFLRFa | 55.21 | 1273.657 | 0 | 637.8356 | 2 | 50.66 | 9 | Amidation |
| NB | P85527.1 | pQDVDHVFLRFa | 53.95 | 1256.63 | -0.7 | 629.3219 | 2 | 74.39 | 15 | Pyro-glu from Q;<br>Amidation |
| NB | P85527.1 | pQDVDHVFLRF | 47.89 | 1257.614 | 0.8 | 629.8148 | 2 | 79.69 | 5 | Pyro-glu from Q |
| NB | P85527.1 | pQDVDHVFLR | 47.86 | 1110.546 | -0.5 | 556.2799 | 2 | 43.23 | 7 | Pyro-glu from Q |
| NB | P85527.1 | pQDVDHVFL | 28.42 | 954.4447 | 1.8 | 478.2305 | 2 | 69.45 | 5 | Pyro-glu from Q |
| NB | P85798.1 | LRNQLDIGDLQ | 50.23 | 1283.683 | -0.5 | 642.8486 | 2 | 42.48 | 10 |  |
| NB | P85798.1 | IPAADKERLLN | 47.66 | 1238.698 | 0.9 | 620.3569 | 2 | 15.09 | 6 |  |
| NB | P85798.1 | LRNQLDIGDL | 38.2 | 1155.625 | 0 | 578.8196 | 2 | 51.53 | 5 |  |
| NB | P85799.1 | SQAYDPYSNAAQFQLSSQSRGYPYQHRL<br>VY | 71.75 | 3523.655 | -0.6 | 881.9204 | 4 | 60.92 | 51 |  |
| NB | P85799.1 | SQAYDPYSNAAQFQLSSQSRGYPYQHRL<br>V | 70.34 | 3360.591 | -0.4 | 841.1547 | 4 | 57.58 | 11 |  |
| NB | P85799.1 | SQAYDPYSNAAQFQLSSQSRGYPYQHRL | 64.21 | 3261.523 | -1.2 | 816.387 | 4 | 53.22 | 8 |  |
| NB | P85799.1 | LPTNLAEDTKKTEQTMRPKS | 55.28 | 2287.184 | 0.1 | 572.8033 | 4 | 21.04 | 22 |  |
| NB | P85799.1 | GYPYQHRLVY | 49.18 | 1294.646 | 0.4 | 648.3304 | 2 | 25.22 | 15 |  |
| NB | P85799.1 | NVPIYQEPRF | 46 | 1261.646 | -0.3 | 631.8298 | 2 | 46.32 | 11 |  |
| NB | P85799.1 | VPIYQEPRF | 43.34 | 1147.603 | -0.1 | 574.8085 | 2 | 42.88 | 6 |  |
| NB | P85799.1 | PIYQEPRF | 28.55 | 1048.534 | 0.4 | 525.2746 | 2 | 46.03 | 7 |  |
| NB | P85828.1 | ITGQGNRIF | 46.68 | 1004.54 | -0.7 | 503.2771 | 2 | 19.63 | 8 |  |
| NB | P85828.1 | SLKAPFA | 41.78 | 732.417 | -0.6 | 367.2155 | 2 | 22.72 | 5 |  |
| NB | P85828.1 | SLKAPF | 36.02 | 661.3799 | 0.3 | 331.6973 | 2 | 23.77 | 4 |  |
| NB | P85829.1 | MVPVPVHHMADELLRNGPDTVI | 62.4 | 2439.24 | 0 | 1220.627 | 2 | 76.66 | 17 |  |
| NB | P85829.1 | VHHMADELLRNGPDTVI | 49.65 | 1915.957 | 0.1 | 639.6598 | 3 | 46.23 | 8 |  |
| NB | P85829.1 | VPVPVHHMADELL | 43.49 | 1455.754 | 1.3 | 728.8854 | 2 | 47.73 | 7 |  |
| NB | P85829.1 | LLRNGPDTVI | 35 | 1096.624 | 0.2 | 549.3194 | 2 | 28.64 | 5 |  |
| NB | P85829.1 | LRNGPDTVI | 22.27 | 983.54 | 0.5 | 492.7775 | 2 | 16.98 | 6 |  |
| NB | P85830.1 | GLDLGLSRGFSGSQAAKHLMLGAAANYA<br>GGPa | 70.62 | 2985.524 | -0.9 | 996.1811 | 3 | 84.68 | 9 | Amidation |
| NB | P85830.1 | GLDLGLSRGFSGSQAA | 62.96 | 1534.774 | 1.2 | 768.3951 | 2 | 55.2 | 6 |  |
| NB | P85830.1 | GLDLGLSRGFSGSQAAKH | 53.33 | 1799.928 | -0.5 | 600.9829 | 3 | 31.47 | 10 |  |
| NB | P85830.1 | GLDLGLSRGFSGSQAAKHLMa | 46.31 | 2043.068 | 1 | 682.0308 | 3 | 57.54 | 8 | Amidation |
| NB | P85831.1 | IDLSRFYGHFNT | 60.52 | 1468.71 | -0.3 | 735.362 | 2 | 64.92 | 29 |  |
| NB | P85831.1 | IDLSRFYGHFN | 56.71 | 1367.662 | -0.1 | 684.8383 | 2 | 62.93 | 18 |  |
| NB | P85831.1 | IDLSRFYGHF | 52.75 | 1253.619 | -0.6 | 627.8165 | 2 | 70.59 | 19 |  |
| NB | P85831.1 | IDLSRFYGHFNTKR | 48.89 | 1752.906 | 0 | 439.2337 | 4 | 43.28 | 23 |  |
| NB | P85831.1 | FYGHFNT | 44.93 | 884.3817 | -0.2 | 443.1981 | 2 | 21.44 | 7 |  |
| NB | P85831.1 | DLSRFYGHF | 25.85 | 1140.535 | 0.2 | 571.275 | 2 | 70.16 | 3 |  |
| NB | P85831.1 | DLSRFYGHFN | 20.23 | 1254.578 | 0.4 | 628.2966 | 2 | 62.68 | 22 |  |
| NB | P85832.1 | LTNYLATTGHGTNTGGPVLT | 82.04 | 1987.001 | -1.4 | 994.5065 | 2 | 47.57 | 22 |  |
| NB | P85832.1 | LTNYLATTGHGTNTGGPVL | 69.52 | 1885.953 | -0.6 | 943.9834 | 2 | 52.3 | 4 |  |
| NB | P85832.1 | NLDEIDRVGWSGFV | 62.73 | 1605.779 | 0.3 | 803.8969 | 2 | 88.41 | 3 |  |
| NB | P85832.1 | LTNYLATTGHGTNTGGPVLTRRFa | 49.49 | 2445.288 | -0.4 | 816.1028 | 3 | 39.57 | 13 | Amidation |
| NB | P85832.1 | NIDEIDRTAFDNFF | 46.68 | 1715.779 | -1.2 | 858.8958 | 2 | 96.46 | 9 |  |
| NB | P85832.1 | LVDELSPVSERETLEFa | 33.35 | 2017.048 | 0.3 | 673.3568 | 3 | 63.4 | 7 | Amidation |
| NB | P85832.1 | ELVDELSPVSERETLEFa | 30.33 | 2146.091 | 0.6 | 716.3712 | 3 | 74.41 | 9 | Amidation |
| NB | Q06601.1 | GNNRPVYIPQRPHPRL | 33.24 | 2107.155 | 0.3 | 422.4384 | 5 | 25.01 | 15 |  |
| NB | Q06601.1 | VYIPQRPHPRL | 23.32 | 1568.894 | -0.2 | 393.2307 | 4 | 29.92 | 20 |  |
| NB | Q06601.1 | AVHYSGGQPLGSKRPNDMLSQRYHFGLa | 65.31 | 3013.509 | -1.6 | 754.3834 | 4 | 38.67 | 9 | Amidation |
| NB | Q06601.1 | PNDMLSQRYHFGLa | 66.71 | 1575.762 | 0.3 | 526.2613 | 3 | 56.93 | 13 | Amidation |
| NB | Q06601.1 | AYTYVSEYKRLPVYNFGLa | 29.98 | 2181.126 | -0.2 | 728.0491 | 3 | 72.4 | 8 | Amidation |
| NB | Q06601.1 | ADYPLRLNLD | 48.67 | 1188.614 | 0 | 595.3142 | 2 | 56.85 | 11 |  |
| NB | Q06601.1 | YPLRLNLD | 43.48 | 1002.55 | 0.6 | 502.2825 | 2 | 49.18 | 8 |  |
| NB | Q06601.1 | RQYSFGLa | 31.09 | 868.4555 | 0 | 435.235 | 2 | 26.91 | 10 | Amidation |
| NB | Q06601.1 | GRQPYSFGLa | 35.27 | 1022.53 | -0.1 | 512.2721 | 2 | 32.39 | 6 | Amidation |
| NB | Q06601.1 | GRDYSFGLa | 31.03 | 912.4453 | 0 | 457.2299 | 2 | 30.51 | 3 | Amidation |
| NB | Q06601.1 | WIDTNDNKRGRDYSFGLa | 29.02 | 2054.992 | 0 | 686.0047 | 3 | 38.47 | 7 | Amidation |
| NB | Q06601.1 | LDYLPVDNPAFH | 51.58 | 1399.677 | 0.6 | 700.8463 | 8 | 65.51 | 4 |  |
| NB | Q06602.1 | EAEPEAEPGNNRPVYIPQPRPPHRL | 50.05 | 2959.505 | -0.5 | 592.908 | 5 | 33.75 | 26 |  |
| NB | Q06602.1 | GNNRPVYIPQPRPPHRL | 33.24 | 2107.155 | 0.3 | 422.4384 | 5 | 25.01 | 15 |  |
| NB | Q06602.1 | VYIPQPRPPHRL | 23.32 | 1568.894 | -0.2 | 393.2307 | 4 | 29.92 | 20 |  |
| NB | Q5DW47.1 | STSLEELANR | 39.7 | 1118.557 | 0.9 | 560.2861 | 2 | 24.18 | 4 |  |
| NB | Q5DW47.1 | STSLEELANRN | 38.16 | 1232.6 | 0.7 | 617.3075 | 2 | 23.07 | 5 |  |
| NB | Q5DW47.1 | pQTFTYSHGWTNa | 18.99 | 1322.568 | -0.1 | 662.2912 | 2 | 51.14 | 10 | Pyro-glu from Q;<br>Amidation |
| NB | Q868G6.1 | ASFDDEYYKRAPMGFQGMRa | 55.4 | 2267.025 | 0.5 | 567.7639 | 4 | 45.63 | 9 | Amidation |
| NB | Q868G6.1 | APMGFQGMRG | 50.96 | 1050.474 | 0 | 526.2442 | 2 | 22.9 | 6 |  |
| NB | Q868G6.1 | GVMDFQIGLQ | 50.62 | 1106.543 | 1 | 554.2793 | 2 | 85.81 | 6 |  |
| NB | Q868G6.1 | APMGFQGMRa | 49.03 | 992.4684 | -1.1 | 497.241 | 2 | 18.87 | 16 | Amidation |
| NB | Q868G6.1 | VLSMDGYQNILD | 47.52 | 1366.644 | 0.6 | 684.3296 | 2 | 80.98 | 15 |  |
| NB | Q868G6.1 | NPRWEFRGKFVGVRa | 47.04 | 1745.959 | -0.2 | 437.4969 | 4 | 24.64 | 9 | Amidation |
| NB | Q868G6.1 | ARMGFHGMRa | 46.29 | 1060.517 | -0.4 | 354.5128 | 3 | 8.86 | 3 | Amidation |
| NB | Q868G6.1 | ALMGFQGVRG | 46.07 | 1034.533 | 0.1 | 518.2739 | 2 | 35.2 | 6 |  |
| NB | Q868G6.1 | SPFRYLGA | 45.4 | 909.4708 | 0 | 455.7427 | 2 | 36.86 | 10 |  |
| NB | Q868G6.1 | APMGFYGTRa | 45.2 | 997.4803 | -0.1 | 499.7474 | 2 | 16.94 | 3 | Amidation |
| NB | Q868G6.1 | APMGFYGTRG | 45.18 | 1055.486 | 0.4 | 528.7504 | 2 | 20.63 | 7 |  |
| NB | Q868G6.1 | ALMGFQGVRa | 44.3 | 976.5276 | -0.5 | 489.2709 | 2 | 29.63 | 13 | Amidation |
| NB | Q868G6.1 | SPFRYLARG | 44.18 | 1122.593 | -0.4 | 375.2049 | 3 | 20.59 | 11 |  |
| NB | Q868G6.1 | GVMDFQIGLQRKKD | 44.03 | 1633.861 | -0.2 | 817.9376 | 2 | 35.7 | 14 |  |
| NB | Q868G6.1 | SPFRYLGARa | 43.32 | 1064.588 | 0.2 | 355.87 | 3 | 16.59 | 8 | Amidation |
| NB | Q868G6.1 | NPRWEFRGKFVGV | 42.84 | 1590.842 | 0.1 | 531.288 | 3 | 43.57 | 15 |  |
| NB | Q868G6.1 | SPFRYLG | 37.59 | 838.4337 | 0 | 420.2241 | 2 | 31.97 | 7 |  |
| NB | Q868G6.1 | SLEEILDEIK | 33.02 | 1187.629 | 0 | 594.8215 | 2 | 88.28 | 6 |  |
| NB | Q868G6.1 | SLEEILDEI | 29.37 | 1059.534 | 0.1 | 530.7741 | 2 | 108.02 | 4 |  |
| NB | Q868G6.1 | ASFDDEYY | 28.99 | 1008.371 | 0 | 505.1929 | 2 | 43.35 | 4 |  |
| NB | XP_006557714.1 | pQQFDDYGHLRFa | 47.97 | 1406.637 | -2.3 | 704.324 | 2 | 68.3 | 4 | Pyro-glu from Q;<br>Amidation |
| NB | XP_006559359.1 | NVASLARTYTLPQNAa | 64.35 | 1616.863 | -1.3 | 809.4379 | 2 | 43.18 | 6 | Amidation |
| NB | XP_006559359.1 | SVSSLAKNSAWPVSL | 62.69 | 1544.82 | -1.3 | 773.4162 | 2 | 68.52 | 8 |  |
| NB | XP_006559359.1 | FLLLPATDNNYFHQKLPSSLRKSL | 56.55 | 2888.555 | 1 | 578.7188 | 5 | 71.13 | 15 |  |
| NB | XP_006559359.1 | NVGSVAREHGLPYa | 55.04 | 1396.721 | -0.8 | 699.3672 | 2 | 21.03 | 15 | Amidation |
| NB | XP_006559359.1 | SVSSLARTGDLPVREQ | 53.68 | 1713.901 | 0.5 | 572.3079 | 3 | 25.97 | 12 |  |
| NB | XP_006559359.1 | YVASLARTGDLPIRGQ | 51.94 | 1715.932 | 0.4 | 572.9847 | 3 | 35.62 | 12 |  |
| NB | XP_006559359.1 | NIASLMRDYDQSRENRVPFPa | 47.38 | 2406.186 | 0.1 | 803.0695 | 3 | 63.84 | 12 | Amidation |
| NB | XP_006559359.1 | HIGALARLGWLP SLRTA | 42.32 | 1831.058 | -0.2 | 611.3598 | 3 | 70.88 | 7 |  |
| NB | XP_006559359.1 | HIGALARLGWLP SLRTARFS | 42 | 2221.26 | -0.4 | 556.322 | 4 | 71.36 | 9 |  |
| NB | XP_006559359.1 | NVGTLARDFALPPa | 40.53 | 1368.751 | -0.1 | 685.3829 | 2 | 60.79 | 16 | Amidation |
| NB | XP_006559359.1 | YVASLARTGDLPIRa | 40.29 | 1529.868 | 0.6 | 510.9635 | 3 | 34.02 | 8 | Amidation |
| NB | XP_006559359.1 | GIFLPGSVILR | 37.83 | 1170.712 | -0.1 | 586.3634 | 2 | 77.93 | 5 |  |
| NB | XP_006559359.1 | LPGSVILRALS | 36.35 | 1124.692 | 2.1 | 563.3543 | 2 | 72.8 | 8 |  |
| NB | XP_006559359.1 | GIFLPGSVILRALS RQa | 36.3 | 1725.041 | -0.9 | 576.0205 | 3 | 95.14 | 10 | Amidation |
| NB | XP_006559359.1 | NVGTLARDFALPPGRRNIASLMRDYDQSR<br>ENRVFPa | 21.57 | 4127.135 | 0.2 | 688.8632 | 6 | 75.08 | 9 | Amidation |
| NB | XP_006559865.1 | AFGLLTYPRIa | 40.74 | 1148.671 | 0.5 | 575.3428 | 2 | 70.98 | 6 | Amidation |
| NB | XP_006559865.1 | SNAPISNLNFN | 35.48 | 1189.573 | 0.3 | 595.7938 | 2 | 48.7 | 4 |  |
| NB | XP_006559865.1 | EKLKPNMRRAFGLLTYPRIa | 28.33 | 2301.325 | 0.6 | 576.339 | 4 | 50.2 | 8 | Amidation |
| NB | XP_006560385.1 | AYRKPPFNGSIFa | 42.26 | 1394.746 | -0.5 | 698.3799 | 2 | 36.92 | 12 | Amidation |
| NB | XP_006560385.1 | KPPFNGSIFa | 39.41 | 1004.544 | 0.2 | 503.2795 | 2 | 43.74 | 6 | Amidation |
| NB | XP_006560385.1 | RKPPFNGSIFa | 32.35 | 1160.645 | 0 | 581.33 | 2 | 28.38 | 7 | Amidation |
| NB | XP_006560385.1 | YRKPPFNGSIFa | 25.22 | 1323.709 | 0.5 | 662.862 | 2 | 36.42 | 6 | Amidation |
| NB | XP_006562922.1 | GFKPEYISTAYGFa | 40.22 | 1477.724 | 0.2 | 739.8695 | 2 | 64.18 | 4 | Amidation |
| NB | XP_006565207.1 | SDPHLSILSKPMSAIPSYKFDD | 81.44 | 2447.204 | 0.4 | 816.7423 | 3 | 71.96 | 17 |  |
| NB | XP_006565207.1 | SPSLRLRFa | 40.12 | 973.5821 | 0.2 | 487.7984 | 2 | 24.53 | 13 | Amidation |
| NB | XP_006565207.1 | SDPHLSILS | 39.05 | 967.4974 | 0.4 | 484.7562 | 2 | 34.74 | 7 |  |
| NB | XP_006565207.1 | SQRSPSLRLRFa | 38.06 | 1344.774 | 0.4 | 449.2654 | 3 | 16.83 | 10 | Amidation |
| NB | XP_006565207.1 | SDPHLSILSKPMSAIP | 32.57 | 1691.892 | -1.1 | 846.9521 | 2 | 64.26 | 4 |  |
| NB | XP_006570344.1 | NSELINSLGLPKNMNNAa | 65.94 | 1940.015 | 0.5 | 971.0152 | 2 | 87.45 | 11 | Amidation |
| NB | XP_006570344.1 | LINSLGLPKNMNNAa | 35.9 | 1609.897 | 1.1 | 805.9568 | 2 | 62.46 | 6 | Amidation |
| NB | XP_016769998.1 | LVDHRIPDLENEMFDSGNDPGSTVVRT | 78.07 | 3012.425 | 0.1 | 754.1135 | 4 | 63.86 | 16 |  |
| NB | XP_016769998.1 | HPISYNTYDERELSRDHPPLLL | 30.5 | 2664.33 | -1.6 | 667.0886 | 4 | 54.55 | 3 |  |
| NB | XP_016769998.1 | IGSLSIVNSMDVLRQRVLLELARRKALQD<br>QAQIDANRRLLLETla | 27.71 | 4913.782 | -0.3 | 819.9707 | 6 | 98.39 | 12 | Amidation |
| PF | Q868G6.1 | NSIINDVKNELFPEDIN | 48.21 | 1972.974 | -0.3 | 987.494 | 2 | 100.35 | 14 |  |
| PF | A8CL69.1 | TSQDITSGMWFGPRLa | 42.05 | 1693.825 | 0.5 | 847.92 | 2 | 79 | 22 | Amidation |
| PF | A8CL69.1 | pQLHNIVDKPRQN | 37.86 | 1443.758 | 0.3 | 482.2602 | 3 | 14.35 | 3 | Pyro-glu from Q |
| PF | A8CL69.1 | pQLHNIVDKP | 36.78 | 1045.556 | 0 | 523.7851 | 2 | 25.52 | 6 | Pyro-glu from Q |
| PF | A8CL69.1 | pQITQFTPRLa | 29.72 | 1084.603 | 0.4 | 543.309 | 2 | 68.45 | 3 | Pyro-glu from Q;<br>Amidation |
| PF | A8CL69.1 | RVPWTPSPRLa | 25.44 | 1206.699 | 1.6 | 604.3575 | 2 | 28.86 | 5 | Amidation |
| PF | A8CL69.1 | pQLHNIVDKPRQNFNDPRF | 23.63 | 2220.119 | -0.9 | 556.0365 | 4 | 47.19 | 4 | Pyro-glu from Q |
| PF | A8CL69.1 | QITQFTPRLa | 22.79 | 1101.63 | -0.5 | 551.8218 | 2 | 37.4 | 7 | Amidation |
| PF | A8CL69.1 | MWFGPRLa | 20.27 | 904.4741 | -0.2 | 453.2443 | 2 | 75.27 | 10 | Amidation |
| PF | A8CL69.1 | GMWFGPRLa | 17.98 | 961.4956 | 0.7 | 481.7554 | 2 | 72.51 | 12 | Amidation |
| PF | ACI90290.1 | TWKSPDIVIRFa | 42 | 1359.766 | 0.1 | 454.2627 | 3 | 63.04 | 13 | Amidation |
| PF | ACI90290.1 | GRNDLNFIRYa | 36.79 | 1265.663 | 0.2 | 633.8388 | 2 | 37.18 | 24 | Amidation |
| PF | ACI90290.1 | AGFKNLNREQ | 35.46 | 1175.605 | 0 | 392.8755 | 3 | 10.1 | 6 |  |
| PF | ACI90290.1 | SPDIVIRFa | 28.69 | 944.5443 | -1.1 | 473.2789 | 2 | 51.32 | 7 | Amidation |
| PF | NP_001161192.1 | PEIFTSPEELRRYIDHVSDYYLLSGKARYa | 45.15 | 3515.784 | 0.4 | 586.9714 | 6 | 95.9 | 8 | Amidation |
| PF | P85527.1 | pQDVDHVFLRFa | 43.1 | 1256.63 | -0.6 | 629.322 | 2 | 76.1 | 19 | Pyro-glu from Q;<br>Amidation |
| PF | P85527.1 | pQDVDHVFLR | 40.39 | 1110.546 | 0.1 | 556.2802 | 2 | 44.76 | 7 | Pyro-glu from Q |
| PF | P85527.1 | QDVDHVFLRFa | 40.1 | 1273.657 | 0.1 | 637.8357 | 2 | 54.09 | 9 | Amidation |
| PF | P85527.1 | pQDVDHVFLRF | 38.93 | 1257.614 | 0 | 629.8143 | 2 | 82.73 | 8 | Pyro-glu from Q |
| PF | P85527.1 | pQDVDHVFL | 27.57 | 954.4447 | 0.2 | 478.2297 | 2 | 70.88 | 3 | Pyro-glu from Q |
| PF | P85798.1 | LRNQLDIGDLQ | 38.97 | 1283.683 | -1 | 642.8483 | 2 | 44.49 | 12 |  |
| PF | P85798.1 | IPAADKERLLN | 33.42 | 1238.698 | 1.2 | 413.9072 | 3 | 15.96 | 6 |  |
| PF | P85798.1 | LRNQLDIGDL | 31.19 | 1155.625 | 0.3 | 578.8198 | 2 | 54.4 | 11 |  |
| PF | P85799.1 | SQAYDPYSNAAQFQLSSQSRGYPYQHRL<br>VY | 51.05 | 3523.655 | -0.1 | 881.9208 | 4 | 64.95 | 41 |  |
| PF | P85799.1 | SQAYDPYSNAAQFQLSSQSRGYPYQHRL<br>V | 42.31 | 3360.591 | 0.4 | 841.1554 | 4 | 59.88 | 14 |  |
| PF | P85799.1 | LPTNLAEDTKKTEQTMRPKS | 37.87 | 2287.184 | 0.3 | 572.8035 | 4 | 24.72 | 28 |  |
| PF | P85799.1 | SQAYDPYSNAAQFQLSSQSRGYPYQHRL | 37.71 | 3261.523 | -3.4 | 816.3852 | 4 | 56.14 | 10 |  |
| PF | P85799.1 | GYPYQHRLVY | 37.44 | 1294.646 | 0.3 | 648.3304 | 2 | 29.02 | 28 |  |
| PF | P85799.1 | NVPIYQEPRF | 37.08 | 1261.646 | -1.1 | 631.8293 | 2 | 48.46 | 16 |  |
| PF | P85799.1 | VPIYQEPRF | 35.84 | 1147.603 | -0.1 | 574.8085 | 2 | 45.61 | 13 |  |
| PF | P85799.1 | PIYQEPRF | 25.33 | 1048.534 | 0.1 | 525.2744 | 2 | 49.81 | 6 |  |
| PF | P85828.1 | ITGQGNRIF | 37.41 | 1004.54 | -0.8 | 503.277 | 2 | 21.59 | 21 |  |
| PF | P85828.1 | SLKAPFA | 34.04 | 732.417 | 0.1 | 367.2158 | 2 | 24.36 | 11 |  |
| PF | P85828.1 | SLKAPF | 29.98 | 661.3799 | 0.1 | 331.6973 | 2 | 26.29 | 15 |  |
| PF | P85829.1 | MVPVPVHHMADELLRNGPDTVI | 36.48 | 2439.24 | 0.7 | 814.0879 | 3 | 81.53 | 22 |  |
| PF | P85829.1 | VHHMADELLRNGPDTVI | 33.81 | 1915.957 | 1.1 | 639.6605 | 3 | 51.24 | 10 |  |
| PF | P85829.1 | LLRNGPDTVI | 31.63 | 1096.624 | 0.4 | 549.3195 | 2 | 30.43 | 6 |  |
| PF | P85829.1 | LRNGPDTVI | 28.06 | 983.54 | 0.1 | 492.7773 | 2 | 17.56 | 6 |  |
| PF | P85829.1 | VPVPVHHMADELL | 20.81 | 1455.754 | 0.4 | 486.2589 | 3 | 51.21 | 6 |  |
| PF | P85830.1 | GLDLGLSRGFSGSQAAKHLMGLAAANYA<br>GGPa | 46.77 | 2985.524 | -0.3 | 747.3881 | 4 | 90.26 | 20 | Amidation |
| PF | P85830.1 | GLDLGLSRGFSGSQAA | 43.99 | 1534.774 | 0.5 | 768.3947 | 2 | 57.69 | 5 |  |
| PF | P85830.1 | HLMGLAAANYAGGPa | 32.71 | 1340.666 | -0.7 | 671.3398 | 2 | 41.33 | 7 | Amidation |
| PF | P85830.1 | GLDLGLSRGFSGSQAAKHLMa | 31.39 | 2043.068 | 0.5 | 682.0304 | 3 | 61.51 | 6 | Amidation |
| PF | P85830.1 | GLDLGLSRGFSGSQAACH | 28.75 | 1799.928 | 0.1 | 600.9833 | 3 | 34.79 | 8 |  |
| PF | P85831.1 | IDLSRFYGHFNT | 45.95 | 1468.71 | -0.5 | 735.3618 | 2 | 69.86 | 42 |  |
| PF | P85831.1 | IDLSRFYGHFN | 42.44 | 1367.662 | -1.9 | 684.8371 | 2 | 65.48 | 27 |  |
| PF | P85831.1 | IDLSRFYGHF | 41.39 | 1253.619 | -0.6 | 627.8165 | 2 | 74.6 | 23 |  |
| PF | P85831.1 | FYGHFNT | 36.1 | 884.3817 | 0.6 | 443.1984 | 2 | 23.94 | 9 |  |
| PF | P85831.1 | DLSRFYGHFN | 34.65 | 1254.578 | 0.1 | 628.2964 | 2 | 65.54 | 4 |  |
| PF | P85831.1 | IDLSRFYGHFNTR | 32.23 | 1752.906 | -0.1 | 439.2337 | 4 | 49.93 | 10 |  |
| PF | P85832.1 | LTNYLATTGHGTNTGGPVL | 52.74 | 1987.001 | 0.2 | 994.5081 | 2 | 50.02 | 12 |  |
| PF | P85832.1 | NLDEIDRVGWSGFV | 47.92 | 1605.779 | 0 | 803.8966 | 2 | 93.23 | 14 |  |
| PF | P85832.1 | LTNYLATTGHGTNTGGPVL | 45.56 | 1885.953 | 0.3 | 943.9843 | 2 | 54.77 | 5 |  |
| PF | P85832.1 | NIDEIDRTAFDNFF | 43.44 | 1715.779 | 1.3 | 858.8979 | 2 | 98.86 | 7 |  |
| PF | P85832.1 | LTNYLATTGHGTNTGGPVLTRRFa | 37.72 | 2445.288 | 0.5 | 612.3295 | 4 | 44.18 | 11 | Amidation |
| PF | P85832.1 | ELVDELSPVSERETLERFa | 32.49 | 2146.091 | 0.9 | 716.3715 | 3 | 77.64 | 9 | Amidation |
| PF | P85832.1 | LVDELSPVSERETLERFa | 31.14 | 2017.048 | -0.5 | 673.3563 | 3 | 66.54 | 10 | Amidation |
| PF | Q06601.1 | AVHYSGGQPLGSKRPNDMLSQRYHFGLa | 60.6 | 3013.509 | 0.6 | 754.3851 | 4 | 40.53 | 8 | Amidation |
| PF | Q06601.1 | PNDMLSQRYHFGLa | 65.73 | 1575.762 | 0.1 | 788.8882 | 2 | 47.77 | 9 | Amidation |
| PF | Q06601.1 | AYTYVSEYKRLPVYNFGla | 30.64 | 2181.126 | 0.3 | 728.0494 | 3 | 72.56 | 3 | Amidation |
| PF | Q06601.1 | ADYPLRLNLD | 46.77 | 1188.614 | 0 | 595.3142 | 2 | 56.69 | 6 |  |
| PF | Q06601.1 | YPLRLNLD | 42.12 | 1002.55 | 0.3 | 502.2823 | 2 | 49.35 | 12 |  |
| PF | Q06601.1 | RQYSFGLa | 30.29 | 868.4555 | -0.2 | 435.235 | 2 | 26.45 | 19 | Amidation |
| PF | Q06601.1 | GRQPYSFGLa | 34.37 | 1022.53 | 0.3 | 512.2723 | 2 | 31.51 | 5 | Amidation |
| PF | Q06601.1 | GRDYSFGLa | 28.95 | 912.4453 | 0.2 | 457.23 | 2 | 30.84 | 8 | Amidation |
| PF | Q06601.1 | WIDTNDNKRGRDYSFGLa | 24.14 | 2054.992 | 0.4 | 686.0049 | 3 | 38.84 | 3 | Amidation |
| PF | Q06601.1 | LDYLPVDNPAFH | 40.17 | 1399.677 | -0.4 | 700.8456 | 2 | 67.71 | 7 |  |
| PF | Q06601.1 | AVHYSGGQPLGS | 39.1 | 1171.562 | 0.2 | 586.7885 | 2 | 14.58 | 6 |  |
| PF | Q06602.1 | EAEPEAEPGNRPVYIPQPRPPHRL | 23.07 | 2959.505 | 0.1 | 592.9084 | 5 | 39.74 | 5 |  |
| PF | Q5DW47.1 | STSLEELANR | 28.15 | 1118.557 | 0.3 | 560.2858 | 2 | 25.81 | 5 |  |
| PF | Q5DW47.1 | STSLEELANRN | 26.91 | 1232.6 | -0.5 | 617.3068 | 2 | 24.93 | 3 |  |
| PF | Q5DW47.1 | pQTFTYSHGWTNa | 22.74 | 1322.568 | 0.6 | 662.2916 | 2 | 52.52 | 6 | Pyro-glu from Q;<br>Amidation |
| PF | Q868G6.1 | ASFDDEYYKRAPMGFQGMRa | 42.06 | 2267.025 | 0.6 | 567.7639 | 4 | 49.19 | 22 | Amidation |
| PF | Q868G6.1 | VLSMDGYQNILDKKDELLGEWE | 41.48 | 2594.257 | -0.4 | 865.7594 | 3 | 98.46 | 12 |  |
| PF | Q868G6.1 | VLSMDGYQNILD | 40.43 | 1366.644 | -0.3 | 684.329 | 2 | 82.61 | 11 |  |
| PF | Q868G6.1 | GVMDFQIGLQ | 40.09 | 1106.543 | -0.7 | 554.2784 | 2 | 87.57 | 16 |  |
| PF | Q868G6.1 | APMGFQGMRG | 38.9 | 1050.474 | -0.5 | 526.244 | 2 | 24.95 | 13 |  |
| PF | Q868G6.1 | APMGFQGMRa | 38.73 | 992.4684 | -0.7 | 497.2411 | 2 | 20.02 | 38 | Amidation |
| PF | Q868G6.1 | APMGFYGTRG | 38.3 | 1055.486 | -0.9 | 528.7497 | 2 | 21.9 | 13 |  |
| PF | Q868G6.1 | ARMGFHGMRa | 37.1 | 1060.517 | 0 | 531.2658 | 2 | 9 | 18 | Amidation |
| PF | Q868G6.1 | APMGFYGTRa | 37.08 | 997.4803 | -1.1 | 499.7469 | 2 | 18.43 | 23 | Amidation |
| PF | Q868G6.1 | ALMGFQGVRa | 36.88 | 976.5276 | -0.7 | 489.2708 | 2 | 31.91 | 23 | Amidation |
| PF | Q868G6.1 | ALMGFQGVRG | 36.33 | 1034.533 | 0.2 | 518.2739 | 2 | 38.06 | 10 |  |
| PF | Q868G6.1 | SPFRYLGA | 35.81 | 909.4708 | -0.3 | 455.7426 | 2 | 40.27 | 14 |  |
| PF | Q868G6.1 | GVMDFQIGLQRKKD | 34.42 | 1633.861 | 0.6 | 545.6279 | 3 | 39.49 | 8 |  |
| PF | Q868G6.1 | SPFRYLGARG | 34.3 | 1122.593 | -0.5 | 375.2049 | 3 | 23.99 | 7 |  |
| PF | Q868G6.1 | SPFRYLGARa | 33.85 | 1064.588 | 0 | 533.3012 | 2 | 19.83 | 7 | Amidation |
| PF | Q868G6.1 | SLEEILDEIK | 33.49 | 1187.629 | 0.1 | 594.8216 | 2 | 93.96 | 13 |  |
| PF | Q868G6.1 | SPFRYLG | 30.94 | 838.4337 | 0.3 | 420.2242 | 2 | 36.16 | 10 |  |
| PF | Q868G6.1 | NPRWEFRGKFVGVRa | 30.44 | 1745.959 | 0.5 | 437.4972 | 4 | 32.34 | 10 | Amidation |
| PF | Q868G6.1 | NPRWEFRGKFVGV | 30.3 | 1590.842 | 0.3 | 531.2881 | 3 | 49.83 | 3 |  |
| PF | Q868G6.1 | ASFDDEYY | 28.5 | 1008.371 | 0.1 | 505.1929 | 2 | 44.7 | 5 |  |
| PF | Q868G6.1 | SLEEILDEI | 25.89 | 1059.534 | 0.4 | 530.7743 | 2 | 108.72 | 6 |  |
| PF | Q868G6.1 | IILDALEELD | 25.61 | 1142.607 | -0.2 | 572.3107 | 2 | 100.26 | 3 |  |
| PF | XP_006557714.1 | pQQFDDYGHLRFa | 41.67 | 1406.637 | 0.5 | 704.326 | 2 | 69.79 | 13 | Pyro-glu from Q;<br>Amidation |
| PF | XP_006559359.1 | SVSSLAKNSAWPVSL | 46.58 | 1544.82 | -0.3 | 773.417 | 2 | 71.29 | 11 |  |
| PF | XP_006559359.1 | NVASLARTYTLPQNAa | 44.1 | 1616.863 | -0.5 | 809.4386 | 2 | 46.96 | 8 | Amidation |
| PF | XP_006559359.1 | FLLLPATDNNYFHQKLPSLRKSL | 42.63 | 2888.555 | 0.4 | 578.7184 | 5 | 77.31 | 22 |  |
| PF | XP_006559359.1 | NVGSVAREHGLPYa | 41.93 | 1396.721 | -0.5 | 699.3674 | 2 | 24.98 | 21 | Amidation |
| PF | XP_006559359.1 | YVASLARTGDLPIRGQ | 40.96 | 1715.932 | 0 | 572.9846 | 3 | 37.62 | 10 |  |
| PF | XP_006559359.1 | HIGALARLGWLPSLRTA | 40.77 | 1831.058 | -0.1 | 611.3599 | 3 | 78.84 | 14 |  |
| PF | XP_006559359.1 | NIASLMRDYDQSRENRPFPa | 39.13 | 2406.186 | 0.9 | 803.0701 | 3 | 69.67 | 13 | Amidation |
| PF | XP_006559359.1 | SVSSLARTGDLPVREQ | 38.63 | 1713.901 | -0.4 | 572.3073 | 3 | 27.58 | 8 |  |
| PF | XP_006559359.1 | NVGTLARDFALPPa | 38.03 | 1368.751 | -0.6 | 685.3825 | 2 | 63.59 | 18 | Amidation |
| PF | XP_006559359.1 | GIFLPGSVILRALSRQa | 37.1 | 1725.041 | 0 | 576.0211 | 3 | 98.8 | 12 | Amidation |
| PF | XP_006559359.1 | YVASLARTGDLPIRa | 33.92 | 1529.868 | 0 | 510.9632 | 3 | 36.19 | 7 | Amidation |
| PF | XP_006559359.1 | LPGSVILRALS | 28.72 | 1124.692 | 0.4 | 563.3533 | 2 | 76.05 | 10 |  |
| PF | XP_006559359.1 | GIFLPGSVILR | 27.16 | 1170.712 | 1.5 | 586.3644 | 2 | 80.54 | 14 |  |
| PF | XP_006559359.1 | HIGALARLGWLPSLRTARFS | 25.34 | 2221.26 | 0.4 | 556.3224 | 4 | 81.47 | 4 |  |
| PF | XP_006559865.1 | AFGLLTYPRIa | 34.68 | 1148.671 | -0.1 | 575.3425 | 2 | 73.38 | 13 | Amidation |
| PF | XP_006560385.1 | AYRKPPFNGSIFa | 37.99 | 1394.746 | -0.3 | 698.38 | 2 | 40.92 | 20 | Amidation |
| PF | XP_006560385.1 | YRKPPFNGSIFa | 26.07 | 1323.709 | 0.1 | 662.8617 | 2 | 41.73 | 5 | Amidation |
| PF | XP_006560385.1 | RKPPFNGSIFa | 24.15 | 1160.645 | 0.4 | 581.3302 | 2 | 33.49 | 11 | Amidation |
| PF | XP_006562922.1 | GFKPEYISTAYGFa | 40.49 | 1477.724 | 0 | 739.8693 | 2 | 66.76 | 9 | Amidation |
| PF | XP_006565207.1 | SDPHLSILSKPMSAIPSYKFDD | 45.28 | 2447.204 | -0.6 | 816.7415 | 3 | 75.81 | 11 |  |
| PF | XP_006565207.1 | SPSLRLRFa | 30.48 | 973.5821 | -0.3 | 487.7982 | 2 | 28.72 | 19 | Amidation |
| PF | XP_006565207.1 | SDPHLSILS | 27.48 | 967.4974 | 0 | 484.756 | 2 | 36.83 | 8 |  |
| PF | XP_006565207.1 | SQRSPSLRLRFa | 24.3 | 1344.774 | 0.2 | 337.2008 | 4 | 20.45 | 5 | Amidation |
| PF | XP_006570344.1 | NSELINSLGLPKNMNNAa | 46.64 | 1940.015 | 0.2 | 971.015 | 2 | 91.06 | 12 | Amidation |
| PF | XP_006570344.1 | LINSLGLPKNMNNAa | 39.96 | 1609.897 | -0.2 | 805.9557 | 2 | 65.66 | 10 | Amidation |
| PF | XP_016769998.1 | LVDHRIPDLENEMFDSGNDPGSTVVRT | 45.09 | 3012.425 | -0.7 | 754.1129 | 4 | 66.6 | 23 |  |
| NF | Q868G6.1 | NSIINDVKNELFPEDIN | 71.58 | 1972.974 | -0.7 | 987.4937 | 2 | 96.19 | 11 |  |
| NF | Q868G6.1 | VLSMDGYQNILDKKDELLGEWE | 58.61 | 2594.257 | 5.3 | 1298.143 | 2 | 95.2 | 4 |  |
| NF | A8CL69.1 | TSQDITSGMWFGPRLa | 41.46 | 1693.825 | 1.1 | 847.9205 | 2 | 75.51 | 4 | Amidation |
| NF | A8CL69.1 | pQLHNIVDKPRQNFNDPRF | 38.15 | 2220.119 | -1.1 | 556.0364 | 4 | 40.26 | 9 | Pyro-glu from Q |
| NF | A8CL69.1 | pQITQFTPRLa | 18.4 | 1084.603 | 2 | 543.3098 | 2 | 65.6 | 8 | Pyro-glu from Q;<br>Amidation |
| NF | A8CL69.1 | GMWFGPRLa | 36.12 | 961.4956 | 1.3 | 481.7557 | 2 | 50.46 | 11 | Amidation |
| NF | A8CL69.1 | MWFGPRLa | 26.19 | 904.4741 | 0.9 | 453.2448 | 2 | 53.31 | 9 | Amidation |
| NF | A8CL69.1 | pQLHNIVDKPRQN | 50.72 | 1443.758 | 0.9 | 482.2604 | 3 | 14.04 | 8 | Pyro-glu from Q |
| NF | A8CL69.1 | pQLHNIVDKP | 45.19 | 1045.556 | 0.8 | 523.7855 | 2 | 23.94 | 9 | Pyro-glu from Q |
| NF | A8CL69.1 | RVPWTPSPRLa | 41.57 | 1206.699 | -0.9 | 604.356 | 2 | 23.89 | 5 | Amidation |
| NF | ACI90290.1 | TWKSPDIVIRFa | 51.85 | 1359.766 | -0.7 | 454.2624 | 3 | 56.58 | 5 | Amidation |
| NF | ACI90290.1 | GRNDLNFIRYa | 36.79 | 1265.663 | 0.2 | 633.8388 | 2 | 30.39 | 9 | Amidation |
| NF | ACI90290.1 | QITQFTPRLa | 33.64 | 1101.63 | -0.4 | 551.8218 | 2 | 32.4 | 10 | Amidation |
| NF | ACI90290.1 | AGFKNLNREQ | 36.45 | 1175.605 | -0.1 | 588.8096 | 2 | 10.17 | 12 |  |
| NF | ACI90290.1 | SPDIVIRFa | 33.7 | 944.5443 | -0.9 | 473.279 | 2 | 45.58 | 8 | Amidation |
| NF | NP_001161192.1 | PEIFTSPEELRRYIDHVS DYLLSGKARYa | 46.23 | 3515.784 | 0.6 | 586.9714 | 6 | 94.48 | 7 | Amidation |
| NF | P85527.1 | pQDVDHVFLRFa | 52.69 | 1256.63 | 0 | 629.3223 | 2 | 72.23 | 6 | Pyro-glu from Q;<br>Amidation |
| NF | P85527.1 | QDVDHVFLRFa | 48.09 | 1273.657 | 0.3 | 425.5596 | 3 | 48.21 | 5 | Amidation |
| NF | P85527.1 | pQDVDHVFLRF | 49.46 | 1257.614 | 0.2 | 629.8145 | 2 | 74.75 | 6 | Pyro-glu from Q |
| NF | P85527.1 | pQDVDHVFLR | 43.61 | 1110.546 | 2.4 | 556.2815 | 2 | 30.9 | 8 | Pyro-glu from Q |
| NF | P85527.1 | pQDVDHVFL | 24.41 | 954.4447 | 0.5 | 478.2299 | 2 | 70.18 | 6 | Pyro-glu from Q |
| NF | P85798.1 | LRNQLDIGDLQ | 49.85 | 1283.683 | 0 | 642.8489 | 2 | 41.44 | 6 |  |
| NF | P85798.1 | LRNQLDIGDL | 46.57 | 1155.625 | 0.7 | 578.8201 | 2 | 35.03 | 3 |  |
| NF | P85798.1 | IPAADKERLLN | 48.73 | 1238.698 | 0.9 | 620.3569 | 2 | 15.09 | 7 |  |
| NF | P85799.1 | SQAYDPYSNAAQFQLSSQSRGYPYQHRL<br>VY | 76.59 | 3523.655 | -0.9 | 881.9201 | 4 | 60.11 | 31 |  |
| NF | P85799.1 | SQAYDPYSNAAQFQLSSQSRGYPYQHRL | 66.47 | 3261.523 | -1.1 | 816.3871 | 4 | 51.7 | 6 |  |
| NF | P85799.1 | SQAYDPYSNAAQFQLSSQSRGYPYQHRL<br>V | 62.02 | 3360.591 | -2 | 841.1534 | 4 | 55.98 | 5 |  |
| NF | P85799.1 | LPTNLAEDTKKTEQTMRPKS | 60.38 | 2287.184 | -0.6 | 572.803 | 4 | 20.72 | 19 |  |
| NF | P85799.1 | NVPIYQEPRF | 46.81 | 1261.646 | -0.8 | 631.8295 | 2 | 45.2 | 8 |  |
| NF | P85799.1 | VPIYQEPRF | 45.86 | 1147.603 | -0.2 | 574.8084 | 2 | 42.06 | 7 |  |
| NF | P85799.1 | GYPYQHRLVY | 39.06 | 1294.646 | -0.3 | 648.33 | 2 | 23.28 | 7 |  |
| NF | P85799.1 | PIYQEPRF | 25.88 | 1048.534 | 0.2 | 525.2745 | 2 | 45.14 | 3 |  |
| NF | P85828.1 | ITGQGNRIF | 48.34 | 1004.54 | 0.4 | 503.2776 | 2 | 19.51 | 3 |  |
| NF | P85828.1 | SLKAPFA | 40.92 | 732.417 | 0.1 | 367.2158 | 2 | 29.4 | 5 |  |
| NF | P85828.1 | SLKAPF | 35.22 | 661.3799 | -0.1 | 331.6972 | 2 | 23.98 | 5 |  |
| NF | P85829.1 | MVPVPVHHMADELLRNGPDTVI | 49.4 | 2439.24 | 0.1 | 814.0874 | 3 | 74.93 | 6 |  |
| NF | P85829.1 | VHHMADELLRNGPDTVI | 42.7 | 1915.957 | -0.5 | 639.6594 | 3 | 44.62 | 5 |  |
| NF | P85829.1 | VPVPVHHMADELL | 49.99 | 1455.754 | 0.2 | 728.8846 | 2 | 33.13 | 7 |  |
| NF | P85829.1 | LLRNGPDTVI | 35.24 | 1096.624 | -0.1 | 549.3192 | 2 | 28.76 | 11 |  |
| NF | P85829.1 | LRNGPDTVI | 28.81 | 983.54 | 0.5 | 492.7775 | 2 | 16.64 | 12 |  |
| NF | P85830.1 | GLDLGLSRGFSGSQAAKHLMGAAANYA<br>GGPa | 75.12 | 2985.524 | 0.3 | 996.1823 | 3 | 83.37 | 8 | Amidation |
| NF | P85830.1 | GLDLGLSRGFSGSQAAKH | 47.59 | 1799.928 | -0.2 | 600.9831 | 3 | 30.26 | 7 |  |
| NF | P85830.1 | GLDLGLSRGFSGSQAAKHLMa | 34.44 | 2043.068 | 1.2 | 682.0309 | 3 | 54.93 | 8 | Amidation |
| NF | P85830.1 | GLDLGLSRGFSGSQAA | 70.04 | 1534.774 | 0.5 | 768.3947 | 2 | 43.37 | 5 |  |
| NF | P85830.1 | HLMGLAAANYAGGPa | 47.07 | 1340.666 | -0.8 | 671.3397 | 2 | 35.45 | 8 | Amidation |
| NF | P85831.1 | IDLSRFYGHFNT | 66.33 | 1468.71 | -0.8 | 735.3616 | 2 | 62.1 | 15 |  |
| NF | P85831.1 | IDLSRFYGHFN | 59.58 | 1367.662 | 0 | 684.8384 | 2 | 59.48 | 12 |  |
| NF | P85831.1 | IDLSRFYGHF | 54.45 | 1253.619 | 0.1 | 627.817 | 2 | 66.68 | 7 |  |
| NF | P85831.1 | IDLSRFYGHFNTKR | 45.5 | 1752.906 | 0.5 | 585.3095 | 3 | 39.93 | 6 |  |
| NF | P85831.1 | DLSRFYGHFN | 33.5 | 1254.578 | -0.6 | 628.8198 | 2 | 65.09 | 14 |  |
| NF | P85831.1 | FYGHFNT | 44.01 | 884.3817 | -0.2 | 443.1981 | 2 | 20.92 | 13 |  |
| NF | P85832.1 | LTNYLATTGHGTNTGGPVLT | 85.24 | 1987.001 | -0.6 | 994.5072 | 2 | 47.09 | 8 |  |
| NF | P85832.1 | NLDEIDRVGWSGFV | 56.68 | 1605.779 | -0.9 | 803.8959 | 2 | 86.63 | 4 |  |
| NF | P85832.1 | NIDEIDRTAFDNFF | 54.74 | 1715.779 | -0.7 | 858.8962 | 2 | 94.63 | 7 |  |
| NF | P85832.1 | LTNYLATTGHGTNTGGPVLTRRFa | 52.84 | 2445.288 | 0 | 612.3292 | 4 | 38.23 | 12 | Amidation |
| NF | P85832.1 | ELVDELSPVSERETLERFa | 29.85 | 2146.091 | -1.2 | 716.3699 | 3 | 72.71 | 4 | Amidation |
| NF | P85832.1 | LVDELSPVSERETLERFa | 26.21 | 2017.048 | -1.1 | 673.3558 | 3 | 61.74 | 3 | Amidation |
| NF | P85832.1 | LTNYLATTGHGTNTGGPVL | 83.65 | 1885.953 | 3.4 | 943.9872 | 2 | 37.57 | 3 |  |
| NF | Q06601.1 | PNDMLSQRYHFGLa | 69.72 | 1575.762 | -0.4 | 526.2609 | 3 | 47.76 | 6 | Amidation |
| NF | Q06601.1 | AVHYSGGQPLGSKRPNDMLSQRYHFGLa | 66.38 | 3013.509 | 0.1 | 754.3846 | 3 | 41.05 | 10 | Amidation |
| NF | Q06601.1 | AYTYVSEYKRLPVYNFGla | 69.78 | 2181.126 | -1.2 | 1091.569 | 2 | 73.7 | 8 | Amidation |
| NF | Q06601.1 | ADYPLRLNLD | 50.74 | 1188.614 | 0.6 | 595.3146 | 2 | 57.13 | 15 |  |
| NF | Q06601.1 | YPLRLNLD | 44.94 | 1002.55 | 0.3 | 502.2823 | 2 | 49.7 | 21 |  |
| NF | Q06601.1 | LDYLPVDNPAFH | 54.33 | 1399.677 | 1.6 | 700.8469 | 2 | 61.58 | 5 |  |
| NF | Q06601.1 | RQYSFGLa | 34.76 | 868.4555 | 0.1 | 435.2351 | 2 | 23.68 | 7 | Amidation |
| NF | Q06601.1 | GRQPYSFGLa | 38.89 | 1022.53 | 0.2 | 512.2722 | 2 | 29.28 | 7 | Amidation |
| NF | Q06601.1 | GRDYSFGLa | 30.15 | 912.4453 | 0 | 457.2299 | 2 | 29.2 | 10 | Amidation |
| NF | Q06601.1 | WIDTNDNKRGRDYSFGLa | 36.47 | 2054.992 | 1 | 686.0054 | 3 | 34.51 | 7 | Amidation |
| NF | Q06601.1 | AVHYSGGQPLGS | 39.03 | 1171.562 | 0.3 | 586.7885 | 2 | 13.57 | 11 |  |
| NF | Q06602.1 | EAEPEAEPGNNRPVYIPQPRPPHRL | 66.9 | 2959.505 | 3.2 | 592.9102 | 5 | 21.55 | 23 |  |
| NF | Q5DW47.1 | STSLEELANR | 36.25 | 1118.557 | 0.9 | 560.2861 | 2 | 24.18 | 7 |  |
| NF | Q5DW47.1 | STSLEELANRN | 39.72 | 1232.6 | 0.7 | 617.3075 | 2 | 23.07 | 9 |  |
| NF | Q5DW47.1 | pQTFTYSHGWTNa | 51.7 | 1322.568 | -0.6 | 662.2909 | 2 | 50.66 | 19 | Pyro-glu from Q;<br>Amidation |
| NF | Q868G6.1 | ASFDDEYYKRAPMGFQGMRa | 58.13 | 2267.025 | -0.1 | 567.7635 | 4 | 43.53 | 9 | Amidation |
| NF | Q868G6.1 | ARMGFHGMRa | 54.74 | 1060.517 | -0.5 | 531.2656 | 2 | 8.87 | 7 | Amidation |
| NF | Q868G6.1 | APMGFQGMRa | 49.97 | 992.4684 | 0 | 497.2415 | 2 | 19.16 | 6 | Amidation |
| NF | Q868G6.1 | SLEEILDEIK | 47.34 | 1187.629 | 0.3 | 594.8217 | 2 | 84.58 | 9 |  |
| NF | Q868G6.1 | NPRWEFRGKFVGV | 45.98 | 1590.842 | 0.3 | 531.2881 | 3 | 40.1 | 9 |  |
| NF | Q868G6.1 | SPFRYLGARa | 42.47 | 1064.588 | 0.1 | 533.3013 | 2 | 15.98 | 4 | Amidation |
| NF | Q868G6.1 | ALMGFQGVRa | 42.22 | 976.5276 | -0.3 | 489.2709 | 2 | 29.18 | 6 | Amidation |
| NF | Q868G6.1 | NPRWEFRGKFVGVRa | 40.99 | 1745.959 | -0.2 | 437.4969 | 4 | 21.41 | 7 | Amidation |
| NF | Q868G6.1 | SPFRYLGA | 40.83 | 909.4708 | 0 | 455.7427 | 2 | 35.07 | 7 |  |
| NF | Q868G6.1 | GVMDFQIGLQRKKD | 40.37 | 1633.861 | 0.1 | 545.6276 | 3 | 34.17 | 7 |  |
| NF | Q868G6.1 | SPFRYLGARG | 39.19 | 1122.593 | -0.4 | 375.2049 | 3 | 19.38 | 4 |  |
| NF | Q868G6.1 | ALMGFQGVRG | 38.17 | 1034.533 | -0.1 | 518.2737 | 2 | 34.43 | 13 |  |
| NF | Q868G6.1 | GVMDFQIGLQ | 37.68 | 1106.543 | 0.3 | 554.2789 | 2 | 84.25 | 6 |  |
| NF | Q868G6.1 | VLSMDGYQNILD | 36.56 | 1366.644 | 0.1 | 684.3292 | 2 | 79.28 | 3 |  |
| NF | Q868G6.1 | APMGFYGTRa | 36.12 | 997.4803 | 1.1 | 499.748 | 2 | 16.97 | 3 | Amidation |
| NF | Q868G6.1 | ASFDDEYY | 30.67 | 1008.371 | 0.1 | 505.1929 | 2 | 40.98 | 4 |  |
| NF | Q868G6.1 | SLEEILDEI | 23.62 | 1059.534 | -1.5 | 530.7733 | 2 | 104.74 | 3 |  |
| NF | Q868G6.1 | IILDALEELD | 28 | 1142.607 | 0.6 | 572.3112 | 2 | 102.74 | 5 |  |
| NF | Q868G6.1 | APMGFQGMRG | 50.06 | 1050.474 | -0.3 | 526.2441 | 2 | 23.14 | 5 |  |
| NF | Q868G6.1 | APMGFYGTRG | 43.78 | 1055.486 | -0.3 | 528.7501 | 2 | 20.72 | 3 |  |
| NF | XP_006557714.1 | pQQFDDYGHLRFa | 59.46 | 1406.637 | 0.4 | 704.3259 | 2 | 41.18 | 4 | Pyro-glu from Q;<br>Amidation |
| NF | XP_006559359.1 | SVSSLAKNSAWPVSL | 70.04 | 1544.82 | 0 | 773.4172 | 2 | 66.98 | 6 |  |
| NF | XP_006559359.1 | NVASLARTYTLPQNAa | 66.96 | 1616.863 | -0.3 | 809.4387 | 2 | 42.45 | 6 | Amidation |
| NF | XP_006559359.1 | NVGSVAREHGLPYa | 62.66 | 1396.721 | -0.5 | 699.3675 | 2 | 20.47 | 11 | Amidation |
| NF | XP_006559359.1 | YVASLARTGDLPIRGQ | 57.46 | 1715.932 | 0.2 | 572.9846 | 3 | 35.02 | 9 |  |
| NF | XP_006559359.1 | FLLLPATDNNYFHQKLPSSLRSKSL | 51.46 | 2888.555 | 1.1 | 578.7189 | 5 | 69.14 | 13 |  |
| NF | XP_006559359.1 | SVSSLARTGDLPVREQ | 47.87 | 1713.901 | 0 | 572.3076 | 3 | 24.99 | 6 |  |
| NF | XP_006559359.1 | NIASLMRDYDQSRENRPFPa | 45.43 | 2406.186 | -0.5 | 803.069 | 3 | 60.51 | 5 | Amidation |
| NF | XP_006559359.1 | YVASLARTGDLPIRa | 35.3 | 1529.868 | 0.8 | 510.9636 | 3 | 31.89 | 4 | Amidation |
| NF | XP_006559359.1 | LPGSVILRALS | 34.9 | 1124.692 | 0.5 | 563.3534 | 2 | 70.13 | 8 |  |
| NF | XP_006559359.1 | NVGTLRADFALPPa | 33.49 | 1368.751 | 0.1 | 685.383 | 2 | 59.01 | 11 | Amidation |
| NF | XP_006559359.1 | GIFLPGSVILR | 32.7 | 1170.712 | 0.2 | 586.3636 | 2 | 76.91 | 5 |  |
| NF | XP_006559359.1 | GIFLPGSVILRALSQRa | 31.65 | 1725.041 | -1 | 576.0204 | 3 | 94.65 | 8 | Amidation |
| NF | XP_006559359.1 | HIGALARLGWLPSLR TARFS | 29.69 | 2221.26 | -0.6 | 556.3218 | 4 | 68.04 | 3 |  |
| NF | XP_006559359.1 | HIGALARLGWLPSLR TA | 24.36 | 1831.058 | 0.4 | 611.3602 | 3 | 67.47 | 5 |  |
| NF | XP_006559359.1 | NVGTLRADFALPPGRRNIASLMRDYDQSR<br>ENRPFPa | 19.42 | 4127.135 | 0.3 | 688.8633 | 6 | 72.55 | 6 | Amidation |
| NF | XP_006559865.1 | AFGLLTYPRIa | 34 | 1148.671 | -0.2 | 575.3424 | 2 | 68.55 | 5 | Amidation |
| NF | XP_006559865.1 | EKLKPNMRRAFGLLTYPRIa | 21.02 | 2301.325 | 0.5 | 576.3389 | 4 | 45.4 | 4 | Amidation |
| NF | XP_006560385.1 | AYRKPPFNGSIFa | 41.64 | 1394.746 | -0.1 | 698.3801 | 2 | 35.3 | 6 | Amidation |
| NF | XP_006560385.1 | KPPFNGSIFa | 36.34 | 1004.544 | 0 | 503.2794 | 2 | 42.18 | 7 | Amidation |
| NF | XP_006560385.1 | YRKPPFNGSIFa | 47.47 | 1323.709 | 0.3 | 662.8619 | 2 | 32.59 | 7 | Amidation |
| NF | XP_006560385.1 | RKPPFNGSIFa | 40.45 | 1160.645 | -0.2 | 581.3298 | 2 | 24.42 | 9 | Amidation |
| NF | XP_006562922.1 | GFKPEYISTAYGFa | 52.25 | 1477.724 | 0.6 | 739.8698 | 2 | 70.58 | 29 | Amidation |
| NF | XP_006565207.1 | SDPHLSILSKPMSAIPSYKFDD | 72.01 | 2447.204 | -0.3 | 816.7418 | 3 | 70.27 | 6 |  |
| NF | XP_006565207.1 | SQRSPSLRLRFa | 32.44 | 1344.774 | -0.2 | 449.2651 | 3 | 15.82 | 6 | Amidation |
| NF | XP_006565207.1 | SPSLRLRFa | 28.98 | 973.5821 | 0.5 | 487.7986 | 2 | 22.89 | 5 | Amidation |
| NF | XP_006565207.1 | SDPHLSILS | 38.23 | 967.4974 | 0 | 484.756 | 2 | 33.67 | 7 |  |
| NF | XP_006570344.1 | NSELINSLGLPKNMNNAa | 64.9 | 1940.015 | 3.8 | 971.0184 | 2 | 85.93 | 7 | Amidation |
| NF | XP_006570344.1 | LINSLGLPKNMNNAa | 54.36 | 1609.897 | -0.3 | 805.9557 | 2 | 61.69 | 6 | Amidation |
| NF | XP_016769998.1 | LVDHRIPDLENEMFDSGNDPGSTVVRT | 65.72 | 3012.425 | -0.4 | 1005.148 | 3 | 63.14 | 13 |  |
| NF | XP_016769998.1 | IGSLSIVNSMDVLRQRVLLELARRKALQD<br>QAQIDANRRLLLETIa | 34.17 | 4913.782 | -0.5 | 819.9706 | 6 | 96.85 | 12 | Amidation |
| NF | XP_016769998.1 | HPISYNTYDERELSRDHPPLLL | 33.16 | 2664.33 | 0.3 | 667.0898 | 4 | 52.44 | 6 |  |

**Table S4.**
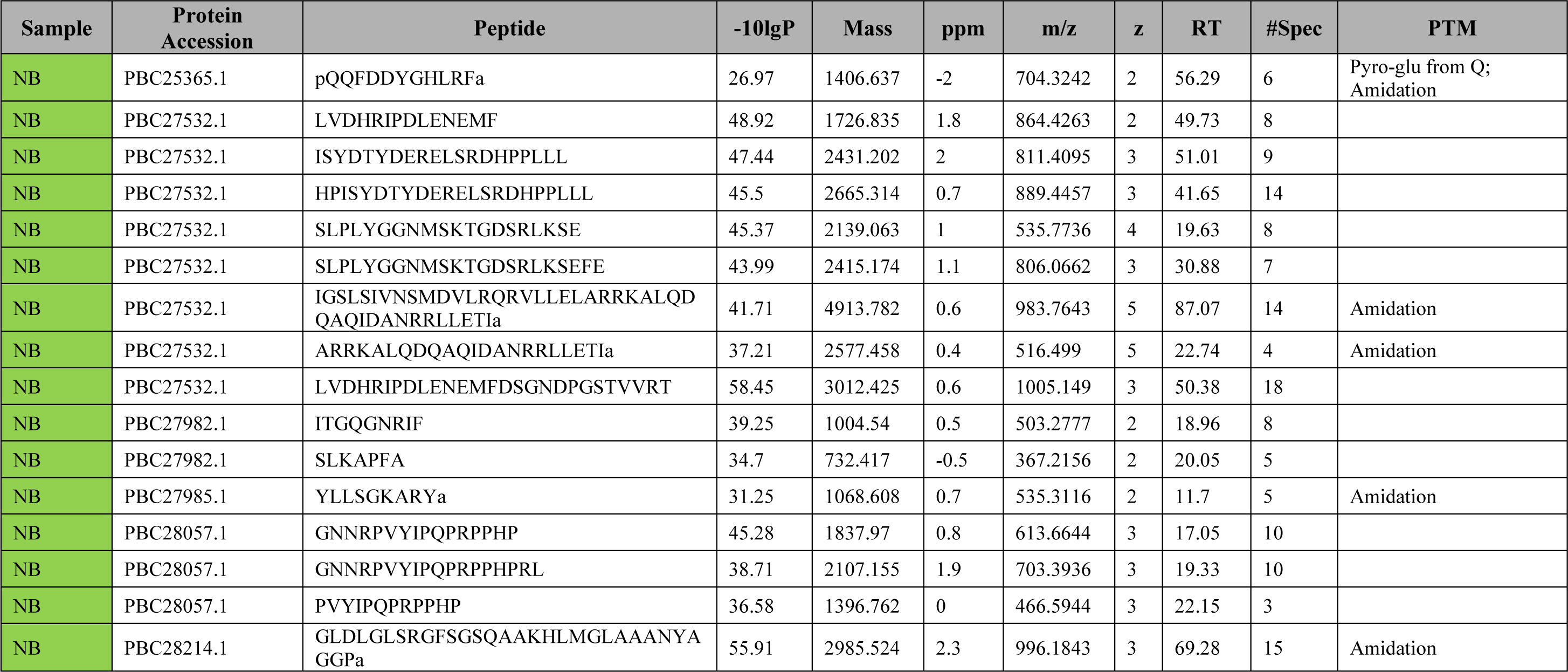
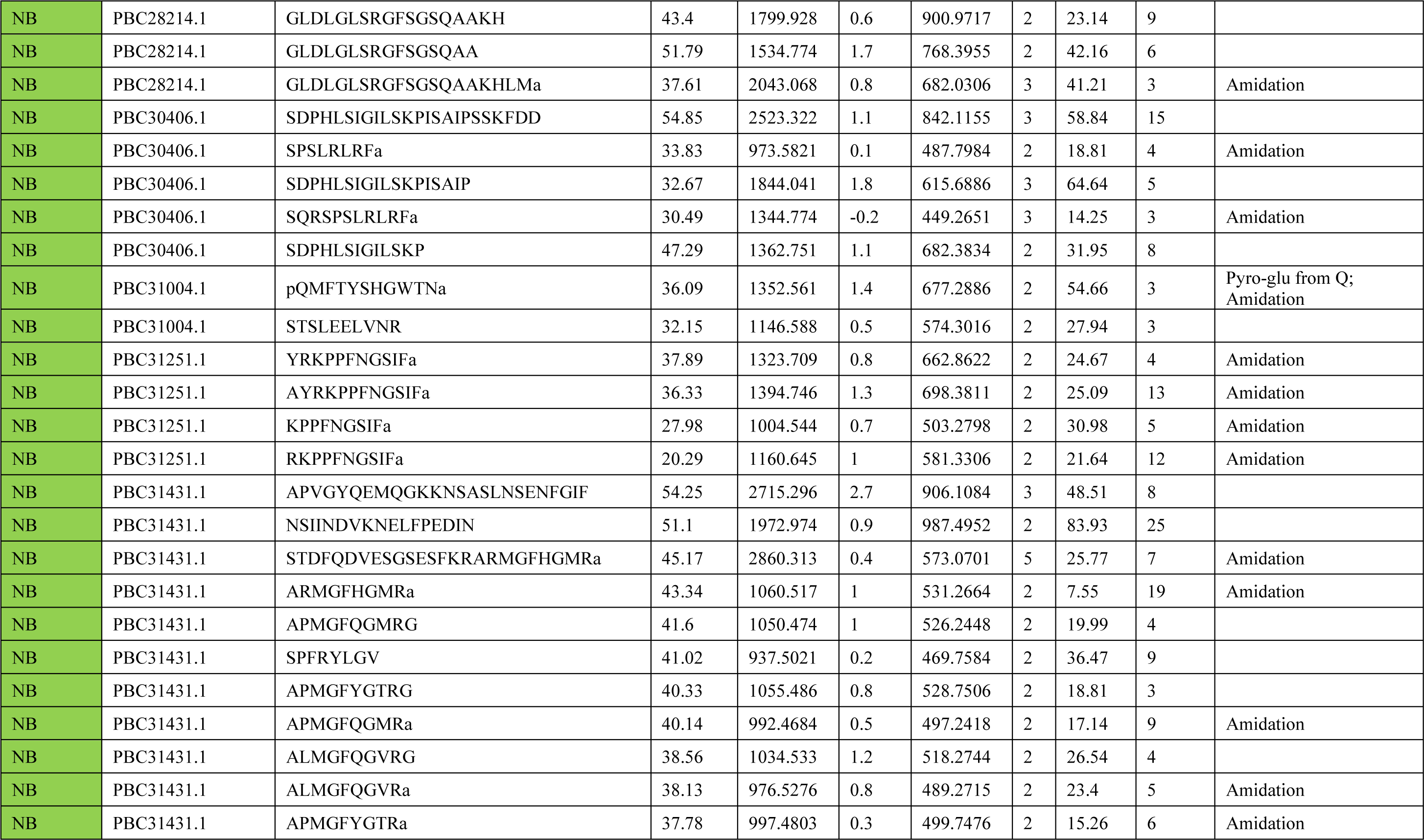
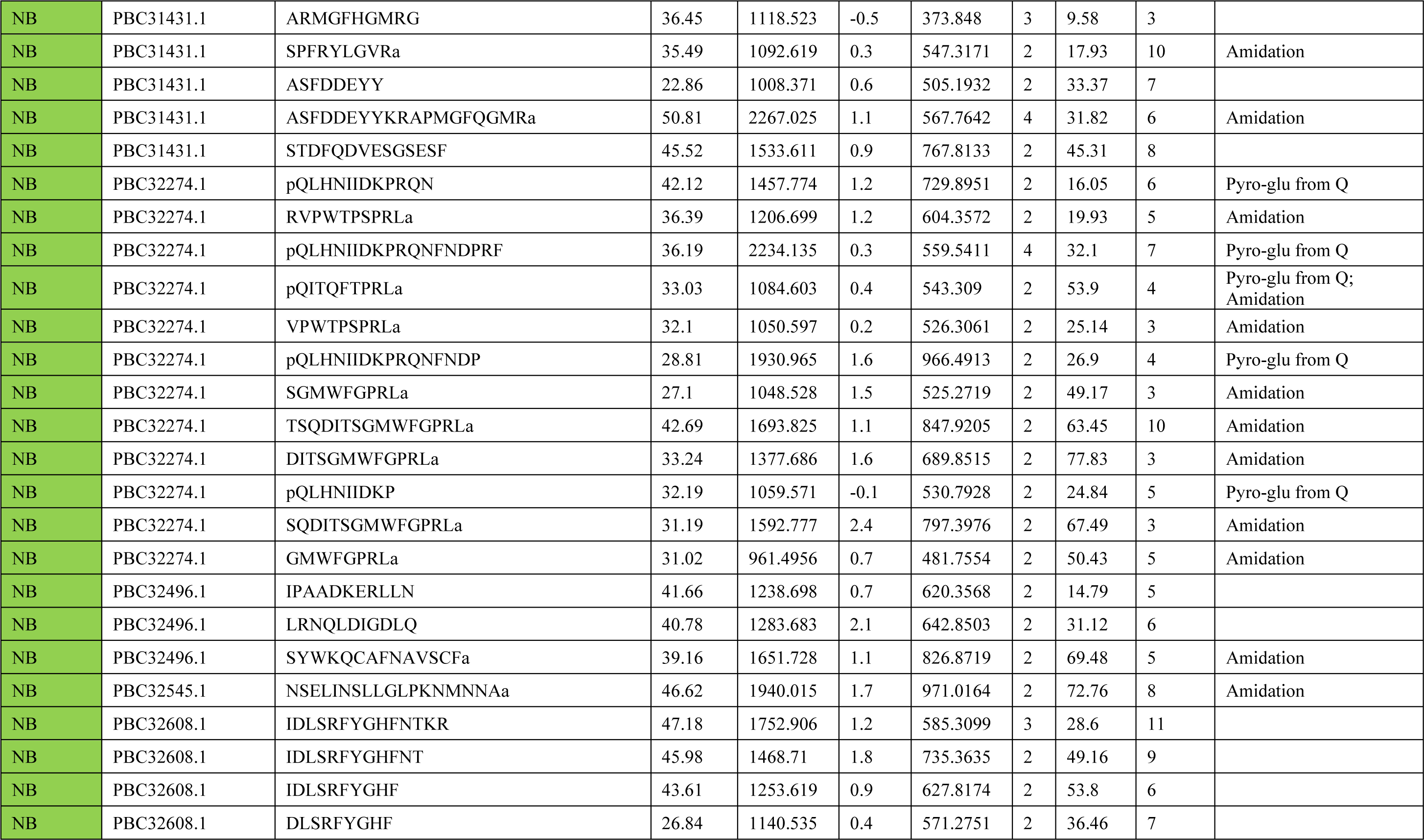
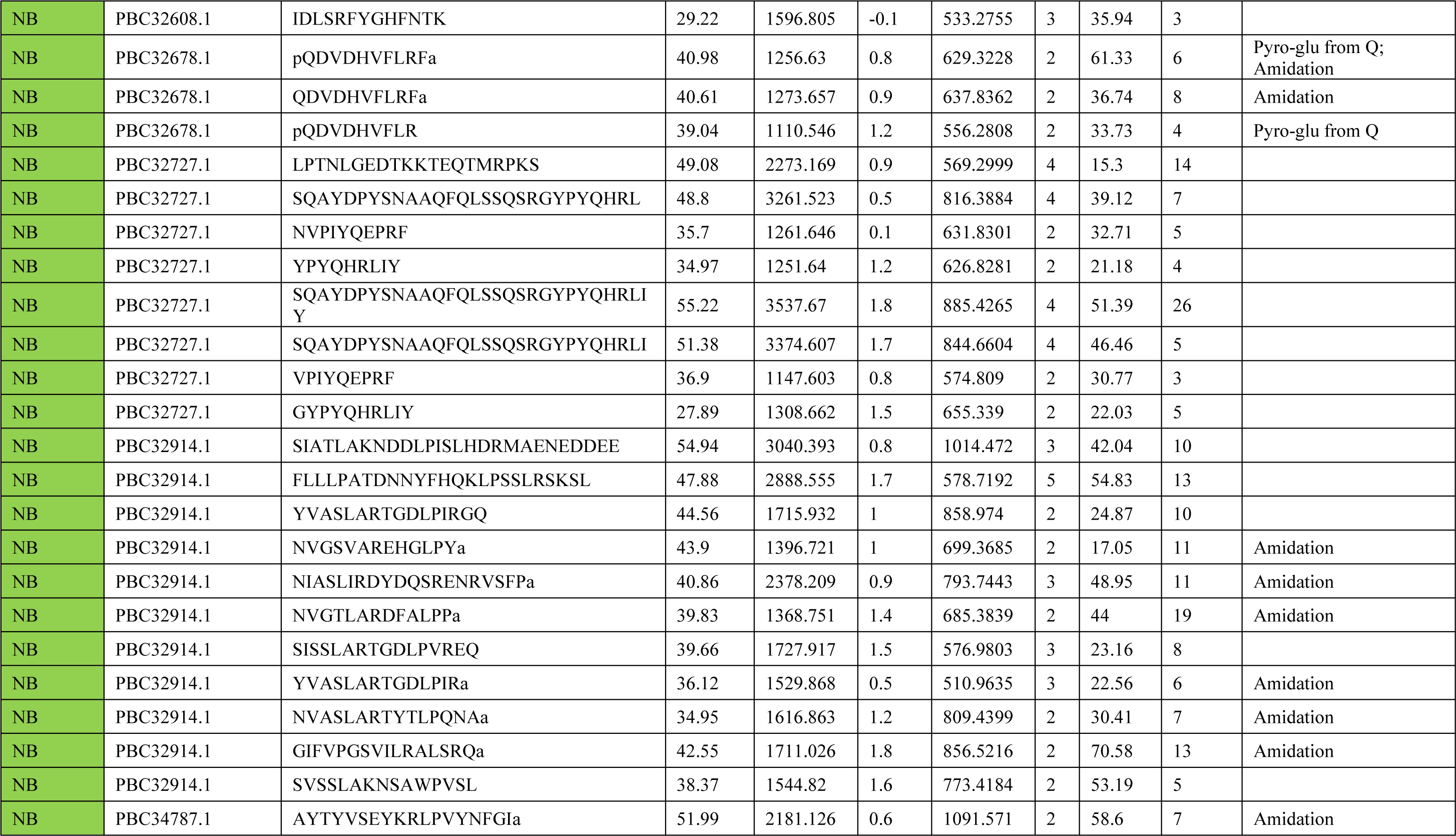
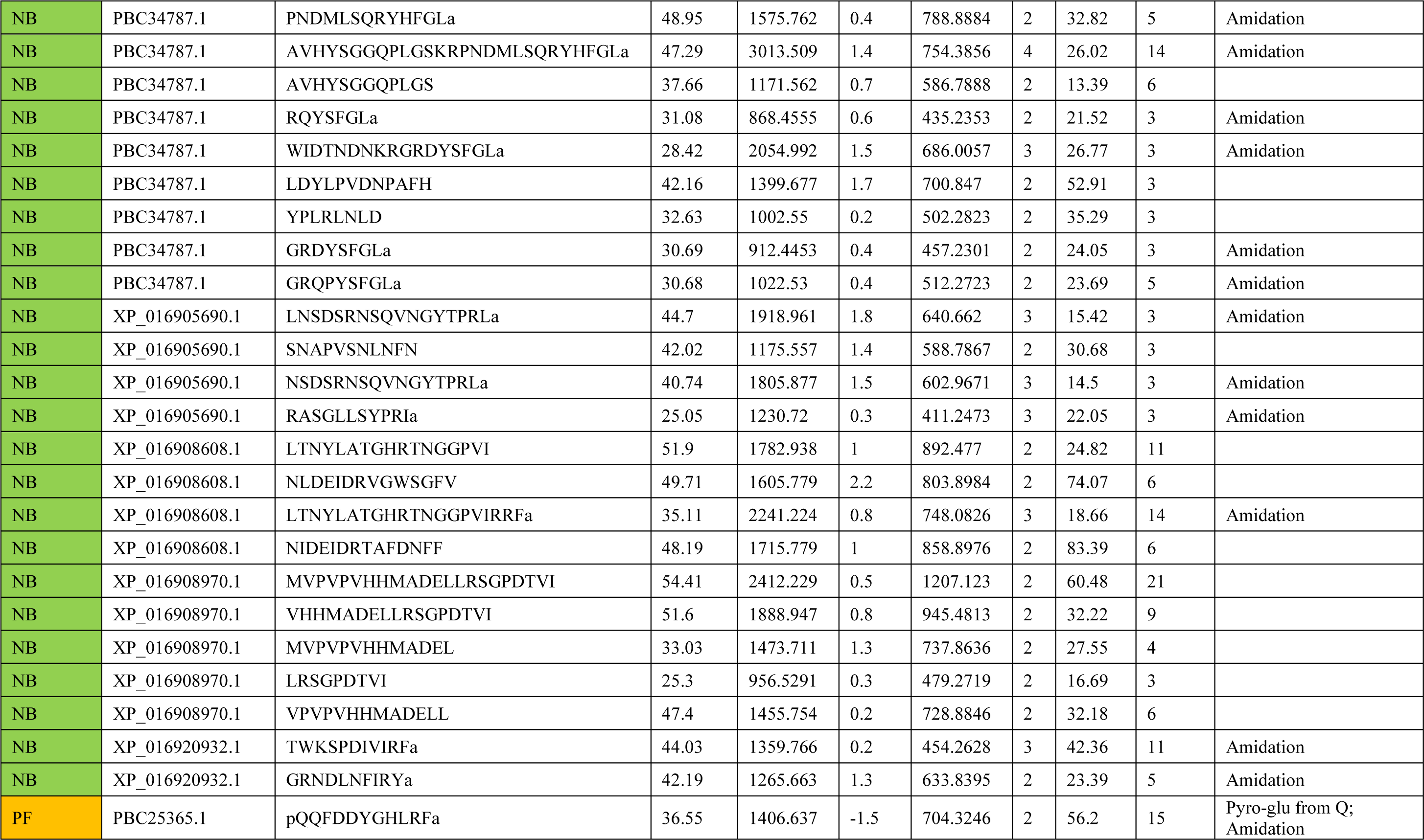
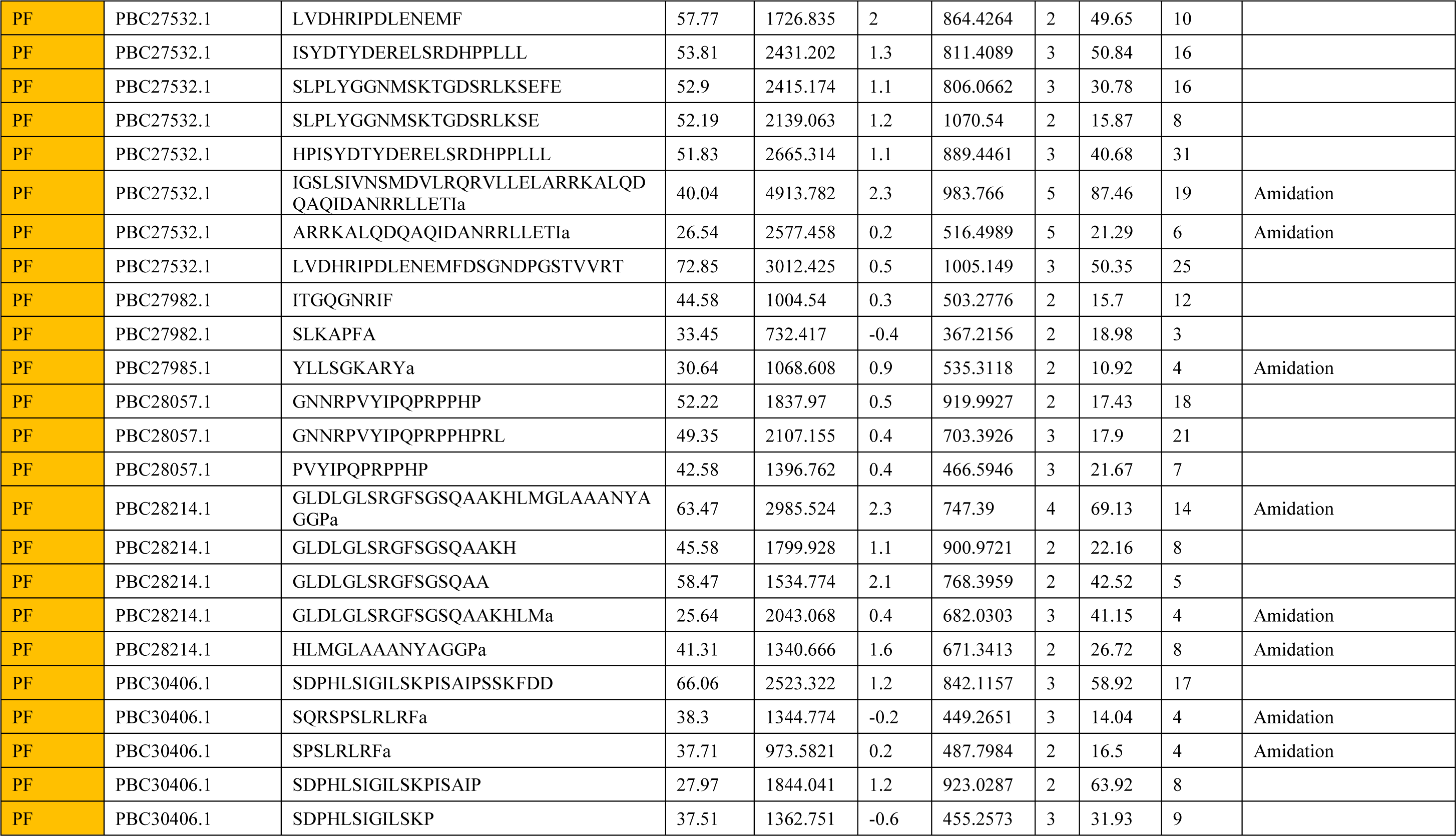
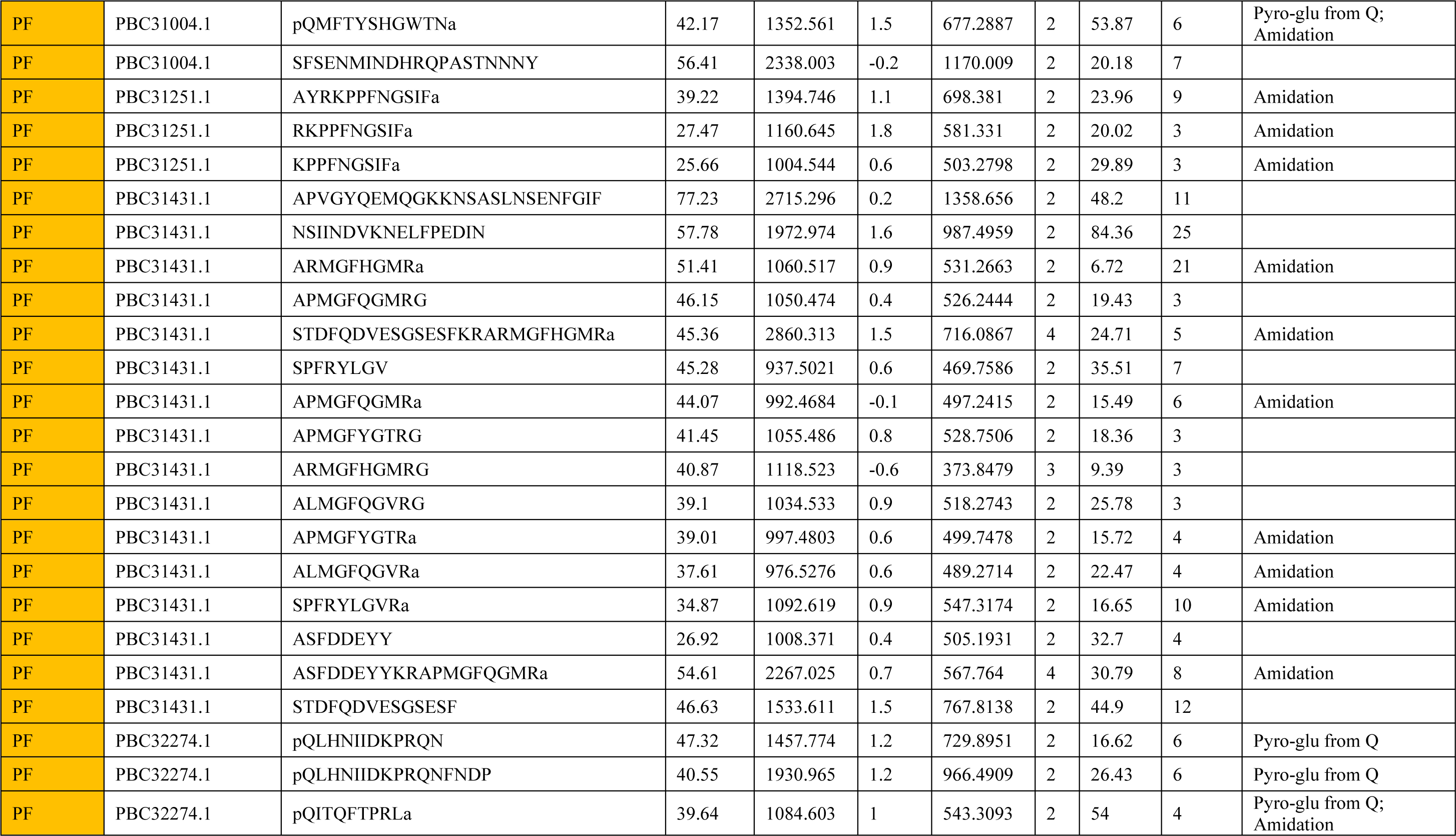
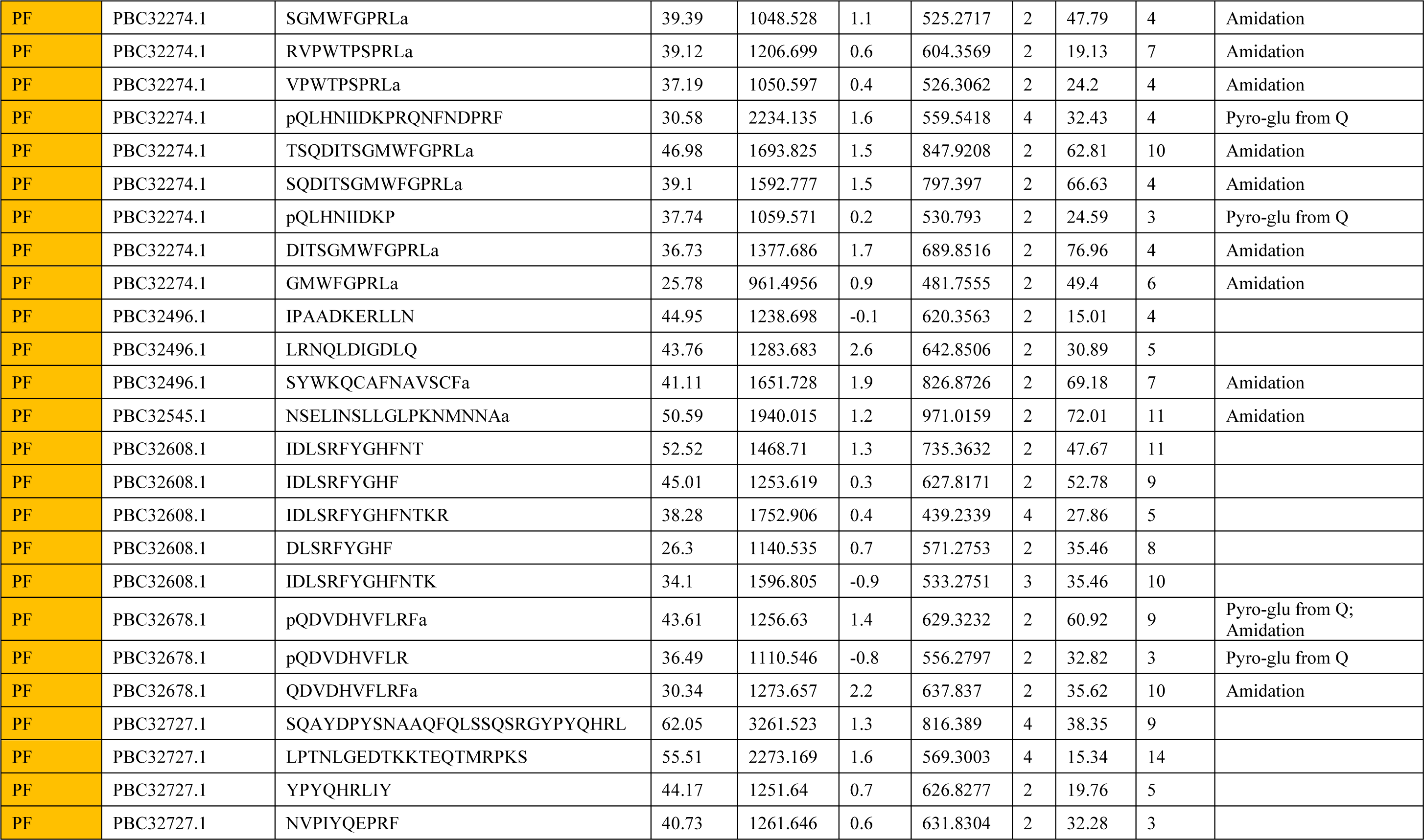
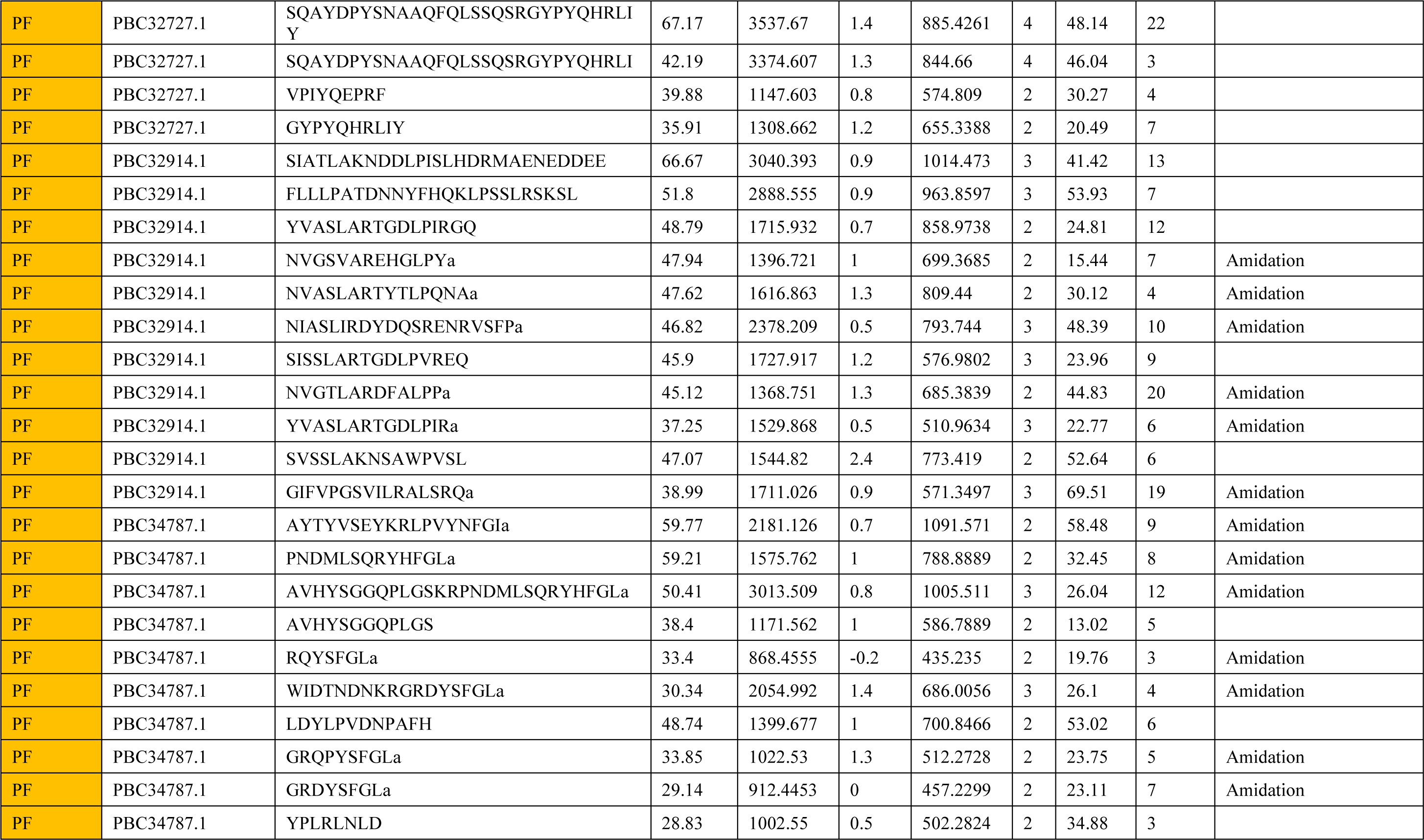
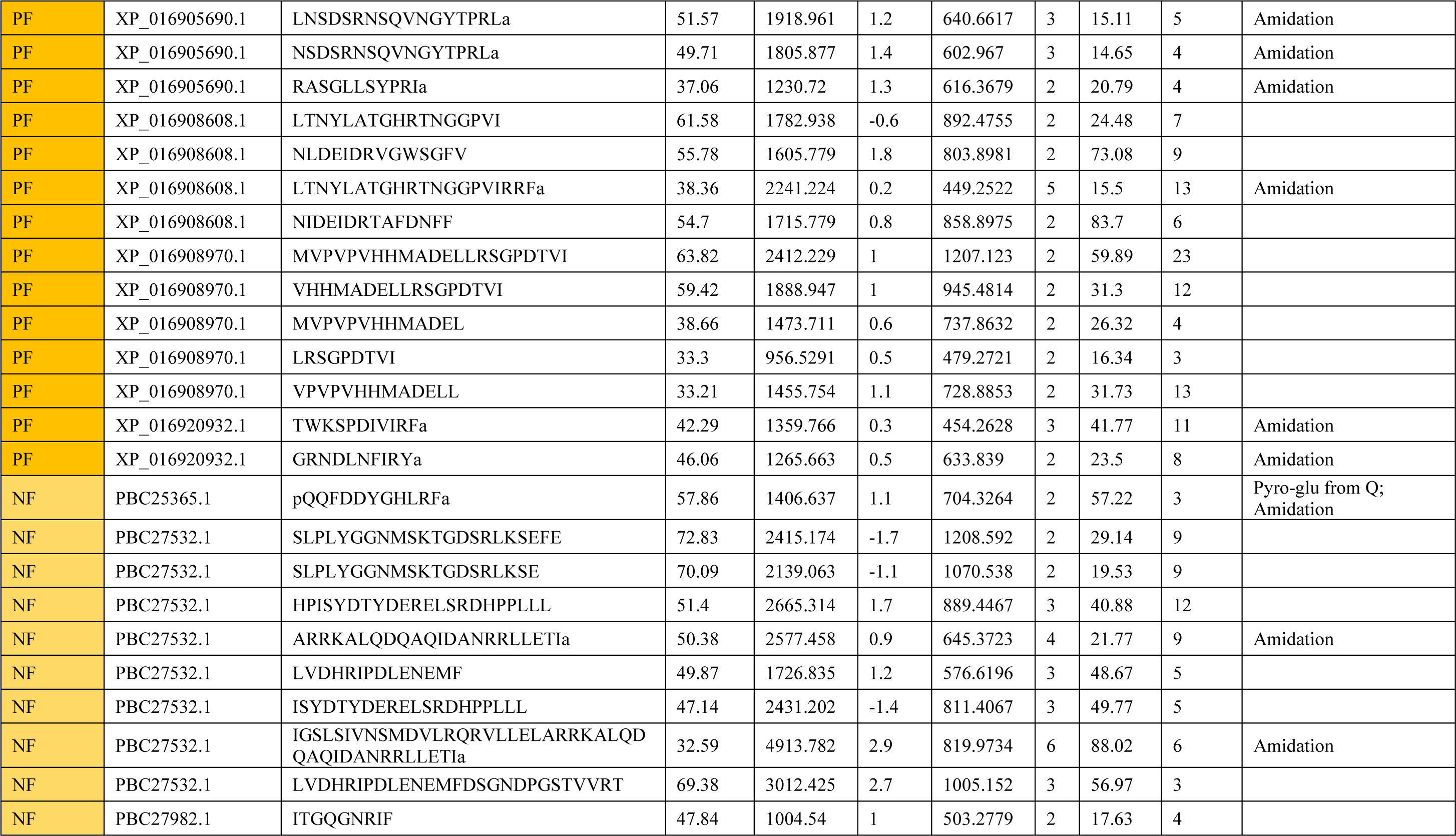
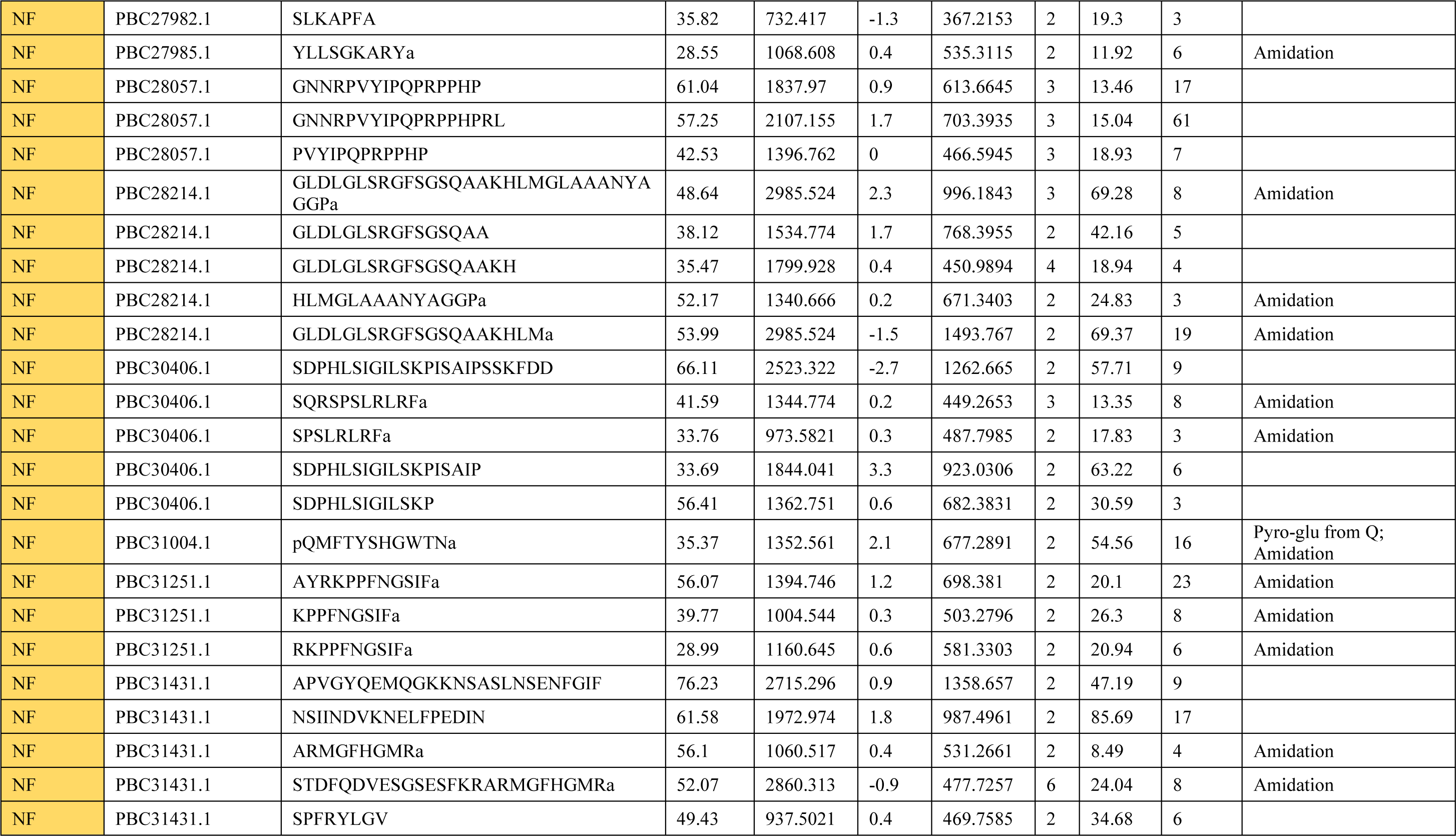
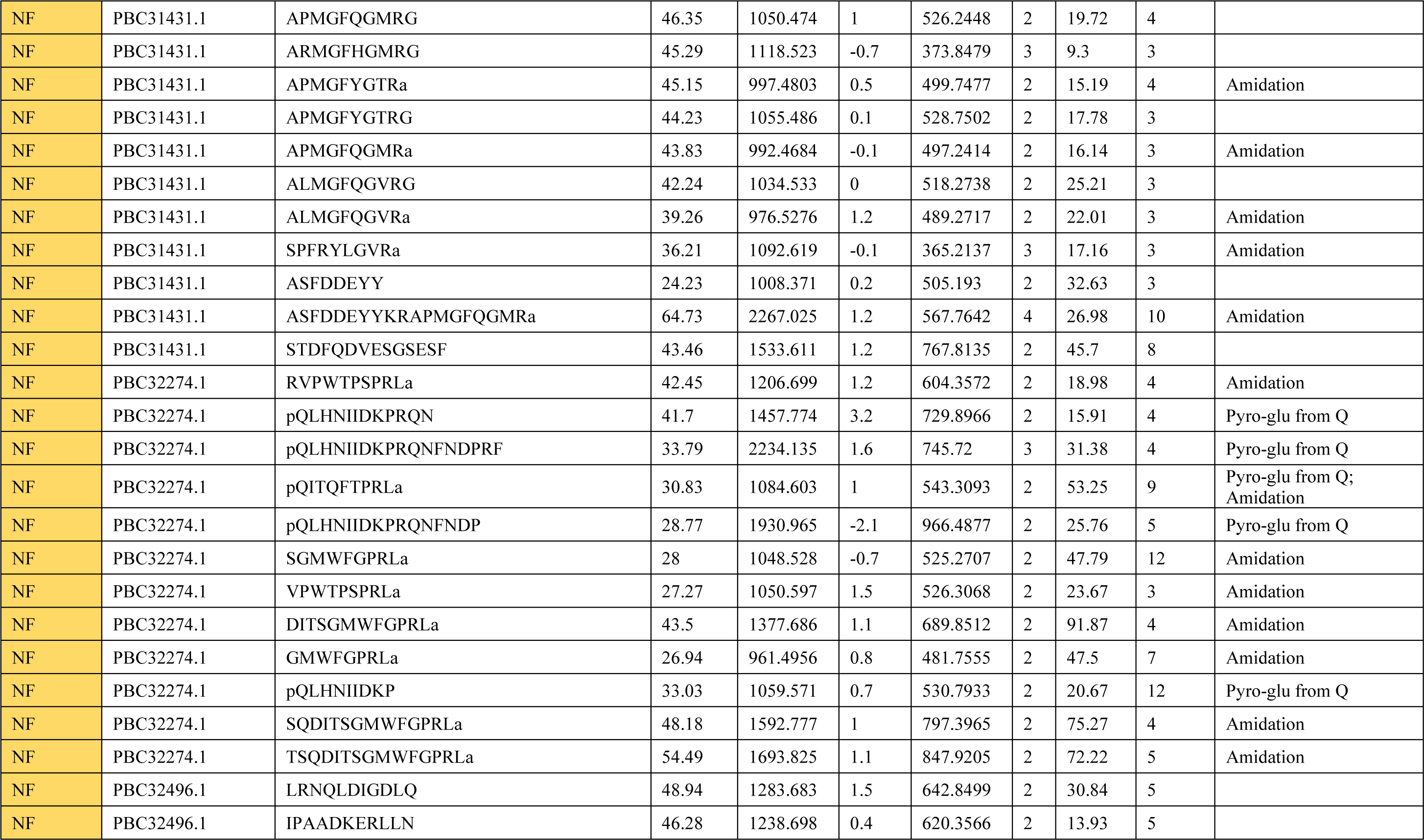
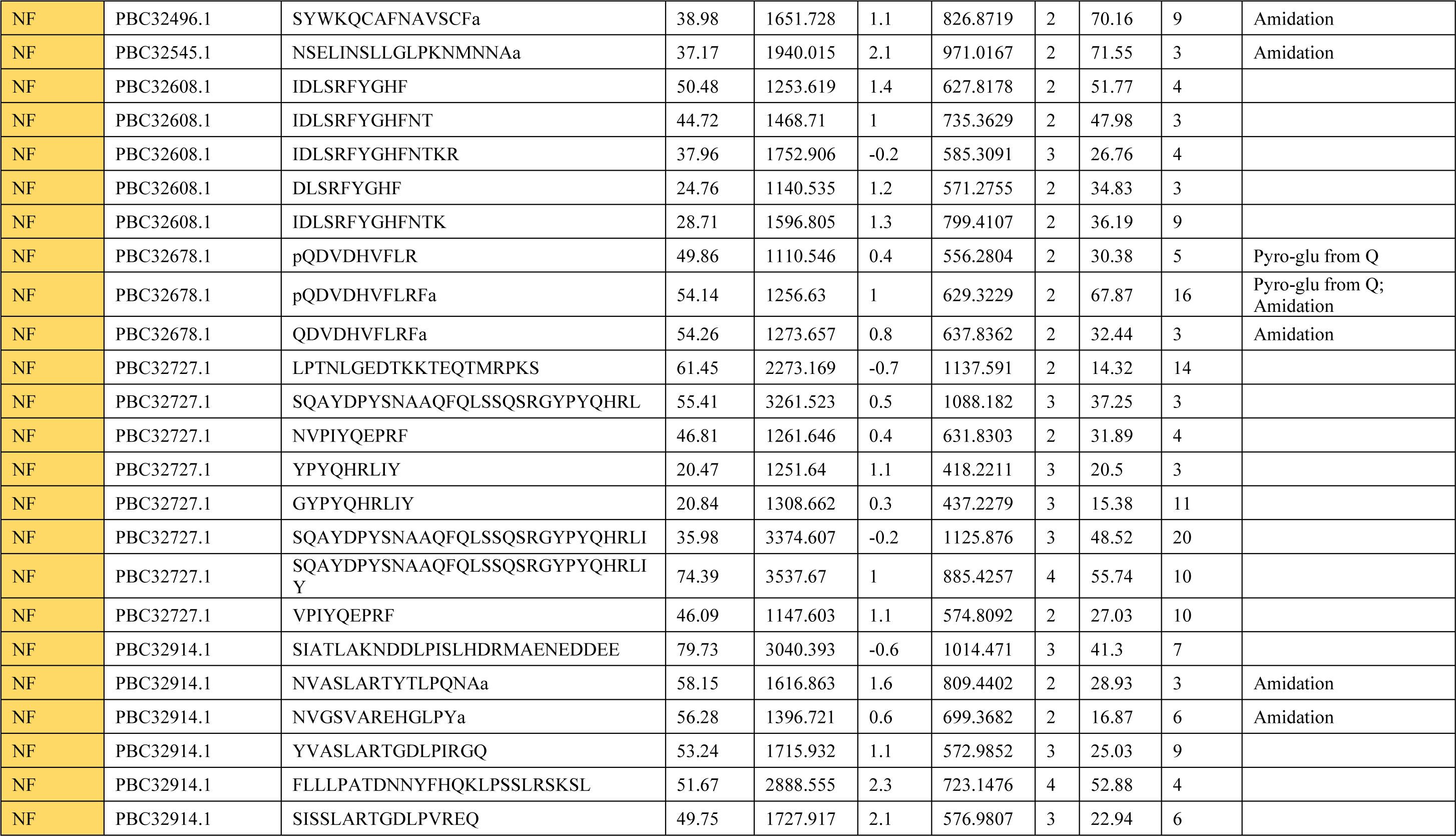
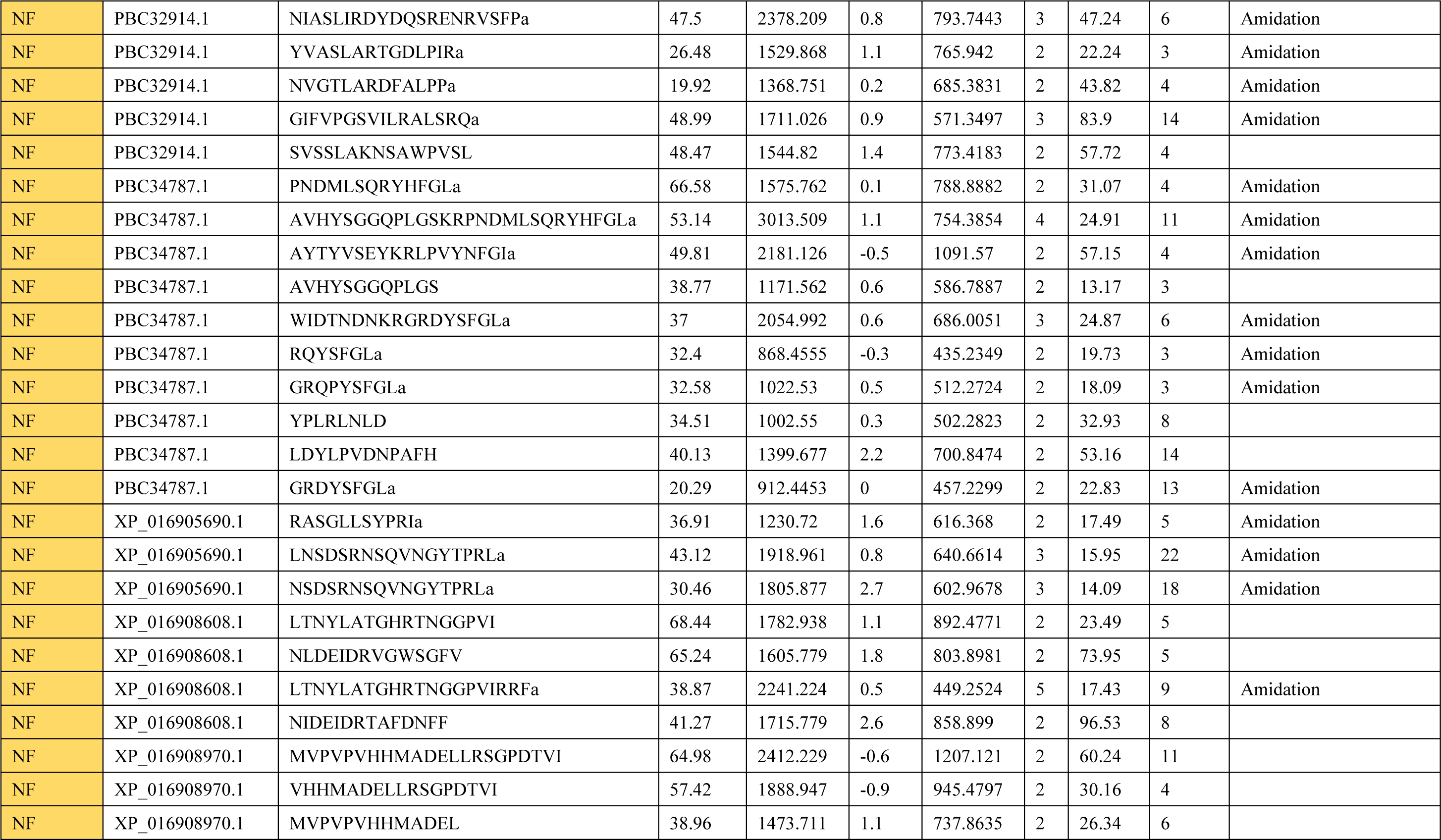
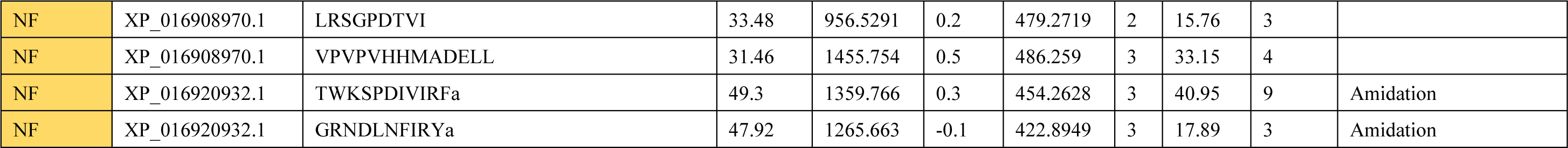
Neuropeptides identified in the brain of Apis cerana cerana workers. “NB” is nurse bee. “PF” is pollen forager. “NF” is nectar forager. “Protein Accession” is the unique number given to mark the entry of a protein in the database NCBInr. “Peptide” is the amino acid sequence of the peptide as determined in PEAKS Search. “-10lgP” is the score indicates the scoring significance of a peptide-spectrum match. “Mass” is monoisotopic mass of the peptide. “ppm” is precursor mass error, calculated as 10^6^ × (precursor mass - peptide mass) / peptide mass. “m/z” is precursor mass-to-charge ratio. “z” is peptide charge. “RT” is retention time (elution time) of the spectrums as recorded in the data. “#Spec” is the number of scanned spectrums of the peptide. “PTM” is post translational modification types present in the peptide.

**Table S5.**
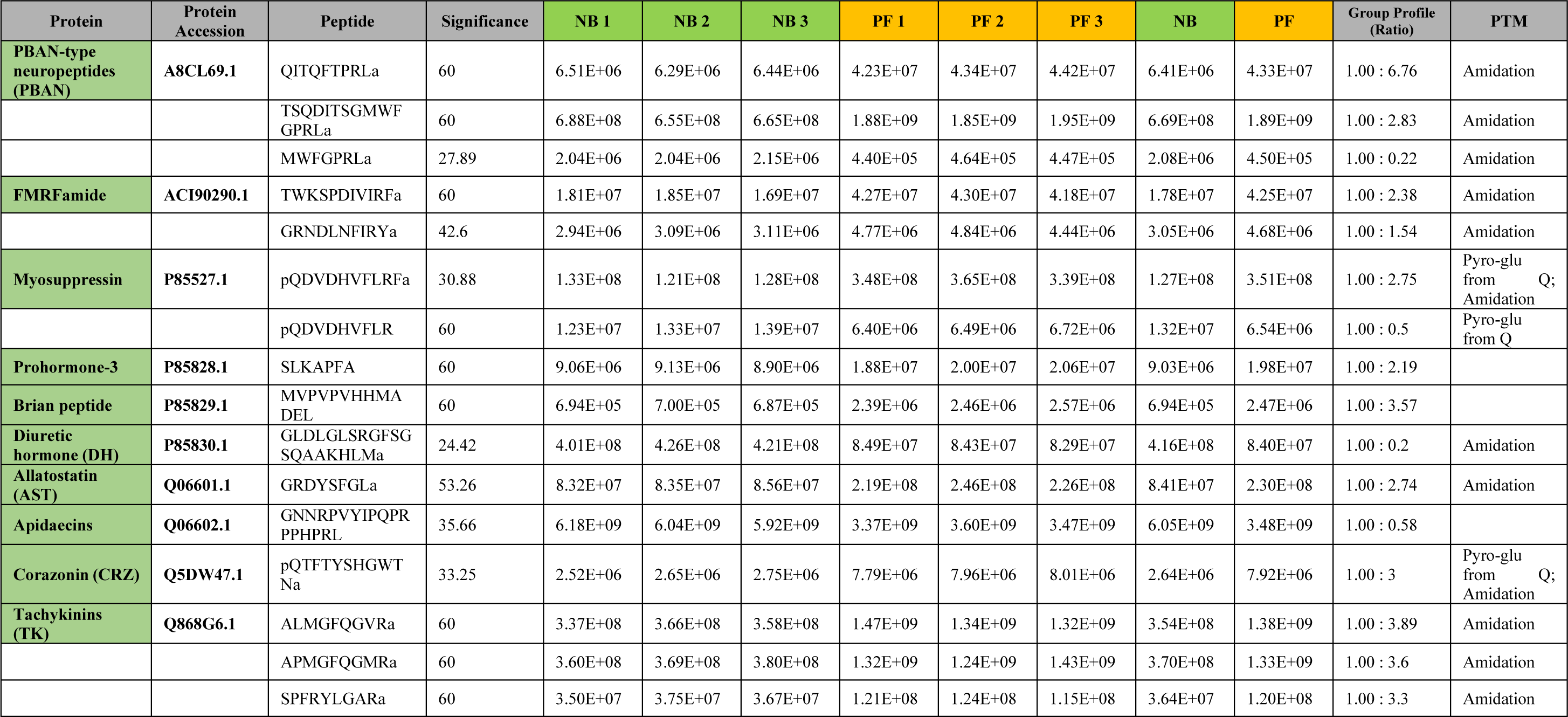
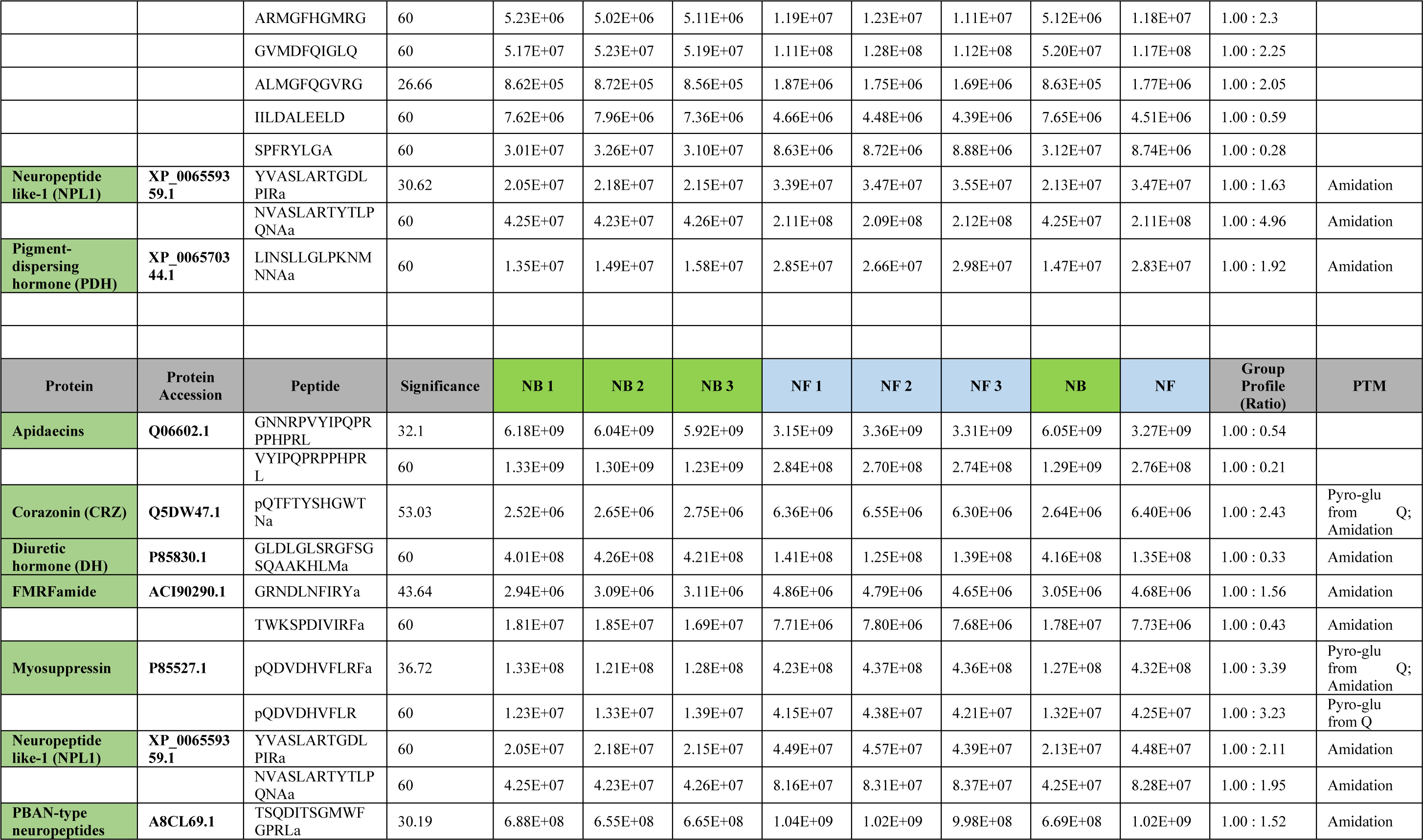
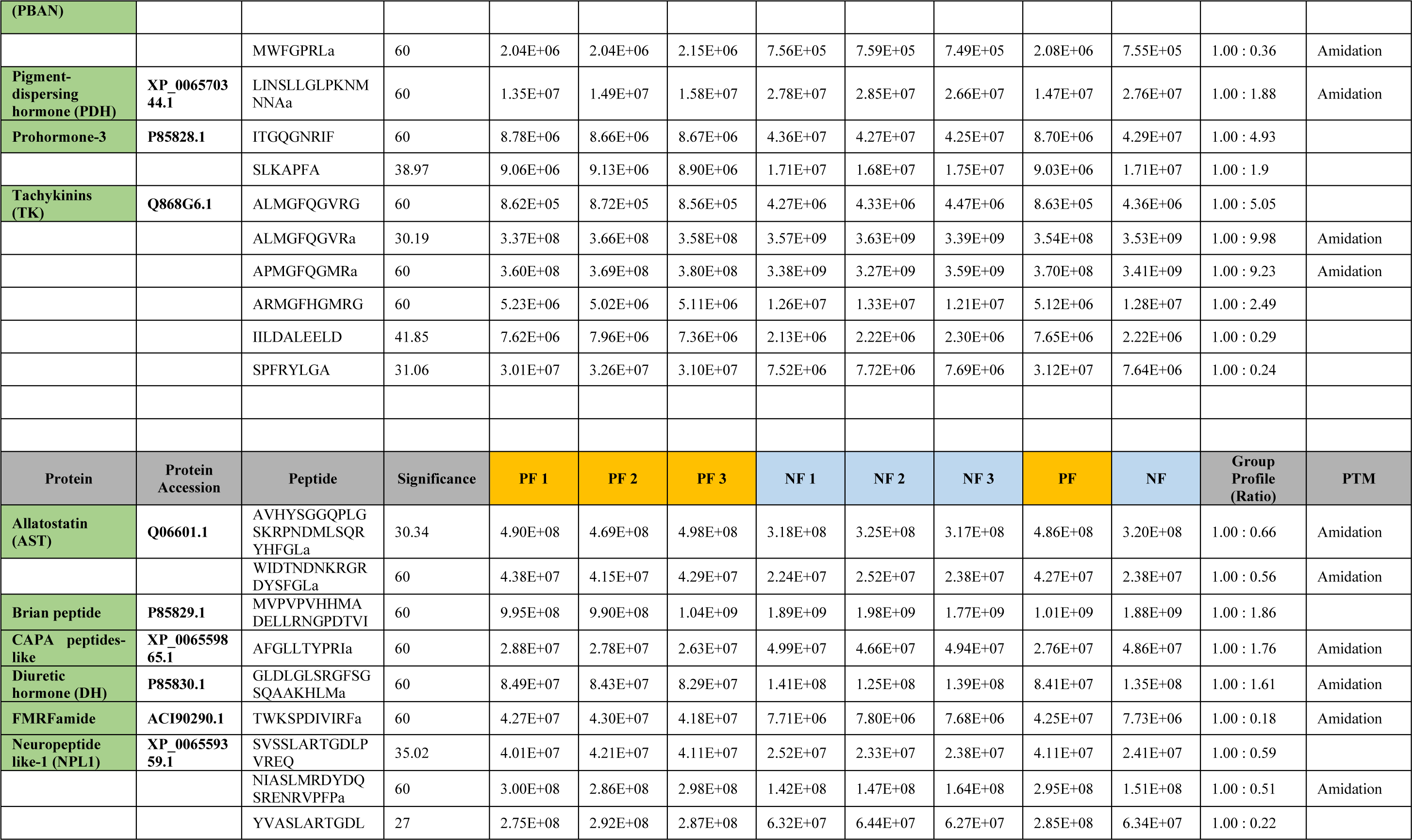
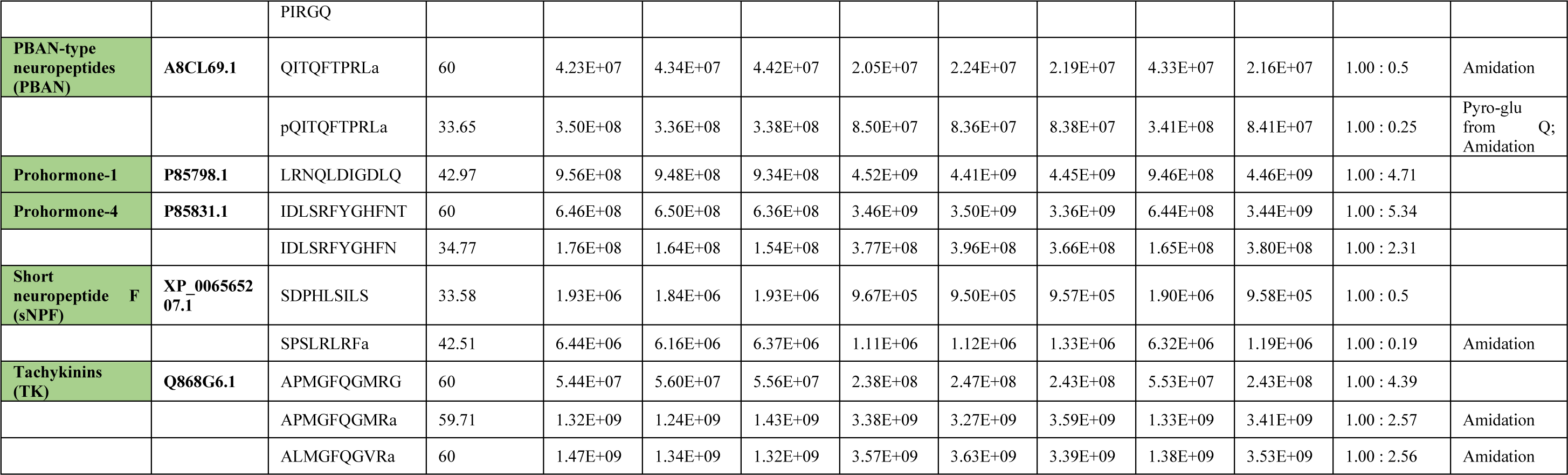
Quantitative neuropeptide comparison of different behavioral phenotypes of *Apis mellifera ligustica* workers. “Protein Accession” is the unique number given to mark the entry of a protein in the database NCBInr. “Peptide” is the amino acid sequence of the peptide. “Significance (-10lgP)” is the peptide confidence score. “NB” is nurse bee. “PF” is pollen forager. “NF” is nectar forager. “Group Profile (Ratio)” is the relative abundance ratio to the base group. “PTM” is post translational modification types present in the peptide.

**Table S6.**
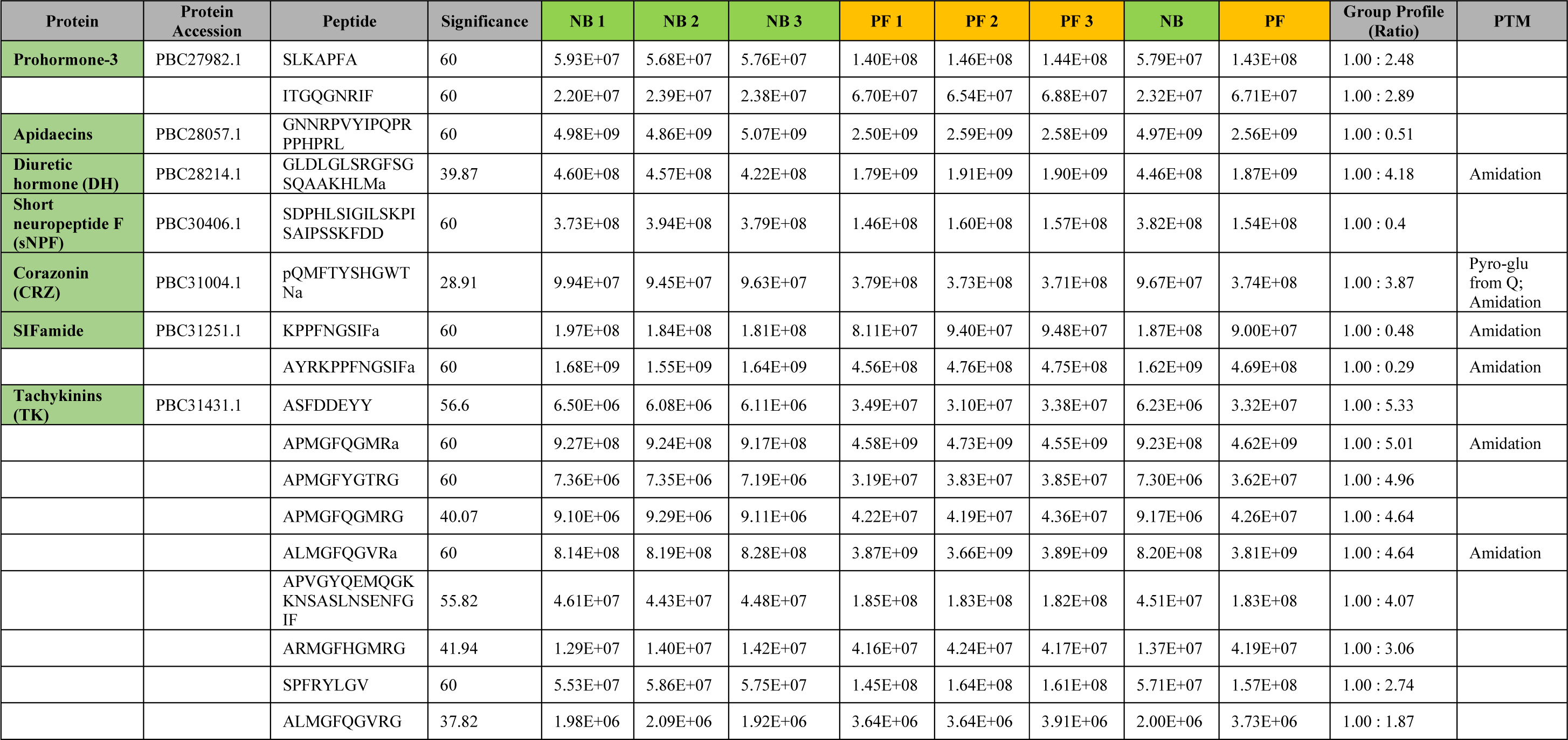
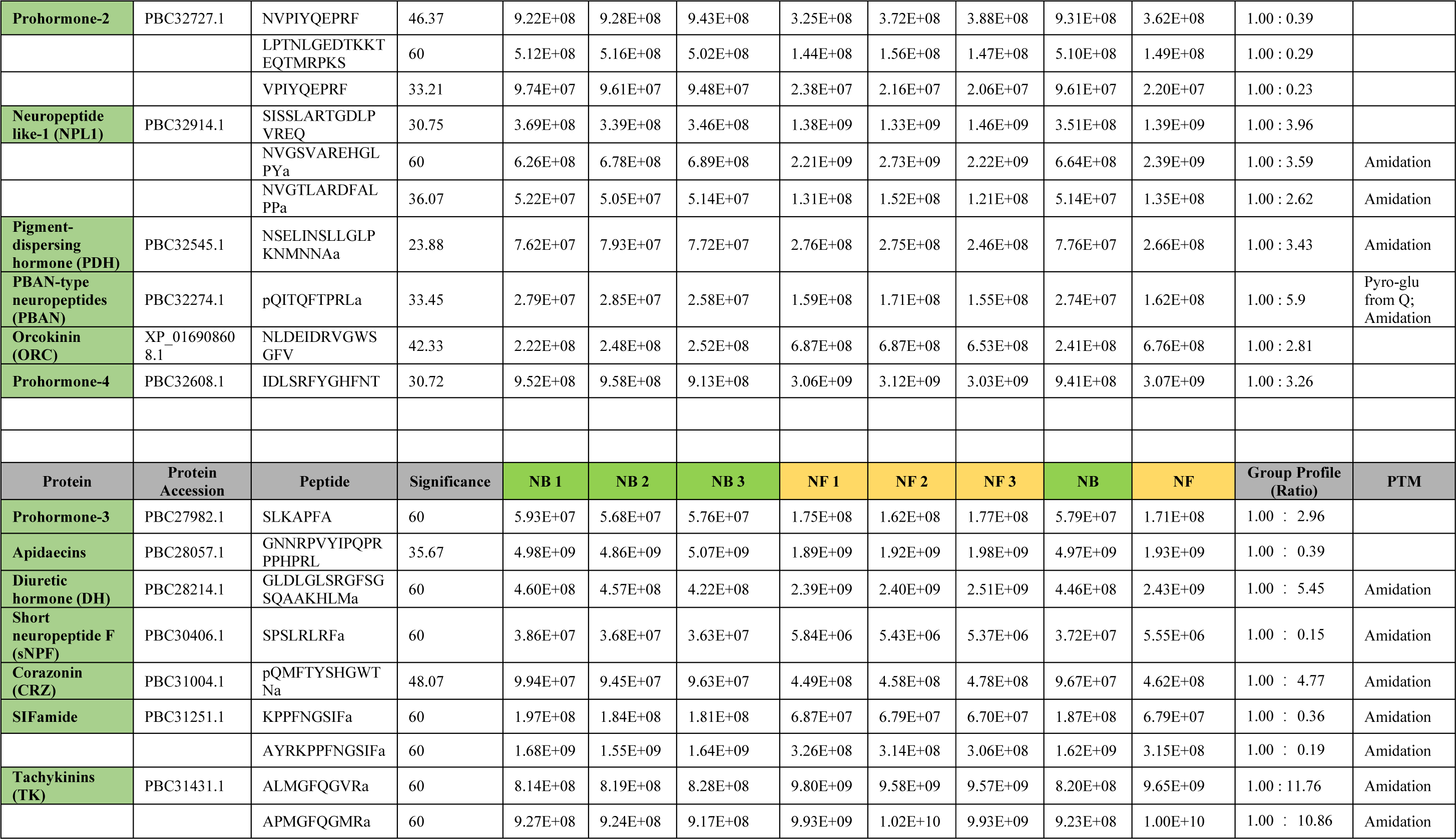
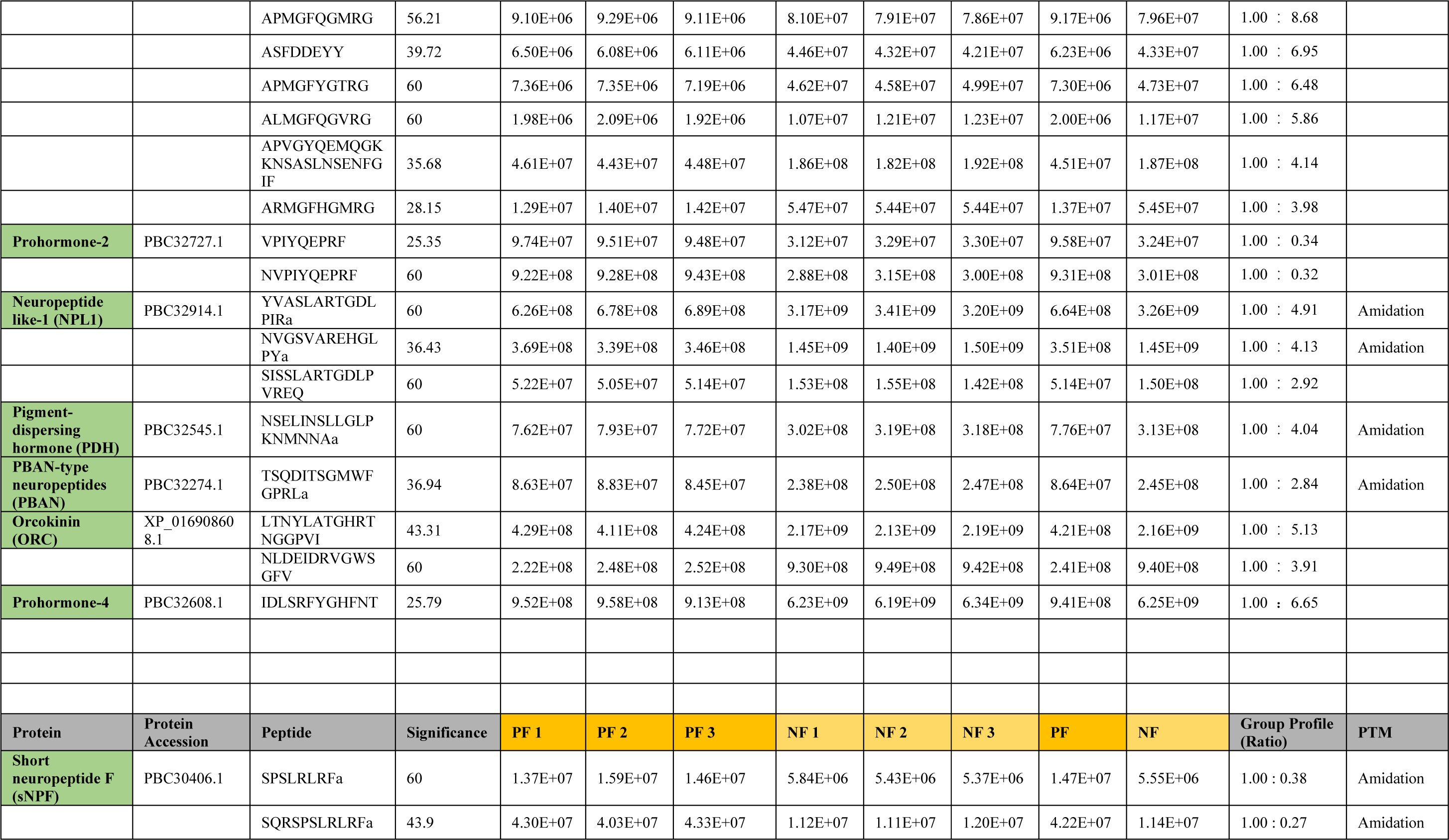
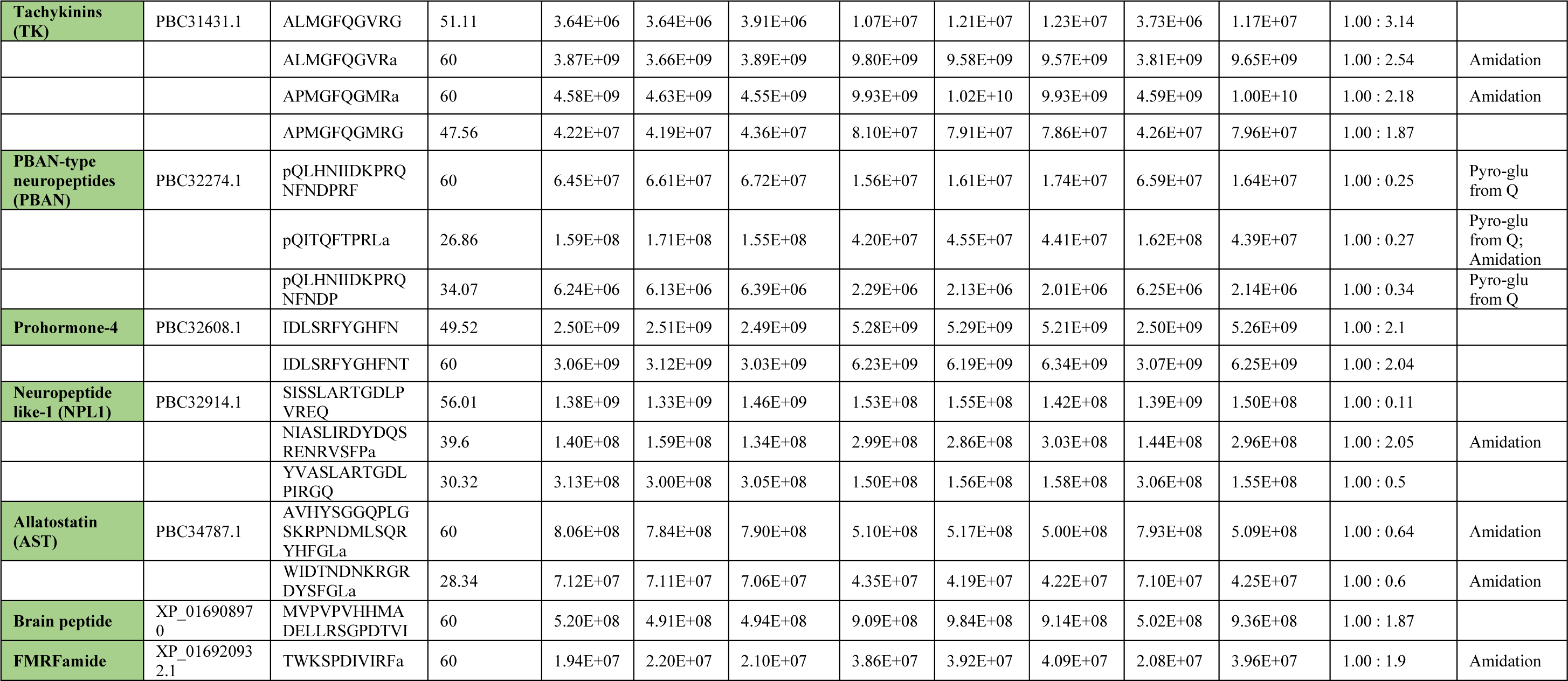
Quantitative neuropeptide comparison of different behavioral phenotypes of Apis cerana cerana workers. “Protein Accession” is the unique number given to mark the entry of a protein in the database NCBInr. “Peptide” is the amino acid sequence of the peptide. “Significance (-10lgP)” is the peptide confidence score. “NB” is nurse bee. “PF” is pollen forager. “NF” is nectar forager. “Group Profile (Ratio)” is the relative abundance ratio to the base group. “PTM” is post translational modification types present in the peptide.

**Table S7.**
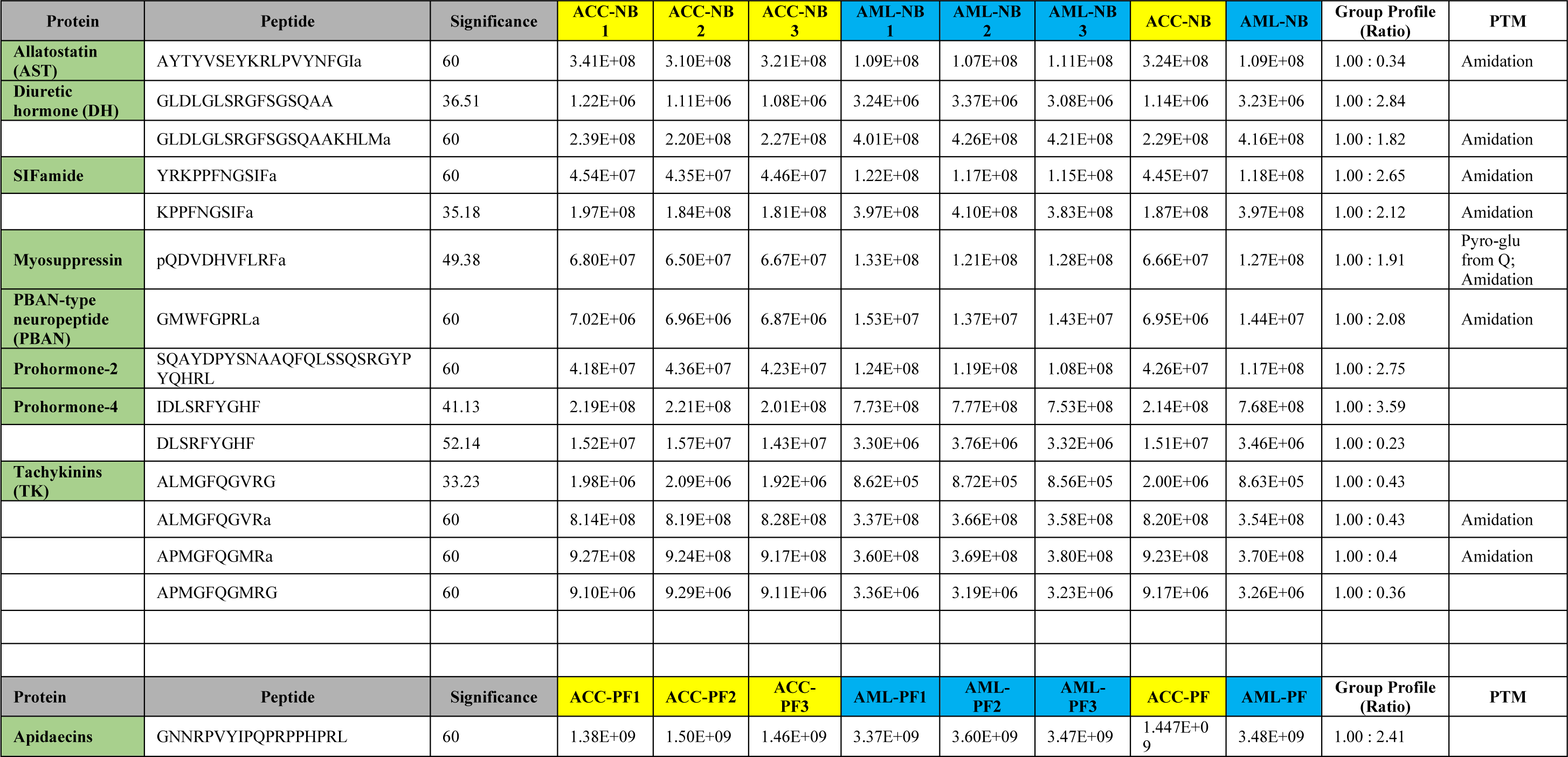
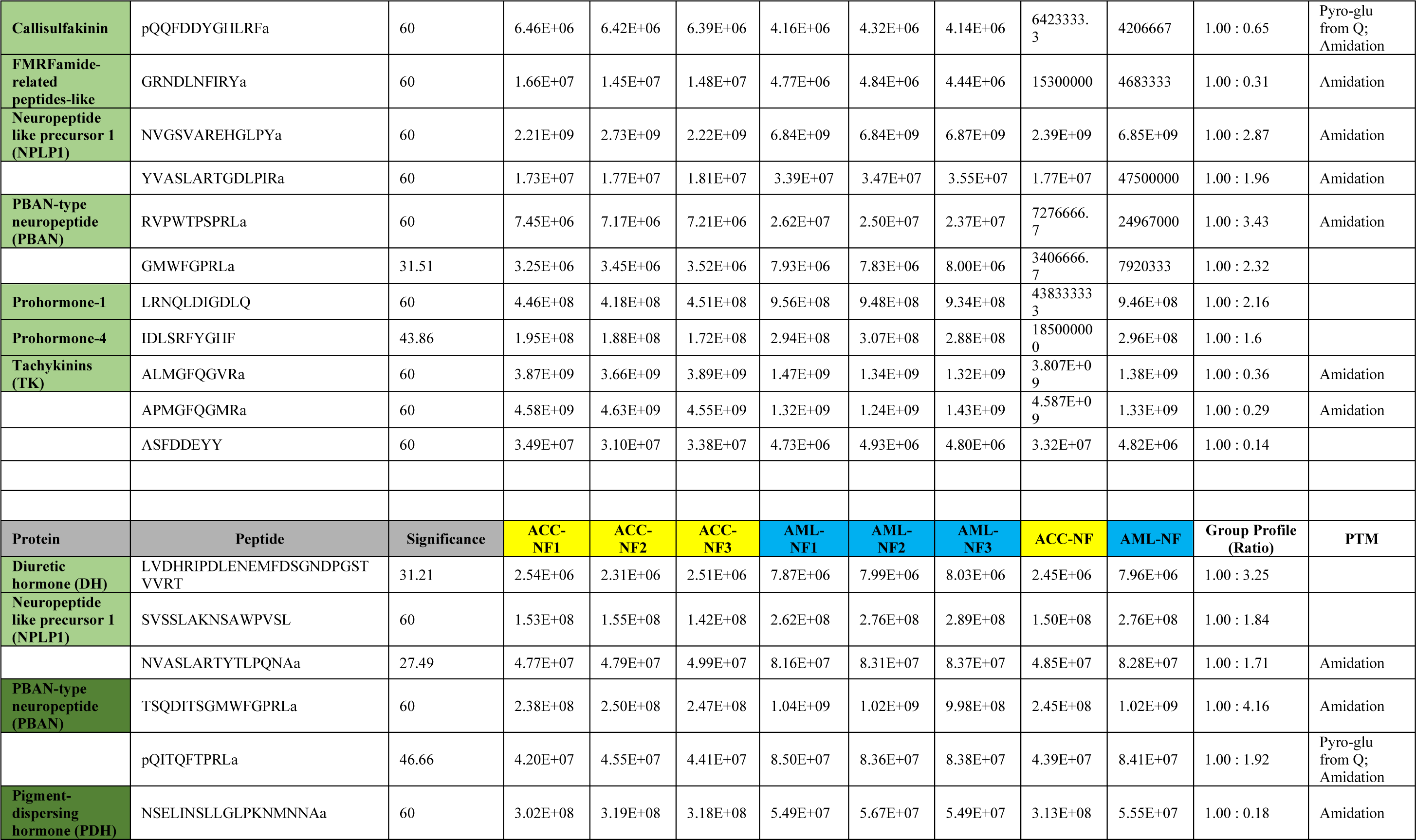
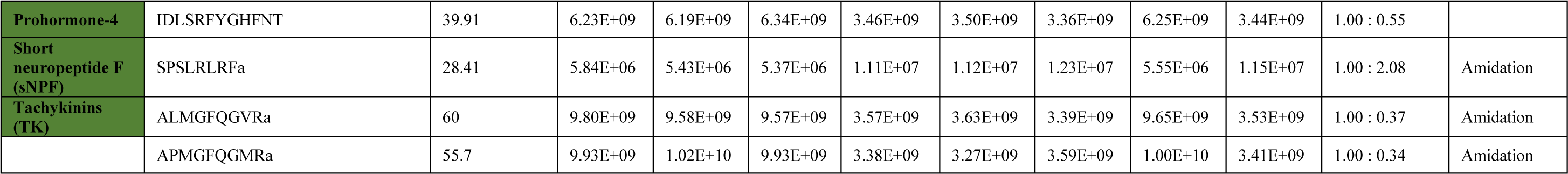
Quantitative neuropeptide comparison between *Apis cerana cerana* and *Apis mellifera ligustica*. “NB” is nurse bee. “PF” is pollen forager. “NF” is nectar forager. “Peptide” is the amino acid sequence of the peptide. “Significance (-10lgP)” is the peptide confidence score. “Group Profile (Ratio)” is the relative abundance ratio to the base group. “PTM” is post translational modification types present in the peptide.

**Table S8.** The proboscis extension response of workers after injection of ddH2O and TRP2.

| **Pollen foragers** | **ddH<sub>2</sub>O** |  |  |  | **TRP2** |  |  |
| --- | --- | --- | --- | --- | --- | --- | --- |
|  | **Concentration** | **Show PER** | **No PER** | **PER ratio** | **Show PER** | **No PER** | **PER ratio** |
|  | 0.1% | 19 | 36 | 34.55% | 9 | 44 | 16.98% |
|  | 0.3% | 21 | 34 | 38.18% | 11 | 42 | 20.75% |
|  | 1.0% | 30 | 25 | 54.55% | 12 | 41 | 22.64% |
|  | 3.0% | 36 | 19 | 65.45% | 15 | 38 | 28.30% |
|  | 10.0% | 38 | 17 | 69.09% | 17 | 36 | 32.08% |
|  | 30.0% | 48 | 7 | 87.27% | 25 | 28 | 47.17% |
|  | Pollen | 22 | 34 | 39.29% | 9 | 44 | 16.98% |
|  | Larva | 11 | 45 | 19.64% | 12 | 41 | 22.64% |

| **Nectar foragers** | **ddH<sub>2</sub>O** |  |  |  | **TRP2** |  |  |
| --- | --- | --- | --- | --- | --- | --- | --- |
|  | **Concentration** | **Show PER** | **No PER** | **PER ratio** | **Show PER** | **No PER** | **PER ratio** |
|  | 0.1% | 10 | 45 | 18.18% | 6 | 52 | 10.34% |
|  | 0.3% | 14 | 41 | 25.45% | 6 | 52 | 10.34% |
|  | 1.0% | 16 | 39 | 29.09% | 7 | 51 | 12.07% |
|  | 3.0% | 19 | 36 | 34.55% | 8 | 50 | 13.79% |
|  | 10.0% | 25 | 30 | 45.45% | 12 | 46 | 20.69% |
|  | 30.0% | 29 | 26 | 52.73% | 15 | 43 | 25.86% |
|  | Pollen | 7 | 46 | 13.21% | 8 | 44 | 15.38% |
|  | Larva | 9 | 44 | 16.98% | 6 | 46 | 11.54% |

| **Nurse bees** | **ddH<sub>2</sub>O** |  |  |  | **TRP2** |  |  |
| --- | --- | --- | --- | --- | --- | --- | --- |
|  | **Concentration** | **Show PER** | **No PER** | **PER ratio** | **Show PER** | **No PER** | **PER ratio** |
|  | 0.1% | 12 | 41 | 22.64% | 8 | 44 | 15.38% |
|  | 0.3% | 13 | 40 | 24.53% | 10 | 42 | 19.23% |
|  | 1.0% | 18 | 35 | 33.96% | 13 | 39 | 25.00% |
|  | 3.0% | 19 | 34 | 35.85% | 16 | 36 | 30.77% |
|  | 10.0% | 22 | 31 | 41.51% | 21 | 31 | 40.38% |
|  | 30.0% | 29 | 24 | 54.72% | 25 | 27 | 48.08% |
|  | Pollen | 5 | 50 | 9.09% | 7 | 48 | 12.73% |
|  | Larva | 21 | 32 | 39.62% | 10 | 45 | 18.18% |

**Table S9.**
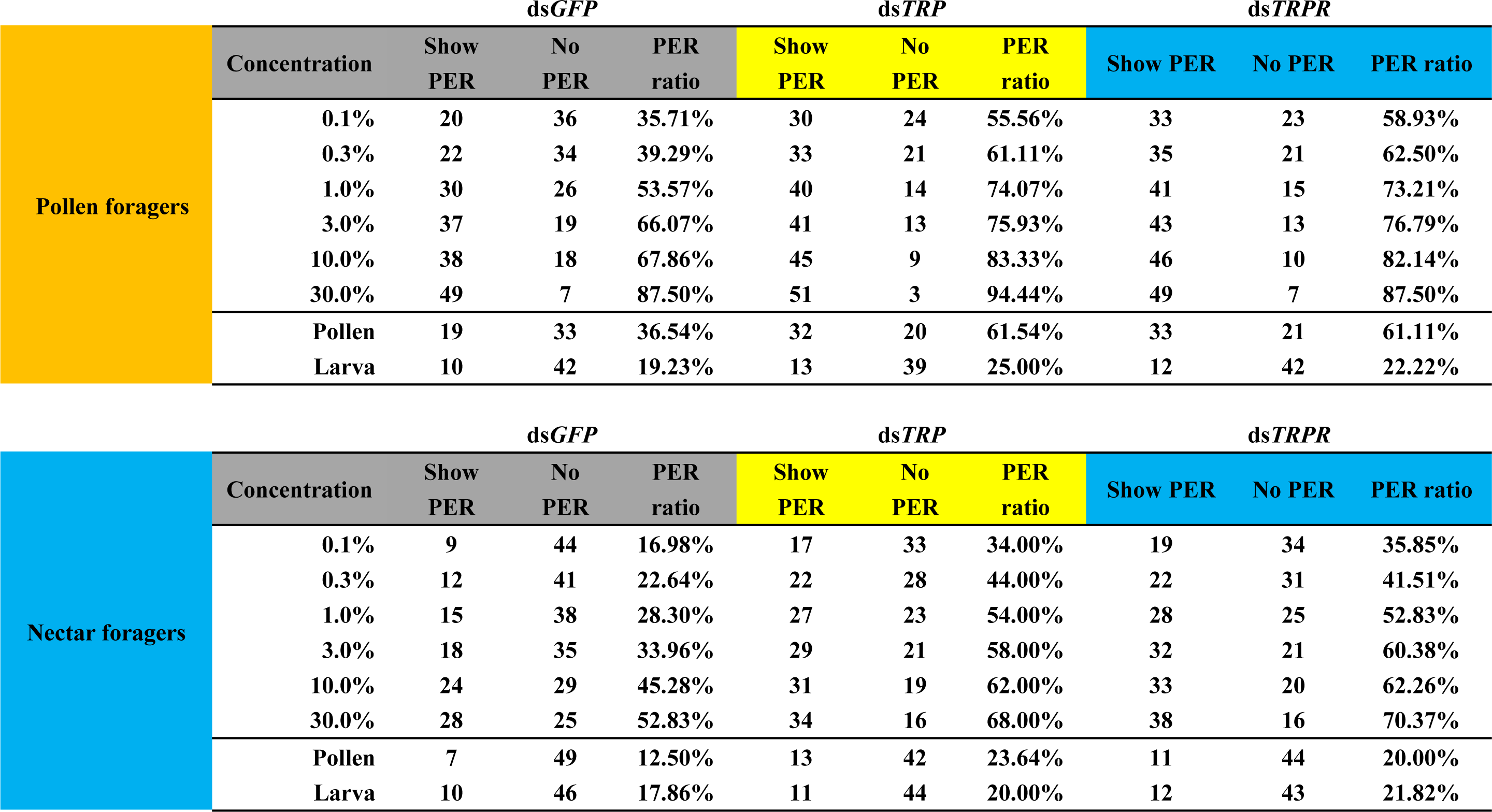
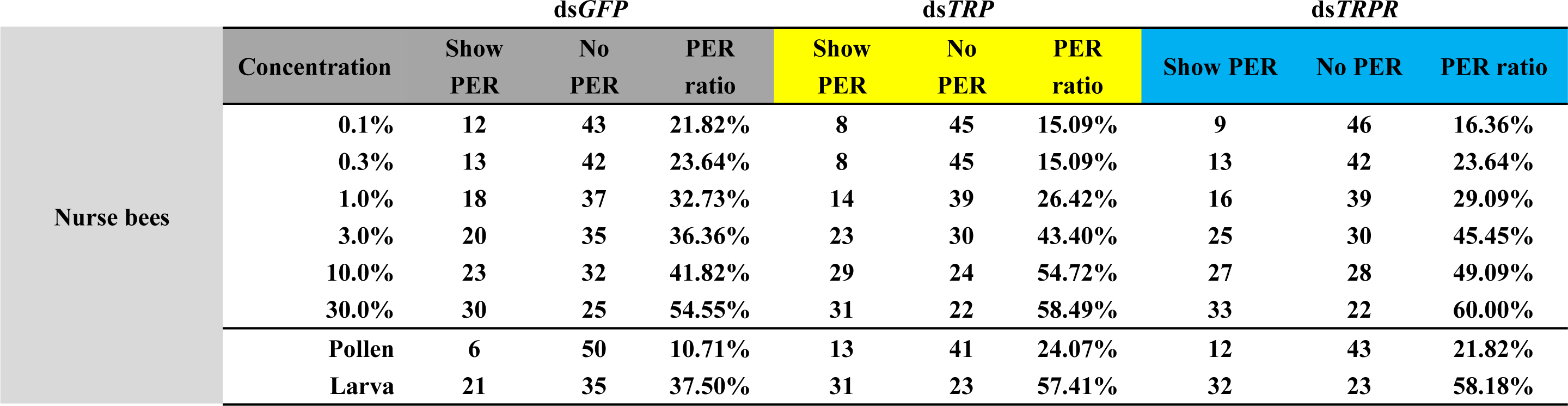
The proboscis extension response of workers after injection of ds*GFP*, ds*TRP*, and ds*TRPR*.

**Table S10.** Statistical differences in sucrose responsiveness after injection of ds*GFP*, ds*TRP*, and ds*TRPR*.

| Concentration | 0.10% | 0.30% | 1.00% | 3.00% | 10.00% | 30.00% |
| --- | --- | --- | --- | --- | --- | --- |
| **Pollen foragers** |  |  |  |  |  |  |
| *dsTRP* vs *dsGFP* | * | * | * | ns | ns | ns |
| *dsTRPR* vs *dsGFP* | * | * | * | ns | ns | ns |
| *dsTRP* vs *dsTRPR* | ns | ns | ns | ns | ns | ns |
| **Nectar foragers** |  |  |  |  |  |  |
| *dsTRP* vs *dsGFP* | * | * | ** | * | ns | ns |
| *dsTRPR* vs *dsGFP* | * | * | * | ** | ns | ns |
| *dsTRP* vs *dsTRPR* | ns | ns | ns | ns | ns | ns |
| **Nurse bees** |  |  |  |  |  |  |
| *dsTRP* vs *dsGFP* | ns | ns | ns | ns | ns | ns |
| *dsTRPR* vs *dsGFP* | ns | ns | ns | ns | ns | ns |
| *dsTRP* vs *dsTRPR* | ns | ns | ns | ns | ns | ns |

ns = $P > 0.05$ , \* $P < 0.05$ , \*\* $P < 0.01$

**Table S11.**
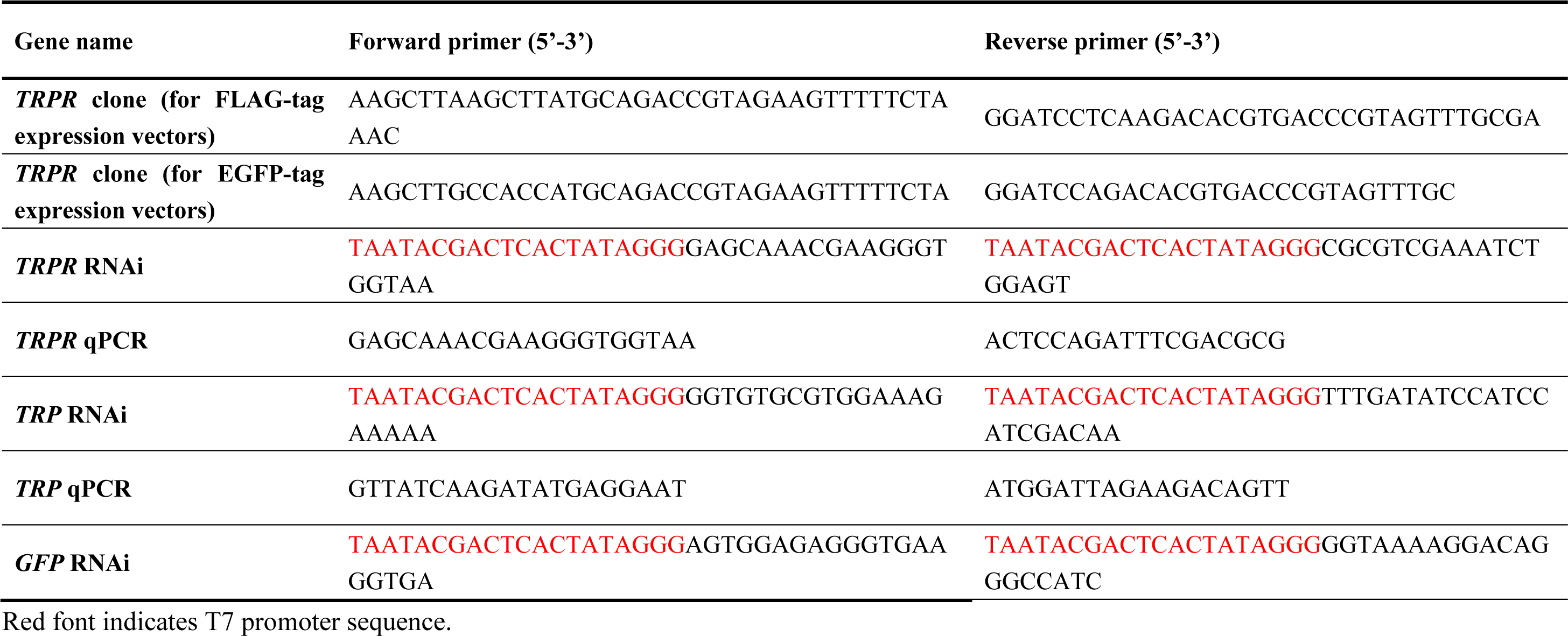
Sequence information of primers used in this study.

